# Dynamics of sex chromosome evolution in a rapid radiation of cichlid fishes

**DOI:** 10.1101/2020.10.23.335596

**Authors:** Athimed El Taher, Fabrizia Ronco, Michael Matschiner, Walter Salzburger, Astrid Böhne

## Abstract

Sex is a fundamental trait that is determined, depending on the species, by different environmental and/or genetic factors, including various types of sex chromosomes. While the functioning and emergence of sex chromosomes have been explored in species scattered across the eukaryotic tree of life, little is known about tempo and mode of sex chromosome evolution in closely related species. Here, we examine the dynamics of sex chromosome evolution in an archetypical example of adaptive radiation, the cichlid fishes of African Lake Tanganyika. Through inspection of male and female genomes from 244 cichlid taxa and the analysis of transcriptomes from 66 taxa, we identify signatures of sex chromosomes in 79 taxa, involving 12 different linkage groups. We estimate that Tanganyikan cichlids have the highest rates of sex chromosome turnover and heterogamety transitions known to date. We further show that the recruitment of chromosomes as sex chromosomes is not at random and that some chromosomes have convergently emerged as sex chromosomes in cichlids, which provides empirical support to the “limited options” hypothesis of sex chromosome evolution.

## Introduction

Sex chromosomes – referred to as Z and W in female and X and Y in male heterogametic sex determination (SD) systems – define, through their properties and combinations, the sex of an individual (*1*). The evolutionary trajectories of sex chromosomes differ from those of autosomes: Due to the restriction of one of the two sex chromosomes to one sex (W to females in ZW, Y to males in XY SD systems), their sex-specific inheritance (e.g., XY-fathers pass on their X exclusively to daughters and their Y to sons), and their reduced levels of recombination, sex chromosomes accumulate mutations more rapidly than autosomes, potentially leading to accelerated functional evolution (*2, 3*).

The functioning of a chromosome as sex chromosome is often short-lived on evolutionary time scales. This relative instability of sex chromosomes is due to turnovers (i.e., changes of the actual chromosome pair in use as sex chromosomes) caused by a new sex-determining mutation in a previously autosomal locus (*4*) or the translocation of the ancestral SD gene to another chromosome (e.g., (*5*)). Sex chromosome turnovers may be accompanied by a transition in heterogamety (*6*). Heterogamety can also change without transition in the chromosome pair that acts as sex chromosomes, which in this case likely involves a turnover of, or a mutation within, the actual SD locus (*7*).

The presumed major driving forces underlying turnovers of sex chromosomes are deleterious mutational load (*8, 9*), sexually antagonistic loci linked to a newly invading SD gene (*10, 11*), selection on restoring sex-ratios (*12*), and genetic drift (*6, 13, 14*). These drivers are predicted to differ in their respective outcome: turnovers induced by mutational load tend to preserve heterogamety (*8, 9*), while sexually antagonistic selection driven turnovers more readily induce a change of heterogamety (*10*).

Finally, the gene repertoire on previously existing sex chromosomes can also be extended by chromosomal fusion with an autosome, which then becomes sex-linked itself, leading to the formation of a neo-sex chromosome (*15*).

The frequency of occurrence of these different paths of sex chromosome evolution varies substantially across animal clades (*16*). For example, in some vertebrates (mammals and birds) the same sex chromosomes are shared across the entire class (*17*) (but see (*18*)). Models (*13*) as well as empirical observations (*15*) suggest that sex chromosomes such as those of mammals and (most) birds have differentiated to a degree that makes turnovers unlikely; these sex chromosomes are in an “evolutionary trap” (*19*). This is because a sex chromosome turnover requires the fixation of one of the previous sex chromosomes as an autosome, which becomes more deleterious and thus unlikely the more specialized and/or degenerated the sex chromosomes are (*20*). In other vertebrate lineages (amphibians, reptiles, and fish), frequent turnover events and continued recombination led to many different and mostly non-degenerated (homomorphic) sex chromosomes (*21, 22*).

As of to date, empirical studies on the dynamics of sex chromosome evolution are limited and scattered across different taxa. In an amphibian system with a rapid rate of sex chromosome turnover, the true frogs Ranidae, mutational load seems to be the major driving force of sex chromosome turnover (*22*). In geckos, a high rate of sex chromosome changes with heterogametic transitions potentially supports sexual antagonism as a key mechanism of these changes (*23*). However, an in-depth analysis of sex chromosome turnovers over short evolutionary timescales and with a broad taxon sampling is currently lacking (*16*).

Here, we examined sex chromosome evolution in an archetypical example of rapid organismal diversification, the adaptive radiation of cichlid fishes in African Lake Tanganyika (LT) (*24*). Teleost fishes are generally known for their species richness (*25*), but cichlids stand out in this clade on the basis of the “explosive” character of several of their adaptive radiations, giving rise to a total estimated number of over 3,000 species (*25*). Rapid speciation in adaptive radiations is usually attributed to ecological specialization and thus diversification in eco-morphological traits (*24*). Here, we were interested if the evolution of sex determination is keeping pace with other traits in cichlids by determining the diversity of SD systems and by investigating the dynamics of sex chromosome turnover across the entire LT cichlid radiation. The previously available data from about 30 African cichlid species (reviewed in (*26, 27*)) suggest that sex chromosomes are not conserved in this group with both, simple and polygenic SD systems being known from the different species investigated. An emerging picture is that certain chromosomes have recurrently been recruited as sex chromosomes in cichlids. However, available studies supporting the convergent recruitment of sex chromosomes have been based on cichlid species belonging to different lineages and the observed patterns have rarely been assessed in a phylogenetic framework, which makes inferences about rates of evolution as well as of convergence *versus* common ancestry difficult (but see (*26*)). Importantly, as of yet, no inclusive analysis of sex chromosome evolution exists for a cichlid adaptive radiation (nor for radiations in other fish families).

In this study, we inspected genomic (*24*) and transcriptomic (*28*) information from 229 Lake Tanganyika cichlid taxa as well as 18 cichlid species belonging to the Haplochromini and Lamprologini lineages phylogenetically nested within the LT radiation (*24, 29*) for signatures of sex chromosomes. Based on this nearly complete taxon sampling of the LT radiation and an available phylogenetic hypothesis based on genome-wide data (*24*), we estimated the amount and direction of sex chromosome turnovers in this young species flock. This allowed us to test for a possible contribution of sexual antagonism in the evolution of sex chromosomes in LT cichlids. Sexual antagonism has been suggested as a driving force of sex chromosome turnovers in sexually dimorphic cichlids of the Lake Malawi radiation (*30, 31*). However, unlike the cichlid adaptive radiation in Lake Malawi, which is composed solely of cichlids of the Haplochromini lineage, the endemic LT cichlid assemblage consists of 16 cichlid lineages (corresponding to the taxonomic assignment into tribes (*32*)), some of which are sexually dimorphic while others are not.

To assess the dynamics of sex chromosome turnover in fishes on a larger scale, we expanded our comparative analyses to other fish systems as well. In particular, we investigated sex chromosome turnovers in ricefishes (genus *Oryzias*), another model system for the evolution of sex chromosomes (*33*).

Sex differences in the recombination rate could contribute to the differentiation of sex chromosomes (*34*). Unlike in the extremely heterochiasmic frogs of the family Ranidae (*22*) and some fish model organisms (*35, 36*), recombination rates along chromosomes do not systematically nor drastically differ between the sexes in cichlids (*36, 37*). Although, in ricefishes, reduced rates of recombination have been linked to maleness in some species (*38*), this does not seem to be a general pattern in this group of fishes (*39*). Ricefishes and cichlids may hence have differing, probably lower, rates of sex chromosome degeneration than the heterochiasmic frogs, in which mutational load resulting from sex chromosome degeneration caused by suppressed recombination mainly drives turnover. In general, we expect fewer – if any – cichlid species to be in the “evolutionary trap” of degenerated sex chromosomes and more sex chromosome turnovers caused by sexual antagonism than by mutational load (*16*). Finally, with the identification of sex chromosomes in genetically very closely related species we pave the way for the subsequent characterization of sex-determining genes and/or the causal mutations leading to sex chromosome turnover.

## Results

### Sex chromosomes in LT cichlids

To identify sex chromosomes in LT cichlids, we screened male and female genomes of 244 taxa (*24*) as well as six transcriptomes of each of 66 taxa (*28*) for signatures of sex-linked regions, applying three complementary approaches: genome-wide association study (GWAS; on the genomic data, approach 1, see Materials and Methods), identification of sex-specific SNPs in the genomic data (approach 2), and tests of allele frequency differences on the transcriptome data (approach 3). Genomic locations of inferred sex-linked regions refer to linkage groups (LGs) of the used reference genome of a phylogenetically equidistant outgroup to the cichlid species of the LT radiation, the Nile tilapia (*Oreochromis niloticus*). To estimate sex chromosome turnover rates (see below), we used two different datasets; a “permissive dataset” including all sex chromosomes identified with approaches 1-3 and a “stringent dataset” excluding sex chromosomes that had support only in approach 2, i.e., lacking transcriptome data or support for small sex-linked and potentially non-expressed regions in the transcriptome data and occurring in tribes too small to be investigated with approach 1.

By combing the results of approaches 1-3, we detected signatures supportive of sex chromosomes in 78 endemic LT cichlid taxa as well as in the riverine Haplochromini *Orthochromis indermauri* (Fig. 1 and 2; tables S1-S3, figs. S1-S6).

**Fig. 1.**
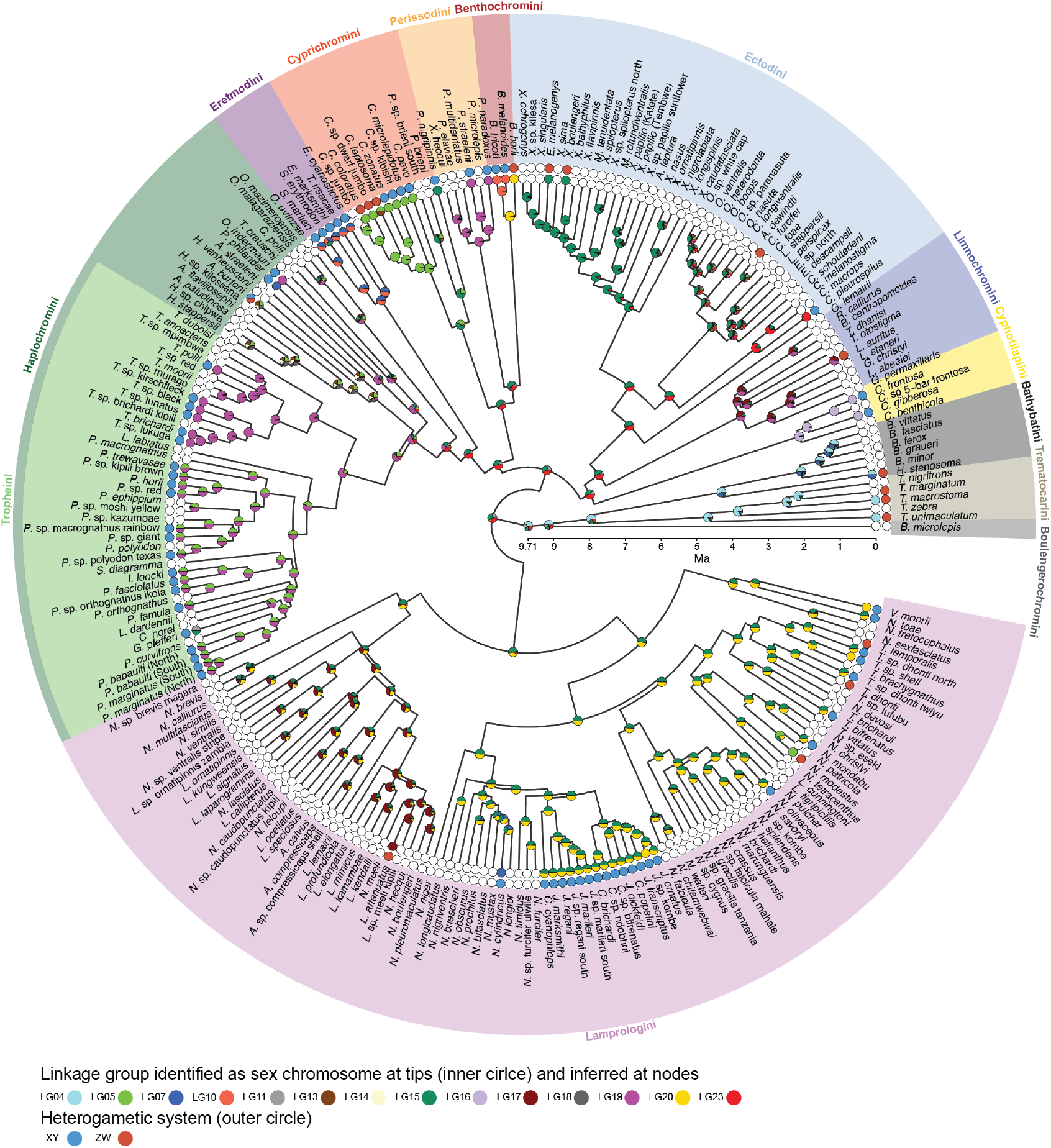
Sex chromosome evolution in the cichlid adaptive radiation in Lake Tanganyika. Sex chromosome state and ancestral state reconstruction in LT cichlids are placed on a time-calibrated species tree (*24*) with tribe-grouping indicated by color shading. The inner circle at tips shows identified sex-linked LGs. Colors refer to LGs of the reference genome; two- or more colored symbols at tips indicate sex chromosomal signals that were detected on two or more reference LGs suggesting chromosomal rearrangements. The outer circle indicates the heterogametic status of each species (blue: XY; red: ZW). White circles at tips indicate that no sex chromosome could be identified. Pie charts at nodes represent the probability for an LG being a sex chromosome at this time derived from ancestral state reconstructions.

**Fig. 2.**
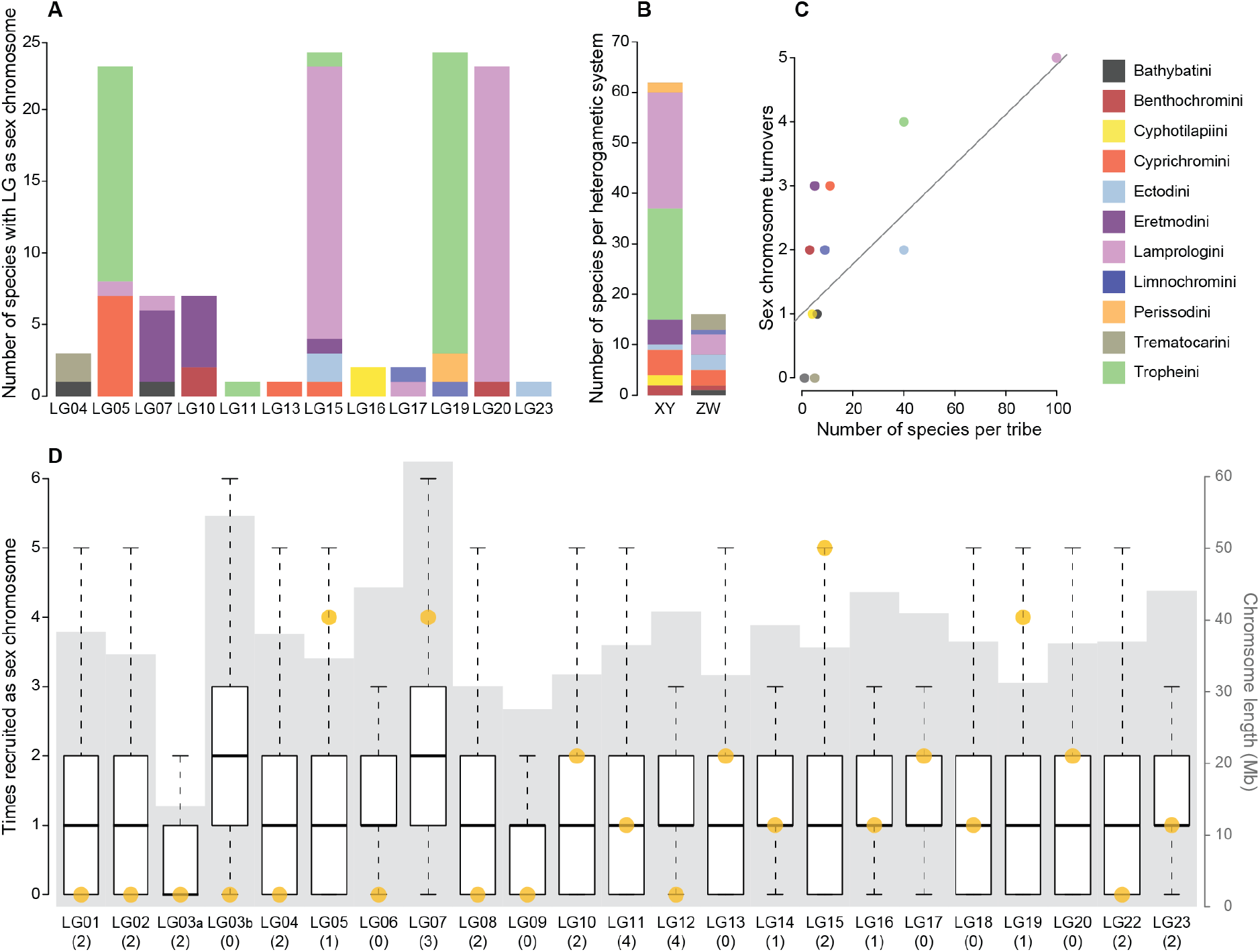
Non-random sex chromosome distribution in Lake Tanganyika cichlids. (**A**) Use of different LGs as sex chromosomes. Bars represent the number of times an LG has been detected as sex-linked at the species level and are colored according to tribe. (**B**) The occurrence of sex determination (SD) systems. Bars represent how often an XY or ZW SD system was identified at the species level and are colored according to tribe. (**C**) Association between species richness and sex chromosome turnover. The number of sex chromosome turnovers leading to the tips of each tribe is associated with the number of species investigated in each tribe (pGLS: *P*=0.0043, coeff=0.039). Dots are colored according to tribes; the line represents the linear model fitted to the data. (**D**) Boxplots showing the expected number of sex chromosome recruitments if recruitment was at random (10,000 permutations). Boxplot center lines represent the median, box limits the upper and lower quartiles, and whiskers the 1.5x interquartile range. Outliers are not shown. Ten reference LGs were never implicated in a turnover event in LT cichlids. Under random recruitment in the simulations this pattern occurred only in 2.01 % of all simulations. Yellow dots indicate the number of observed sex chromosome recruitments per LG derived from ancestral state reconstructions, gray background shading represents chromosome length in Mb and numbers below each boxplot indicate the number of previously described sex-determining genes on these LGs.

Approach 1 (GWAS), which was applied to the larger (that is, more species-rich) cichlid tribes from LT only, indeed revealed the presence of sex chromosomes shared among several species of their respective tribes. We thus identified an XY SD system on LG19 in Haplochromini/Tropheini (thereby confirming an XY system previously known from one species in this clade, *Tropheus* sp. “black” (*40*)), an XY and ZW system on LG05 in Cyprichromini (thereby confirming a ZW system previously described in *Cyprichromis leptosoma* (*40*)), and an XY system on LG15 and LG20 in Lamprologini. Surprisingly, with this approach, we did not detect a shared sex chromosome within the second-most species-rich cichlid tribe of LT, Ectodini.

Approach 2, the inspection of the genomes within tribes for an accumulation of sexspecific SNPs (i.e., XY or ZW SNPs) and outlier regions thereof revealed actually two different XY systems on LG19 within Haplochromini/Tropheini, one covering the first ~22 Mb of LG19 (in the genus *Tropheus* and in *O. indermauri*) and a second one located at the end of LG19 cooccurring with XY SNPs at the beginning of LG05 (in the second Tropheini clade grouping all genera but *Tropheus*, that is, 15 species belonging to the genera *Pseudosimochromis, Petrochromis*, and *Interochromis*, Fig. 1, fig. S4). We also recovered the narrow sex-linked region on LG20 detected with GWAS in Lamprologini, corroborating the effectiveness of this approach. As in approach 1, we did not detect a sex-differentiated region shared across species in Ectodini with approach 2.

When applied to the smaller tribes, approach 2 revealed rather narrow but clear outlier regions that were shared between subsets of species within the tribes Benthochromini (XY LG10, two species), Trematocarini (ZW LG04, two species) and Cyphotilapiini (XY LG16, two species) as well as in all members of the Eretmodini (XY, LG07 and LG10). We also detected a less pronounced and smaller ZW-outlier region on LG09 in the same two Cyphotilapiini species, a pattern potentially explained by variation in X-linked markers across the different species while simultaneously lacking homologous sites on the Y (hemizygosity in males). Due to this uncertainty and the stronger signal on LG16, we excluded the ZW-signal on LG09 of Cyphotilapiini in the subsequent analysis (in the permissive and stringent datasets). Within Bathybatini and Perissodini, we identified a chromosome-wide increase of ZW SNPs on LG07 and of XY SNPs on LG19, respectively, which, however, failed our thresholds for the permissive dataset (see Materials and Methods). Upon inspection of XY-ZW differences per species within these tribes (fig. S5), this pattern turned out to be caused by only one species in each tribe (*Hemibates stenosoma* and *Plecodus paradoxus*, respectively), which both showed signs of a differentiated sex chromosome across the entire length of the respective LG. Note that a ZW system on LG07 has previously been described in *H. stenosoma* (*40*) confirming the signal we detect. We could also confirm, with approach 3 (see below), the XY system spanning the full length of LG19 in *P. paradoxus* as well as in another Perissodini species, *Plecodus straeleni*, for which we did not have whole genome data of both sexes. We hence included the sex chromosomes of these species in all down-stream analyses.

Approach 3, the species-specific investigations of sex-specific SNPs based on replicate transcriptome data, confirmed all sex-differentiated regions shared among several species that spanned larger chromosomal regions (i.e., the two XY systems on LG19 and LG05/LG19 in Haplochromini/Tropheini, the XY and ZW systems on LG05 in Cyprichromini, the XY system in Eretmodini). With this approach, we also detected a ZW system on LG15 in two Ectodini species (*Xenotilapia boulengeri* and *Enantiopus melanogenys*). Approach 3 further permitted us to identify sex-linked LGs unique to eight additional species and not shared with their respective sister species. For example, we detected an XY system on LG23 in the Ectodini *Callochromis pleurospilus*, and a ZW system on LG20 in the Benthochromini *Benthochromis horii* (Fig. 1, table S2). In another four species, the RNA data showed a significant overrepresentation of either XY or ZW-SNP windows that, however, could not be attributed unambiguously on reference LGs (Fig. 1, table S2).

Overall, in nine of the 13 investigated tribes of the cichlid radiation in LT, several species shared the same SD system (chromosomal region and heterogametic type); however, we did not find a shared sex chromosome across members of different tribes.

We detected sex linkage on 12 out of the 23 reference LGs (Fig. 2). Eight of these reference LGs were sex-linked in species belonging to different tribes (Fig. 2A). Two reference LGs (LG14 and LG18) that we did not identify as sex chromosomes within any of the endemic LT cichlid radiation species, have respectively been identified as sex chromosomes in labstrains and one natural population of the haplochromine cichlid *Astatotilapia burtoni* (occurring in LT and affluent rivers) (*41, 42*). In addition to the published data for *A. burtoni*, we also included the previously published XY LG07 sex chromosome of *Pseudocrenilabrus philander* (Lake Chila) (*26*) in our subsequent analyses; this haplochromine species was included in the phylogenetic reconstruction used here (Fig. 1) but represented by only a single individual in the genomic dataset and hence not accessible to our three approaches.

In 62 of the LT cichlids (79.5% of the LT species with a sex chromosomal signal), the sex linkage was compatible with an XY system (Fig. 2B).

### Sex chromosome evolution in LT cichlids

Next, to determine when particular sex chromosomes emerged and to trace heterogamety transitions in the course of the cichlid adaptive radiation in LT, we performed ancestral state reconstructions along a time-calibrated species tree (*24*). We performed these analyses on the permissive as well as on the stringent dataset.

We reconstructed 30 sex chromosome turnovers in the radiation and LG04 as the likely sex chromosome at its root (permissive dataset; 27 turnover events with the stringent dataset), translating into an estimated rate of 0.186 turnovers per Myr (Fig. 1, fig. S7, permissive dataset; turnover rate with the stringent dataset was 0.187 turnovers per Myr). On average, we therefore expect one sex chromosome turnover event between two species that diverged ~2.7 Ma. This rate estimate was ten times higher than the one that we calculated for ricefishes (Adrianichthyidae; 0.02 transitions per Myr; fig. S8 and table S4; 19 species investigated, see Materials and Methods).

The distribution of sex chromosomes in LT cichlids differed from random expectations (Fig. 2D). There was no association between the size of a reference LG, the number of genes on a reference LG, or the number of known SD candidate genes on a reference LG and the frequency at which these LGs became a sex chromosome in LT cichlids (Fig. 2D). Our findings thus corroborate that SD is a rapidly and non-randomly evolving trait in cichlids. We further found that the number of turnovers in a tribe is associated with its species richness (Fig. 2C, pGLS: *P*=0.0043, coeff=0.039), suggesting that the turnover rate has been relatively constant throughout the radiation.

Our heterogamety reconstructions further suggested that XY is the most likely ancestral state in the cichlid adaptive radiation in LT (fig. S9). Subsequently, 11 transitions occurred from XY to ZW (permissive dataset; 11 towards ZW and one towards XY with the stringent dataset). Heterogamety transitions are predicted to have a directional bias towards new dominant sex chromosomes (*13*), suggesting that in cichlids from LT – just like in cichlids from Lake Malawi (*30, 31*) – new W chromosomes are dominant over ancestral Ys.

When integrating the reconstructed transitions in heterogamety and sex chromosomes, we found heterogamety changes that were uncoupled from turnovers in LGs and that were hence not captured in our rate estimate of sex chromosome turnover: A transitions from XY to ZW was detected on LG05 in Cyprichromini and on LG04 in Trematocarini and in Bathybatini (*H. stenosoma*) (Fig. 1; figs. S7 and S9).

The overlap of heterogametic and sex chromosome turnovers also showed that the majority (23 *versus* seven) of the observed sex chromosome turnovers in LT cichlids preserved the heterogametic state, suggesting that mutational load, predicted to keep heterogametic state (*22*), might be a major driver of sex chromosome turnover in cichlids as well. The transitions with a change in heterogamety offer the possibility to investigate the actual potential of sexual antagonistic selection between very young species (the divergence time between e.g., *Paracyprichromis* and *Cyprichromis*, between which a turnover has occurred, is ~3.8 Ma). The heterogametic status of the four species for which we could not identify the sex-linked LG (see above) led to additional heterogamety transitions that were not reflected in the sex chromosome turnover rate.

Overall, the heterogamety transition rate in LT cichlids (0.028 transitions per Myr with the permissive dataset; 0.031 per Myr with the stringent dataset) was about four times higher than in ricefishes (0.007 transitions per Myr; ancestral state: ZW). To explore heterogamety changes on a greater taxonomic scale, we also calculated heterogamety transition rates for all ray-finned fishes included in both the Tree of Sex database (http://www.treeofsex.org/) and a recent comprehensive phylogeny (*43*) (543 species analyzed in total). Our analysis estimated a rate of 0.009 transitions per Myr for ray-finned fishes as a whole and identified XY as the ancestral state (table S5; fig. S8). Across the ray-finned fish phylogeny, transitions from XY to ZW were significantly younger than those from ZW to XY (fig. S8B, *P*=0.01428).

### Chromosome fusions and novel sex chromosomes

Novel sex chromosomes can be created by chromosome fusions (*44*), which can contribute to reproductive isolation and eventually drive speciation over mis-segregation at meiosis, changes in recombination rates, novel physical combinations of loci, and changes in gene expression (*45–47*). The here identified signatures of sex-linkage suggest that several sex-chromosome/autosome fusions have occurred in the course of the cichlid radiation in LT or that autosome/autosome fusions occurred prior to the recruitment of the then fused autosomes as sex chromosome (Fig. 1). The distribution of sex-differentiated genomic regions indicated a fusion (or large chromosomal translocations) between LG05 and LG19 in Haplochromini/Tropheini and between LG15 and LG20 in Lamprologini (Fig. 1, fig. S1). There was also some support for the previously described genome rearrangements in the tribe Eretmodini (*48*), which showed an increase of XY SNPs on several LGs (Fig. 1, fig. S3). Additional sex-differentiated regions point to species-specific fusion events (e.g., LG11 and LG15 in *Gnathochromis pfefferi*). Our analyses also confirmed the reported sex linkage of LG04 as well as of LG07 in *H. stenosoma (40*) (fig. S5). Chromosome fusions have previously been implicated with the evolution of novel sex chromosomes in other taxa, as well as in the haplochromine cichlid *A. burtoni* (*41, 42*).

Interestingly, the so far only karyotypically investigated member of the tribe Tropheini, *Ctenochromis horei*, has a reduced number of chromosomes in a male and an unsexed individual (2n=40) compared to other Haplochromini, which usually feature 2n=42 (*48*). We did not detect the LG05/LG19 XY system found in many other Tropheini in *C. horei*. Hence, while the karyotype of this species indeed supports chromosomal fusions in the Haplochromini/Tropheini, this data cannot help to resolve when and how these events occurred. The data at hand are sparse but it might be that several large chromosomal rearrangements occurred before the novel chromosomes were recruited as sex chromosomes, asking for further investigations of the driving forces of these fusions.

### Convergent evolution of sex chromosomes

On some LGs, the regions that showed sex linkage were largely the same between members of different tribes (fig. S10), which can either be explained by common ancestry or by the independent (convergent) recruitment of those LGs as sex chromosome. In particular on LG19, several closely related species including six *Tropheus* species (Haplochromini/Tropheini), the riverine haplochromine *O. indermauri*, and the Perissodini *P. paradoxus* and *P. straeleni* feature an XY system in the same chromosomal region (fig. S10). Our ancestral state reconstruction suggested an independent origin of the LG19 SD system in Perissodini and *Tropheus*, in each case early in their tribe’s evolutionary history, and another independent origin in the terminal branch leading to *O. indermauri* (fig. S7). Phylogenetic inference from Y- and X-haplotypes indeed supported the independent evolution of LG19 as XY sex chromosome in Perissodini (Fig. 3), while grouping together the Y-haplotypes of the *Tropheus* species and *O. indermauri*. This suggests common ancestry of the XY system in the two haplochromine clades with an origin either early on in haplochromines (implying several losses later in the evolution of this tribe; likely because of this, such a scenario was not supported by ancestral state reconstruction) or a later origin and inheritance of the sex chromosomal system in *Tropheus* and *O. indermauri* from an extinct or unsampled taxon.

**Fig. 3.**
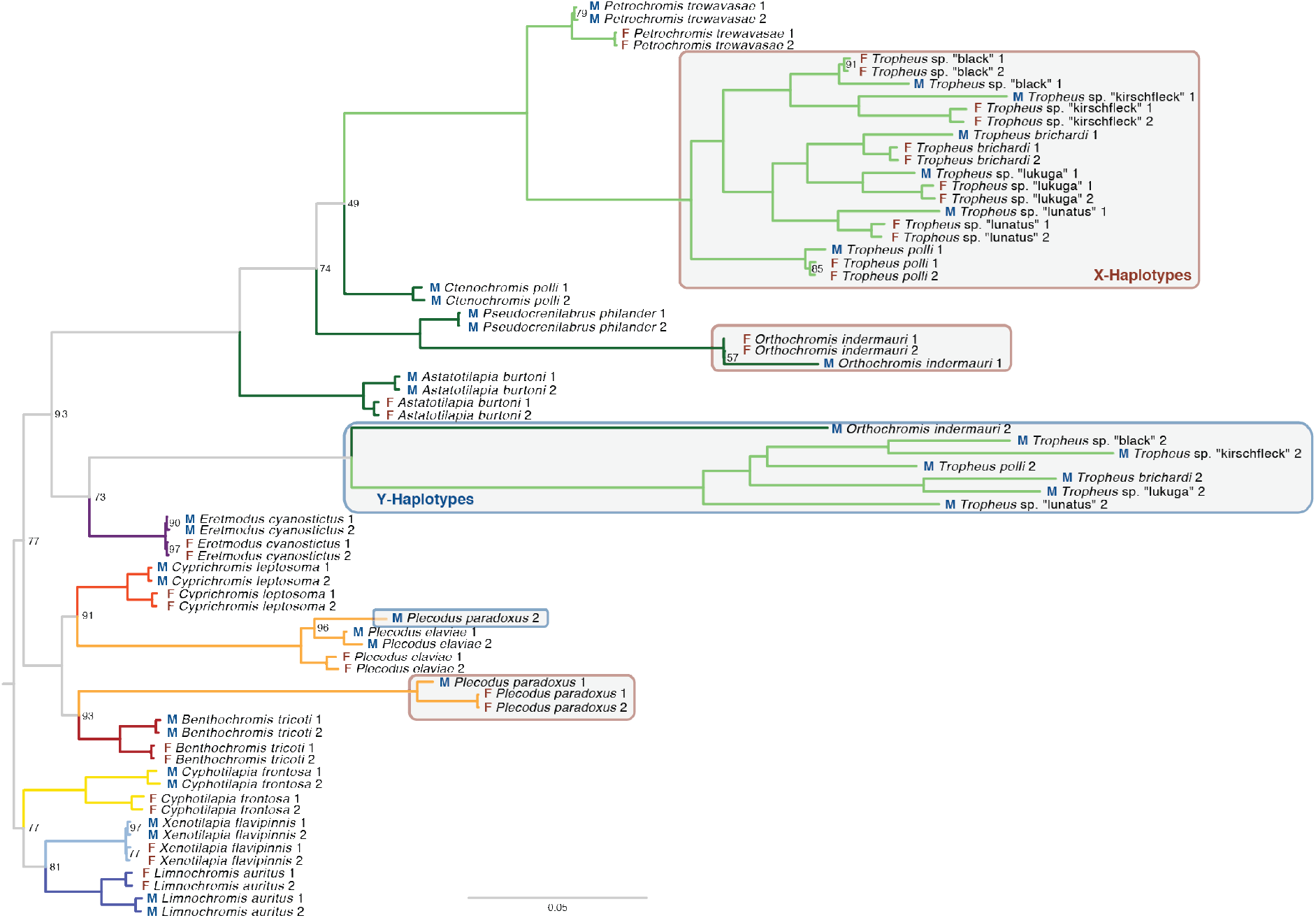
Convergent evolution of LG19 as XY sex chromosome in two Tanganyikan cichlid tribes. Phylogenetic tree of X- and Y-haplotype sequences does not group the *P. paradoxus* Y-haplotype with the *Tropheus* Y-haplotypes but supports the species tree, suggesting convergent evolution. The Y-haplotype of the non-LT riverine haplochromine *O. indermauri* groups with the Y-haplotypes of the *Tropheus* species, supporting monophyly of this sex chromosomal system. The scale bar indicates the number of substitutions per site; values at nodes represent bootstrap support (% of 1,000 bootstraps, if no value is shown the node support was 100%).

The XY sex chromosome system on LG05/LG19 found in the second clade of Haplochromini/Tropheini (grouping all genera except *Tropheus*) must be derived from another independent evolutionary event, since the regions on LG19 that show XY alleles in the two Haplochromini/Tropheini clades are not overlapping (fig. S10) and also do not group together in the phylogenetic tree of LG19 haplotypes (Fig. 3). Other convergent cases of sex chromosome recruitment supported by our ancestral state reconstruction involved LG05 (in Cyprichromini and the haplochromine *A. burtoni* (*41, 42*)) and LG07. LG07 has independently been recruited as a sex chromosome in *H. stenosoma* (Bathybatini) (*40*), in Eretmodini, in the lamprologine *Neolamprologus cylindricus* (Fig. 1, fig. S7), in several Lake Malawi cichlids (Haplochromini) (*30, 31*), as well as in *P. philander* (Haplochromini) (*26*), making it the most widespread sex-linked LG known in cichlids to date.

### Sex chromosome differentiation

A comparison of the proportion of sex-specific sites on the different sex-linked LGs revealed a continuum of sex chromosome differentiation in the cichlid adaptive radiation in LT (Fig. 4, fig. S10), ranging from a few kb (LG20 in Lamprologini) to almost full chromosomal length (LG05 in Cyprichromini, LG19 in *Tropheus* and Perissodini). Varying lengths of sex-differentiated regions were even detected within the same LG when being used as sex chromosome by different lineages (e.g., the sex-differentiated region on LG05 spans only 8 Mb in Tropheini but the entire LG in Cyprichromini).

**Fig. 4.**
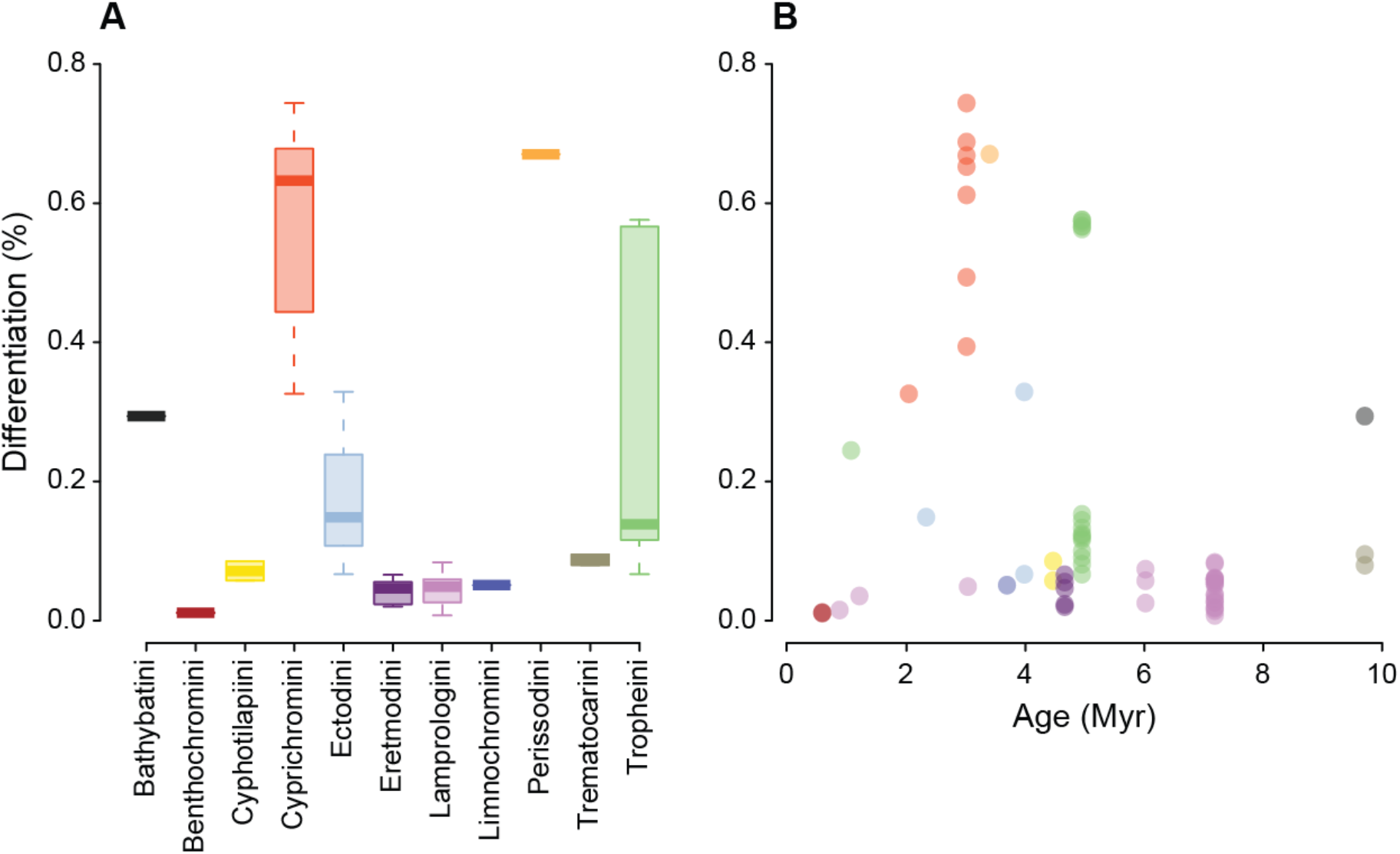
Sex chromosome differentiation in Lake Tanganyika cichlids. (**A**) Size distribution of sex-differentiated regions. The size of these regions corresponds to the proportion of the LG with windows that have more sex-specific SNPs than two times the mean across all windows. (**B**) Per-species proportion of the chromosome(s) showing sex differentiation and corresponding estimated ages of the sex chromosomal system based on ancestral state reconstructions on a time-calibrated species tree. The degree of differentiation is not associated with the estimated age of origin (pGLS: *P*=0.9049, coeff=0.0011).

The canonical model of sex chromosome evolution predicts progressing differentiation of sex chromosomes with time (*2*). Contrastingly, we found no association between the estimated age of origin of a sex chromosome and its degree of differentiation (Fig. 4, pGLS: *P*=0.9049, coeff=0.0011). Some very young sex chromosomes showed signs of differentiation, i.e., sex-specific sites, along almost the full length of an LG, suggesting widespread suppression of recombination along these sex chromosomes.

### Candidate genes of sex determination in LT cichlids

Our inspection of known genes implicated in SD revealed that such genes were located on all LGs, including those for which no sex linkage was detected, with no particular overrepresentation on certain LGs (fig. S11). The regions with the strongest signal for being sex-differentiated did not contain any of these genes (table S2). However, through the inspections of the regions with the strongest signs of sex linkage we identified promising new candidate genes for SD in these regions, such as *tox2* in Lamprologini, an HMG-box transcription factor involved in the hypothalamic-pituitary-gonadal system. *Tox2* resembles the mammalian master SD gene *Sry* (*49*), which also codes for an HMG-box protein.

In cichlids from Lake Malawi and Lake Victoria (*30, 50*), sexually antagonistic color genes underlying a characteristic orange-blotched color pattern are linked to SD genes, creating the potential for speciation by sexual selection. In LT cichlids, which in general do not feature the orange-blotched phenotypes, we did not find any obvious pattern in the localization of color genes on sex-linked LGs (fig. S11).

## Discussion

Here we report the identification of genomic signatures supportive of sex chromosomes in 79 taxa of cichlid fishes, most of which belonging to the cichlid adaptive radiation of LT, based on the analysis of whole-genome data from virtually all cichlid species of the radiation (*24*) and transcriptome data from a representative set of 66 taxa (*28*).

Models (*13*) and empirical observations (*19*) suggest that, beyond a certain degree of differentiation, sex chromosome turnover becomes unlikely. On the other hand, frequent turnovers, sex reversal, and continued recombination can contribute to counteract sex chromosome differentiation (*9, 51*). Our analyses revealed that, in the cichlid adaptive radiation of LT, sex chromosome turnovers seem to have occurred very frequently (Fig. 1), indicating that the cichlids’ sex chromosomes have not (yet) reached a threshold preventing turnover, but that their sex chromosomes remain dynamic instead.

Sex chromosome recruitment in LT cichlids is non-random with respect to the recruited chromosome (Fig. 2). This pattern becomes even more apparent when the LT cichlids are compared to other African cichlid species (Fig. 5), revealing that some LGs (in particular LG05, LG07, and LG19) emerged multiple times as sex chromosomes whereas others never appeared as such. This corroborates the hypothesis that particular chromosomes are preferentially (*52*) or even cyclically (*9, 51*) recruited as sex chromosomes. Within LT cichlids, sex chromosome turnovers have likely been driven by a combination of mutational load and sexual antagonism. However, we detected a prevailing persistence of male heterogamety in LT cichlids, which is a common pattern in fishes (*53*), suggesting a smaller role for sexual antagonism than previously postulated. Furthermore, the observed prevalence of XY systems is compatible with models of speciation driven by sexual selection and sex-ratio distortion in cichlids that predict higher probabilities for the maintenance of male heterogamety (*54*).

**Fig. 5.**
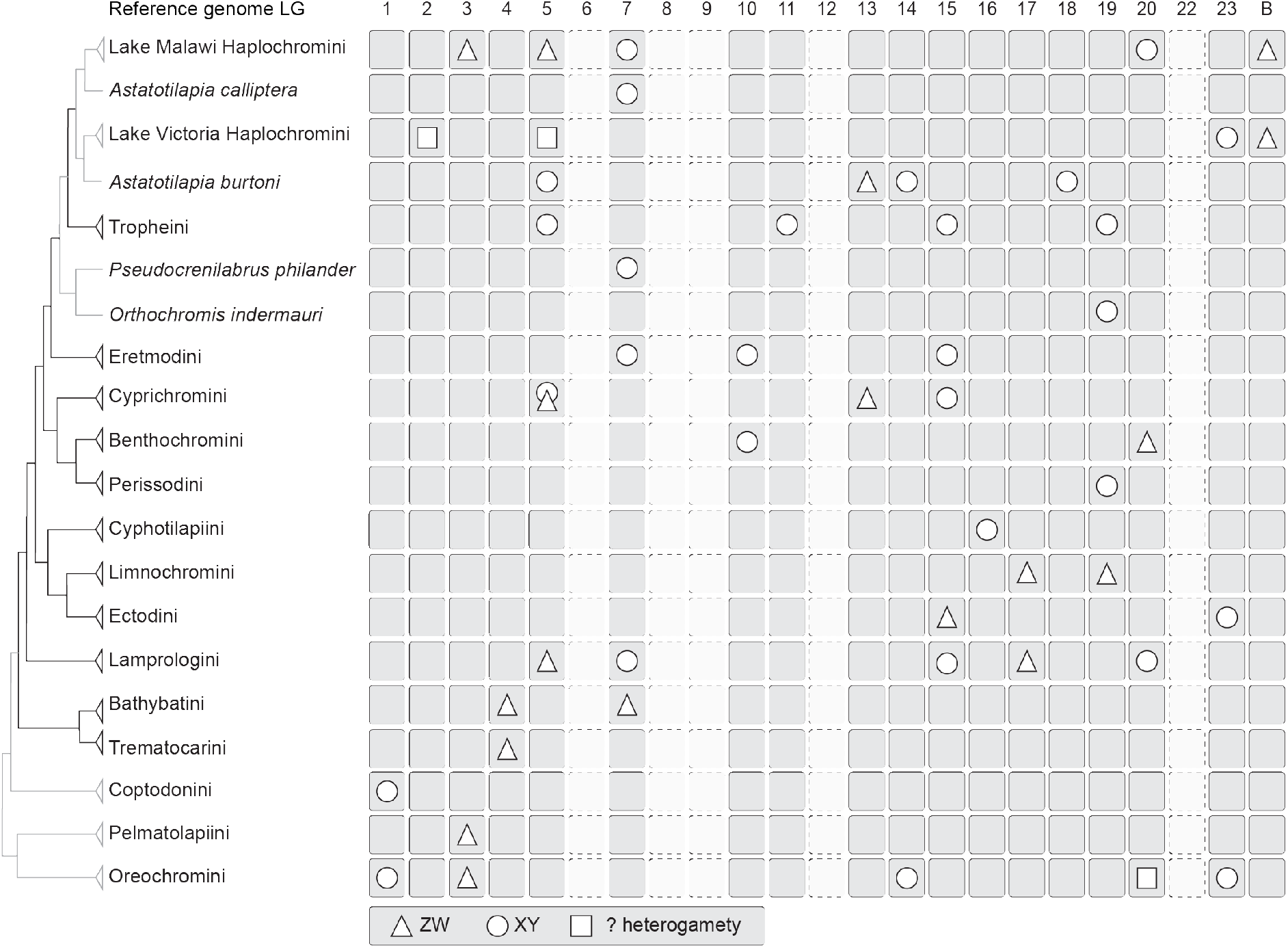
Sex chromosome evolution in African cichlids. Phylogenetic relationships (*24, 26*) and sex chromosome occurrence with reference to the genome of the Nile tilapia (*O. niloticus*) in African cichlids. Cichlid lineages found in Lake Tanganyika are indicated with black branches, cichlids from other lakes or rivers with gray branches. Sex chromosome information is derived from this study, and previously published summaries (*26, 27*).

The evolution of a novel sex determiner driven by linkage to a sexually antagonistic color locus has previously been documented in haplochromine cichlids from Lake Malawi (*30*), which are characterized by pronounced levels of sexual dimorphism. In our set of mostly riverine Haplochromini and Tropheini species (the LT representatives of this clade), we found that a single sex chromosome system prevails, XY on LG05/LG19, which was probably established after a turnover from the rather strongly differentiated XY LG19 system present in the genus *Tropheus*. It thus appears that in the Tropheini, in which sexual dimorphism is much less pronounced (and even absent in some species) compared to the radiations of Haplochromini in lakes Malawi and Victoria, sexual antagonism does not play a prominent role as a driving force for sex chromosome turnover. Still, several Tropheini species seem to have lost the XY LG05/LG19 SD system and we were mostly unable to detect a new system that replaced it based on the available transcriptome data, probably because the sex-linked chromosomal regions are rather small. These species will be particularly interesting to investigate further for the presence and the drivers of very young, novel sex chromosomes with a potential role for sexual antagonism impacting sex chromosome turnover (*55*). In addition, the observed cases of young homologous sex chromosome turnovers between closely related species (e.g., in the genus *Cyprichromis* on LG05 or in Trematocarini on LG04), which are indeed compatible with a role for sexual antagonism as driving force in cichlid sex chromosome evolution (*10*), open the route for further analysis of the causal mutations driving sex chromosome turnovers. Especially the presence of several ZW as well as XY species in Cyprichromini, potentially caused by a single transition event on the same chromosome, will allow in the future to trace which alleles have been affected by a heterogamety change. Such analyses may eventually reveal the causal mutation(s) (supposedly within the SD gene) of the heterogamety turnover and the dominance relationships between XY and ZW systems.

We failed to detect signatures of sex linkage in several of the LT cichlids, which certainly can, to some extent, be explained by our limited sample size per species, the lack of sex chromosomes shared between several species in some tribes/genera, the lack of strongly differentiated sex chromosomes, and/or the limited power to detect small sex-specific regions, especially when using transcriptome data, as well as complex polygenic SD systems. While the limited sample size per-species in the currently available data may have left some SD regions undetected, we found it particularly intriguing that we could identify sex-linked regions only in three species across the second-most species-rich tribe, Ectodini (the available transcriptome data are representative in terms of species-richness per tribe (*28*)). Some species of this tribe display an impressive level of sexual dimorphism, suggesting similar or even more pronounced sexual antagonistic selection compared to tribes such as the Haplochromini/Tropheini, which show relatively strongly differentiated sex chromosomes. It will thus be interesting to examine if Ectodini (and also members of other LT cichlid tribes) have species-specific very small, if any sex-linked genome regions that our approaches failed to detect. This could further reveal if selective forces on SD differ within the radiation and if our assessment of the sex chromosome turnover rate is underestimating the true dynamics of sex chromosome change in LT cichlids.

The sex chromosomal status of many species clearly remains to be identified in LT cichlids. Still, our ancestral state reconstructions estimated a sex chromosome turnover rate in LT cichlids that is ten times higher than the one in ricefishes, another group of fishes with an astonishing diversity of sex chromosomes, as well as the one published for true frogs, which was previously considered the fastest sex chromosome turnover rate known in vertebrates (*22*). Note that extremely high numbers of SD system turnovers have also been described in geckos (*23*), but these have so far not been used to calculate a comparable rate estimate.

Chromosome fusions could drive speciation through incompatibilities in genome structure (*45–47*) and cytogenetic analyses have indeed provided evidence for chromosome fusion and fissions in some cichlid species (*48*); however, their impact on cichlid diversification has not yet been assessed. Sex-chromosome/autosome fusions generating an odd number of chromosomes in one sex and leading to the formation of neo-sex chromosomes can be driven by altering expression of genes on the translocated chromosome (*56, 57*), sexually antagonistic selection resolving conflict by restricting an antagonistic allele to a sex chromosome (*58*), or meiotic drive (*59*). Until now, differences in chromosome number between male and female cichlids have not been reported, with the notable exception of copy number variations in female-determining B chromosomes in Lake Victoria and Lake Malawi cichlids (*60, 61*). For the limited number of cytogenetically investigated LT cichlid species, males and females have the same number of regular chromosomes, and across cichlids in general, chromosome numbers differ little (*48*). Overall, our analyses provide support for several large chromosomal rearrangements between the identified sex-linked LGs, suggesting that structural changes in the genome and the emergence of sex chromosomes are coupled in cichlids. The causality of this relationship remains to be investigated, just as the impact of genome rearrangements on reproductive isolation and eventually diversification in cichlids. The available data on rearrangements are sparse, but it might be that several large chromosomal rearrangements occurred before the novel chromosomes were recruited as sex chromosomes, making inferences of the driving forces of these fusions worth investigating in more detail.

A next, necessary step will be the identification of sex-determining genes and mutations causing sex chromosome turnover. This is facilitated by the close relatedness of LT cichlids allowing the generation of interspecies hybrids and also through the opportunity to study multiple sex chromosome turnover events and directions, including the repetitive occurrence of heterogamety transitions without sex chromosome change. While the repeated recruitment of the same LG as sex chromosome indicates a particularly well-suited core set of SD genes on the one hand, several transitions to otherwise not recruited LGs on the other hand question their supremacy. Although this could represent recycling of sex chromosomes to some extent, we lack the molecular and most importantly functional evidence for any master SD gene in cichlids of LT or any other radiation.

In conclusion, the estimated rapidity of sex chromosome turnover within (LT) cichlids supports the hypothesis that SD mechanisms, albeit sharing the same function of sex determination, can be extremely labile. It remains to be tested if sex chromosome turnovers are so frequent as a side effect of a generally rapid evolution of cichlid fishes or if they even drive this evolution, potentially by contributing to speciation.

## Materials and Methods

### Experimental Design

In this study, we investigated genomic and transcriptomic datasets i) to identify and characterize sex chromosomes in species covering the entire Lake Tanganyika cichlid radiation, ii) to trace the evolutionary history of sex chromosomes within the radiation to shed light on the dynamics of sex chromosome turnover in a rapidly diversifying lineage and iii) to embed our results in a broader context by comparing estimates of turnover rates and potential drivers of sex chromosome evolution to other taxa.

### Sequencing data

We used whole genome sequencing (WGS) data in the form of mapped reads in BAM files as well as variant call format from Ronco *et al*. (*24*) and raw transcriptome data from El Taher *et al*. (*28*) (see table S1 for details on species included and per species sample sizes). Based on a recent compilation of LT cichlid species (*32*), the WGS data included 225 taxa (174 described species with 4 of those represented with two local variants/populations each, and 47 undescribed species). The data further included 16 non-LT radiation haplochromine cichlid taxa (13 described species one of which represented with two local variants, and two undescribed species) and three riverine non-LT Lamprologini taxa (two described and one undescribed species) summing to a total of 244 taxa and 469 individual genomes (table S1). The transcriptome data that we used, were comprised of 66 taxa of LT cichlids (4 undescribed species, 61 described species one of which represented with two regional variants), with typically three males and three females per species (details are provided in table S1; 7 out of the 66 species had differing replicate numbers) and three tissues per individual (brain, gonad, gills; details on read numbers provided in table S3). In total, the dataset comprised 248 cichlid taxa.

### Variant calling for WGS data

Mapped reads in coordinate-sorted BAM format were derived from Ronco *et al*. (*24*) (for mapping coverage statistics see Supplementary Table 1 in Ronco *et al*. (*24*)), which are based on mapping against the Nile tilapia (*Oreochromis niloticus*) genome (NCBI RefSeq GCF_001858045.1_ASM185804v2). Unplaced scaffolds of the reference genome were concatenated lexicographically into an “UNPLACED” super chromosome.

In addition to the variant file containing all species derived from Ronco *et al*. (*24*), variants were called for each tribe separately with GATK’s (*62*) (v.3.7) HaplotypeCaller (per individual and per chromosome) and GenotypeGVCFs (per 1 Mb window), and merged with GATK’s CatVariants. Variants were further filtered with BCFtools (https://github.com/samtools/bcftools, v.1.6), applying the settings ReadPosRankSum<−0.5, MQRankSum<−0.5, FS<20.0, QD>2.0, MQ>20.0 and placing tribe-specific thresholds on minimum and maximum read depths to account for varying sample sizes (Bathybatini: 50-300; Benthochromini: 25-100; Cyphotilapiini: 50-200, Cyprichromini: 100-400; Ectodini: 250-1500; Eretmodini: 50-200; Tropheini/Haplochromini: 375-1,375; Lamprologini: 700-3,000; Limnochromini: 50-300; Trematocarini: 50-300). For the tribes Lamprologini, Tropheini/Haplochromini, Ectodini, and Limnochromini we further applied InbreedingCoeff>-0.8.

Indels were normalized with BCFtools’s norm function, monomorphic sites were excluded, and SNPs around indels were masked depending on the size of the indel: for indels with a size of 1 bp, 2 bp were masked on both sides, and 3, 5, and 10 bp were masked for indels with sizes of 3 bp, 4-5 bp, and >5 bp, respectively. Individual genotypes were then masked with VCFtools (*63*) (v.0.1.14) if they had low quality (--minGQ 20) or depth (--minDP 4). Filtered variants were phased and missing genotypes were imputed with Beagle (*64*) (v.4.1). We then retained only biallelic sites that had no more than 50% missing data prior to phasing. For sites that were polymorphic but no individual had the reference genome allele, we set the first alternative allele as reference allele.

### Approach 1 Tribe-wise association tests for sex on WGS data using GWAS

In total, we used three approaches to identify signatures supportive of sex-linked genomic regions (approach 1-3). Approaches 1 and 2 were applied on the tribe-level, the taxonomic rank above genus, but below family and which, in the case of the LT radiation, groups species with divergence times of 9.7-6.2 My. These two approaches will detect signatures of sex chromosomes shared across species, which are likely to exist due to the close relatedness of the species within a tribe.

For approach 1 (figs. S1 and S2), the phased sets of variants for tribes with at least 10 species (Lamprologini: sample size of 196 individuals representing 100 species; Ectodini: sample size of 81 individuals representing 40 species; Haplochromini including the LT-endemic Tropheini: sample size of 99 individuals of 55 species; and Cyprichromini: sample size of 21 individuals of 11 species) were each transformed into bim and bed format with PLINK (*65*) (v.1.90b). Next, we ran association tests (GWAS) for sex on these tribe-specific variant files using the univariate linear mixed model integrated in GEMMA (*66*) (v.0.97) accounting for population stratification. After visual inspection of GWAS results for potentially sex-associated regions on the tribe level (i.e., peaks or shifts of increased significance), genotypes of the 100 most significantly sex-associated SNPs for Haplochromini and Cyprichromini (broad signal for sex-association along the entire length of LG19 and LG05, respectively) and of outlier SNPs (narrow peak regions on LG15, LG20 and unplaced contigs comprising 51 SNPs with a −log10(*P*-value) > 3; extraction of the top 100 most significantly sex-associated SNPs revealed same clustering and no further sex-associated region since those SNPs were scattered across the genome) were clustered and visualized with the R package Pheatmap (v.1.0.12, https://cran.r-project.org/web/packages/pheatmap/index.html) in R (v.3.5.2) and inspected for grouping by sex.

### Approach 2 Tribe-wise tests for an accumulation of sex-specific SNPs

Approach 2 was applied to tribes that contain more than a single species (table S1 for all sample sizes) again under the hypothesis that closely related species might share a sex chromosome: We here tested for an accumulation of sites with sex-specific alleles, referred to as XY and ZW sites depending on the heterogametic sex (figs. S3 and S4), under the assumption that a sex chromosomal region will show an accumulation of sex-specific alleles due to linkage caused by suppressed recombination. To this end, we subset the unphased, filtered sets of variants per tribe and included only species for which we had individuals of both sexes (table S1). We then removed indels and sites with more than 20% missing data and more than two alleles with VCFtools (v.0.1.14). The resulting files were loaded into R (v.3.5.0) with VCFR (*67*) (v.1.8.0.9). Sex-specific sites were classified as follows: Each variant site was recoded per species as a “nosex” site if the male and the female individual had the same genotype; as “noinfo” if one or both individuals had no genotype call; as “XY” if the male was heterozygous and the female homozygous; and as “ZW” if the female was heterozygous and the male homozygous. Next, we calculated within each tribe the sum of nosex, ZW, and XY sites in windows of 10 kb with a slide of 2 kb as well as the difference between XY and ZW sites per window. Next, we calculated the mean genomewide percentage of nosex, ZW, and XY sites over all windows and multiplied these values with the number of called sites per window to obtain expected values for XY, ZW, and nosex under the assumption that most variant sites across the genome show no particular sex difference. The expected values per window were compared to the observed values using a Fisher’s Exact test with the exception of the tribe Lamprologini in which the large counts of sites per window rendered exact calculations with a Fisher’s Exact test impossible so that we applied a Pearson’s Chi-squared test. These tests will indicate windows that significantly differ from the genome-wide mean. We next inspected how windows differed from the genome-wide mean and designated and plotted a window with its corresponding *P*-value as (i) XY if the observed XY-value was greater than the expected one, and the observed ZW-value smaller than the expected one and, as (ii) ZW if the observed ZW-value was greater than the expected and the observed XY-value smaller than the expected one. If both, the observed XY-and observed ZW-values were larger than the expected value, a window was declared ambiguous and not further considered. If both, observed XY- and observed ZW-values were equal or smaller than the expected values, a window was declared nosex and not considered further. Fisher’s Exact test and Pearson’s Chi-squared *P*-values of XY- and ZW-windows were plotted together (on −log10 and log10 scale respectively) and with an overlay of the calculated XY-ZW difference for each window normalized by dividing the obtained value through the number of species analyzed. The obtained plots were inspected for the presence of LG-wide or regional shifts in XY-ZW difference and outliers from the expected XY or ZW sites. We also calculated and visualized the XY-ZW difference in each window at the species level. In order to assess a false-discovery threshold, we permutated the observed data within each tribe 100 times by randomly assigning the SNPs to the observed genomic positions. We recalculated the XY-ZW difference per window as well as the expected values. We assessed from each permutation the highest absolute XY-ZW difference of a window and the smallest *P*-value for XY/ZW sites. The largest absolute XY-ZW difference normalized by species number across all permutations within each tribe was then used as minimal threshold to define the sex-linked regions in the observed data. The lowest *P*-value of all XY/ZW windows across all permutations was −log10(*P*-value) = 5.04 (obtained in the tribe Haplochromini/Tropheini), which corresponds to a FDR ~4 after Bonferroni correction. To minimize the possibility of false positives after a comparison of all observed data across all tribes, we finally retained only drastic XY/ZW-outlier regions that in addition of exceeding the tribe-wise threshold of XY-ZW difference derived from each tribe’s permutation, also had a −log10(*P*-value)>20 (corresponding to a FDR of 2.30×10^−26^ after Bonferroni correction).

Upon a first inspection of sequence content of sex-linked regions, we noticed in the empirical RNA and DNA data XY and ZW peaks in different tribes/species within the same region on LG02 of the reference genome. This region (21.36 Mb - 21.93 Mb) is annotated with 26 protocadherin tandem gene copies. We suspect that this array of similar genes impacts mapping and hence masked this region from our sex chromosomal call. Furthermore, it has previously been shown that LG03 is a sex chromosome in *Oreochromini* and that the assembly quality of this region is poor due to presence of repetitive elements leading to difficulties in the identification of sex-linked regions on this LG (*68*). This is also reflected in our data by an excess of missing data on this LG and hence less reliable SNP data. We therefore also excluded outlier regions on LG03 as potential sex chromosome.

Since approach 2 was applied on the tribal level, we next needed to identify how many and which species are responsible for the sex chromosome signals detected within a tribe, i.e., identify the sex chromosomes on the species level from this approach. To this aim, we visualized, per window, species level XY-ZW differences in the outlier regions and clustered individual genotypes (with the possible values “homozygous reference”, “homozygous alternative”, or “heterozygous”) with divisive hierarchical clustering based on a pairwise dissimilarities matrix of Gower’s distances calculated with the R package FSA (v.0.8.30) (https://github.com/droglenc/FSA). Resulting dendrograms were inspected for grouping by sex rather than species and boxplots of species-specific XY-ZW difference for support by increased absolute XY-ZW difference. Due to the reduced sample size per species and to avoid falsepositive signals, final calls based on the outlier regions were only made if at least two species within a tribe shared the same signature of sex linkage (table S2).

### Approach 3 Species-specific association tests for sex on transcriptome data

For approach 3, identification of sex-linked regions on the species level by species-specific association tests (fig. S6), we pooled tissue-specific transcriptomes of brain, gonad and gills into one transcriptome per individual and quality filtered and trimmed these with Trimmomatic (*69*) (v.0.33) with a 4 bp window size, a required window quality of 15 and a minimum read length of 30 bp resulting in typically six multi-tissue transcriptomes per species (table S3 for read numbers). The following analysis was then run for each species: We performed reference-free *de novo* variant calling with KisSplice (*70*) (v.2.4.0) with settings “-s 1 -t 4 -u” and “--experimental”. The identified SNPs were placed on the Nile tilapia genome assembly with STAR (*71*) (v.2.5.2a) with the settings “--outFilterMultimapxNmax 1”, “-- outFilterMatchNminOverLread 0.4”, “--outFilterScoreMinOverLread 0.4”. The genome index used for this mapping was generated with the corresponding STAR parameters “--runMode genomeGenerate”, “--sjdbOverhang 124”, “--sjdbGTFfeatureExon exon” and the genome annotation file (RefSeq GCF_001858045.1_ASM185804v2). Kiss2Reference (*70*) was used to classify KisSplice variants aligned to the Nile tilapia reference genome, and kissDE (*70*) (v.1.4.0) was applied to determine variants that differed between the two sexes. The resulting files were loaded into R. The KisSplice events were filtered with the following attributes: Only SNPs were kept; SNPs placed on mitochondrial DNA or on unplaced scaffolds of the reference genome were removed; only SNPs with significant *P*-values for an allele difference between the sexes (*P*≤0.05 after adjustment for multiple testing following the Benjamini and Hochberg method (*72*)) were retained. Significant SNPs were classified as (i) “XY” if they had zero read counts in all females and a minimum of one count in at least two males and as (ii) “ZW” if they had zero counts in all males and a minimum of one count in at least two females. Next, the density of these XY and ZW SNPs was assessed in 10 kb non-overlapping windows (fig. S6 first plot).

The difference between XY and ZW SNPs per 10 kb window was then calculated and only outlier windows were kept (fig. S6 second and third plot). These outlier windows were defined as windows with a difference of XY-ZW SNPs greater than the 75th percentile value + 1.5 times the inter quartile range. We then compared the distribution of XY and ZW SNPs in all outlier windows with a paired two-sided Mann-Whitney test (fig. S6 fourth plot). If the two distributions were significantly different from each other (*P*-value<0.05), the heterogametic system was defined as the distribution (XY or ZW) with the higher total amount of SNPs. As a last step, we quantified XY or ZW SNPs of outlier windows (depending on the previously defined heterogametic system) per reference LG (corrected by chromosome length) and defined as potential sex chromosome the LG(s) with a number of SNPs higher than the 75^th^ percentile value + 3 times the inter-quartile range. In order to keep only the most extreme outliers and to further avoid false positives, only the LG(s) with a number of SNPs higher than the standard deviation were kept for this final call. In species for which a heterogametic system was identified, we further visualized all SNPs of the outlier windows of that system along the genome for illustrative purposes (fig. S6 fifth plot).

### Final sex chromosome systems definition

Sex-linked chromosomes, sex-differentiated regions and heterogametic state (XY/ZW) per species were inferred from sex-association in GWAS (approach 1), the sex-specific allele test (approach 2), and species-specific sex-differentiated site accumulations identified by allele differences test based on transcriptomes (approach 3). For approach 1 and 2, which at first result in tribe-level identification of sex chromosomes (table S2 columns B and C), we retained sex chromosome calls on the species level as follows: We required the same sex-linked region to be present in at least two species of a tribe to base a sex chromosomal call on WGS data only. This might underestimate the presence of sex chromosomes in our dataset but further reduces the number of false positives. Based on approach 3, which includes more individuals per species and was run on the species-level, we could confirm the sex chromosomes with larger sex-differentiated regions identified by approaches 1 and 2. We failed to detect some of the rather small sex-linked regions with approach 3, such as the narrow ~5 kb region in Lamprologini, which we think is due to a combination of the low number of genes present in these regions and probably low levels of expression of these genes in adults. However, approach 3 allowed us to confirm sex chromosomes shared across species and identify species-specific sex chromosomes that we would not call/identify otherwise (note that similar sample sizes and transcriptome approaches have previously been used to identify sex chromosomes e.g., (*73, 74*)). The effectiveness of our method is evidenced by our ability to identify the same signatures of sex-linkage of all three previously identified sex chromosomes of LT cichlids (*40*) (table S2).

Still, given the particularly reduced sample sizes for the small tribes in approach 2, we further decided to generate two sex chromosome call sets, a “permissive” dataset retaining all sex chromosomes identified by either approaches 1, 2, and 3 or combinations thereof and a stringent dataset, excluding all sex chromosomes that were exclusively identified in approach 2. We performed all subsequent analyses with both sets and report the results.

### Reconstruction of sex chromosome turnovers in cichlids

In order to reconstruct sex chromosome evolution across the LT radiation, we coded the final sex chromosome set as a probability matrix which included 14 different LGs identified in at least one species as sex-linked, incorporating the published data for two labstrains and a cross derived from a natural population of *A. burtoni (41, 42*) and a population of *P. philander (26*) (permissive dataset; stringent dataset 13 LGs). Note that *P. philander* was present in the current dataset with a single individual only and the *A. burtoni* samples included here derive from two different populations not allowing the confirmation of previously published data. Species, for which we could not identify a sex-linked LG and none was published to the best of our knowledge, were included in the analysis and attributed equal probability for all 14 LGs (permissive dataset; 13 LGs in the stringent dataset). We placed these sex chromosome identities onto a time-calibrated phylogeny of LT cichlids (*24*), which we pruned to include only the species studied here, using phytools. We followed the approach described in Jeffries *et al*. (*22*) and inferred ancestral sex chromosome states using a stochastic mapping approach implemented in phytools. We compared the likelihood scores (based on the Akaike Information Criterion (AIC)) for three different transition rate models, equal rates (ER), symmetrical (SYM), and all rates different (ARD), which identified ARD as the best model for transition rates between states. We simulated 1,000 stochastic character maps along the phylogeny. In addition, we ran stochastic mapping for each chromosome separately, coding the use of the chromosome as a sex chromosome in a given species as a binary (yes/no) trait to account for the fact that some tips of the phylogeny are in two or more states (i.e., two or more reference LGs showed sex-linkage likely due to chromosomal rearrangements/fusions) rather than having the equal probability of being in one out of two states. Note that for *A. burtoni*, even four different LGs have been reported as sex chromosome (*41, 42*). We then combined the 14 separate reconstructions (permissive dataset; 13 in the stringent dataset) into one phylogenetic representation. The results obtained with the two approaches were very similar and we hence continued calculations with the binary reconstructions.

We determined the timepoints of sex chromosome turnover events as points on branches where the inferred probability of using a given chromosome as a sex chromosome dropped below 0.5 for the first time starting from the tips of the phylogeny with the function densityMap of phytools. Based on Jeffries *et al*. (*22*) we did not consider species that had no detectable sex chromosome as having losses but only considered transition events that led to the emergence of a new sex chromosome, i.e., we only retained gains.

Likewise, we ran a second independent analysis with 1,000 stochastic mappings to reconstruct ancestral states for the type of heterogamety (XY/ZW). In addition to the reconstructed turnover points, we here added a turnover on the terminal branch leading to *A. burtoni*, since for this species, both XY and ZW sex chromosomes have been described (*42*).

To test if gene content or chromosome size drives the observed pattern of sex chromosome recruitment in LT cichlids, we randomly picked 30 times (the number of sex chromosome recruitments derived from ancestral state reconstruction) a window of 10 kb of the reference genome and attributed the LG containing this window as sex chromosome to a species. We simulated this operation 10,000 times and counted how many times each LG was recruited in each simulation. We than counted in how many simulations nine or more LGs were not recruited, as this was the observed pattern.

We then tested for an association of the number of sex chromosome turnovers leading to the tips of each tribe with the number of species investigated in each tribe with a phylogenetic generalized linear model (pGLS) using the R package ape (*75*) (v.5.2).

### Reconstruction of sex chromosome turnovers in other teleosts

We then ran the same two analyses for ricefishes (Adrianichthyidae), which, to the best of our knowledge, are the only fish family with detailed data on sex chromosomes with synteny inference based on a comparison to a common reference genome (*Oryzias latipes*). Information on sex chromosomes was taken from Hilgers and Schwarzer (*33*) and placed on a time-calibrated phylogeny of the family Adrianichthyidae (19 species, table S4), extracted from a recent comprehensive ray-finned fish phylogeny (*43*). We could not include sex chromosome data of three species (*Oryzias sakaizumii, Oryzias wolasi*, and *Oryzias woworae*), as these were not included in the phylogeny and no other comprehensive time-calibrated tree comprising these fishes was available to us.

To compare our data on a macroevolutionary scale, we calculated transition rates for ray-finned fishes of the Tree of Sex database (http://www.treeofsex.org/). We used the data for all Tree of Sex species that were also included in the recent comprehensive ray-finned fish phylogeny (*43*) (table S5). As several species names were not initially included in the phylogeny (*43*), we inspected species names of Tree of Sex for typos, older versions of species names and synonyms in FishBase (www.fishbase.org) and Eschmeyer’s Catalog of Fishes Online Database (https://www.calacademy.org/scientists/projects/eschmeyers-catalog-of-fishes), and corrected the names accordingly. This allowed us to map SD data for 472 species from the Tree of Sex database onto the phylogeny. We further added published data for cichlids (*26, 27, 31, 41, 42*, and this study), resulting in an additional 72 species. Sex determination data from the Tree of Sex database were simplified and coded as a probability matrix with three states, namely “XY” (including species classified by Tree of Sex as “XY heteromorphic”, “XY homomorphic”, “XO”, or “XY polygenic”), “ZW” (including species classified by Tree of Sex as “ZW heteromorphic”, “ZW homomorphic”, “ZO”, or “ZW polygenic”) and “NonGSD” (including species classified by Tree of Sex as “apomictic”, “hermaphrodite”, “ESD_other”, “pH”, “size”, “density”, “TSD”, or “other”). The final matrix is provided in table S5. Similar to our approach described above, all other species with no information on sex determination were included with an equal probability for all three states.

### Convergent evolution of sex chromosomes

We detected the same region on LG19 as sex-linked in species belonging to the tribes Haplochromini and Perissodini. Within Haplochromini, this sex-linked region was present in six species of the genus *Tropheus* (tribe Tropheini, the endemic LT Haplochromini) as well as in *O. indermauri* (a distantly related riverine haplochromine from the Lufubu river, which drains into LT). Our ancestral state reconstruction suggested an independent origin of the LG19 SD system in Perissodini, *Tropheus* and *O. indermauri*. To further investigate the hypothesis of convergence, we extracted all SNPs from LG19 which were XY in at least one of the species with a positionally overlapping XY-system from a variant call file containing all species investigated in the present study (*24*) with VCFtools. In addition to a male and a female of these eight species, we included representatives without this XY-LG19 system from the other tribes (a male and a female of each of *Xenotilapia flavipinnis*, *Plecodus elaviae*, *Petrochromis trewavasae*, *Eretmodus cyanostictus*, *Benthochromis tricoti*, *C. leptosoma*, *Limnochromis auritus*, and *Cyphotilapia frontosa*) as well as other haplochromine species (a male *P. philander*, a male and a female *A. burtoni*, a male *Ctenochromis polli*). We only kept variants with less than 10% missing data. We next extracted the two haplotype sequences of each individual for all variants in FASTA format. Assuming that the variant phasing with beagle was not error-free across whole chromosomes, we inspected the haplotypes and corrected the phasing for the eight LG19-XY species. This was done such that for sites where an XY male was heterozygous while the corresponding XX female was homozygous, the allele of the male shared with the female was designated as haplotype 1 (the presumed X-allele) and the other allele as haplotype 2 (the presumed Y-allele). We then inferred a phylogenetic tree by maximum likelihood with IQ-TREE (*76*) (v.1.7-beta12) under the GTR+F+ASC substitution model to account for ascertainment bias and assessing branch support with 1000 ultrafast bootstrap approximations (*77*). We rooted the obtained phylogenetic tree in accordance with the species tree (Fig. 1).

### Defining the degree of sex chromosome differentiation, potential sex-determining regions, and candidate genes

On the above-defined sex chromosomes, we characterized species-specific sex-differentiated regions by counting the numbers of XY- and ZW-SNPs in windows of 10 kb. The density of XY- or ZW-windows is shown in fig. S10. We defined the size of the sex-differentiated region as the proportion of the LG covered by windows that have a density of sex-specific SNPs that is more than twice as high as the genome-wide mean over all windows such that the sum of all sex-differentiated windows defines the cumulative length of the sex-differentiated regions and the minimum and maximum window coordinates define the range of the sex-differentiated region on the LG. We tested for an association between sex chromosome differentiation and the estimated age of origin of the sex chromosome derived from the turnover point with a phylogenetic generalized linear model (pGLS) using the R package ape. From the results of approaches 1-3, we identified sex-differentiated regions shared between several species and overlaid these with candidate genes involved in sex determination and pigmentation. Pigmentation genes in the reference genome were defined over gene ontology annotations including the term “pigmentation” and its child terms. We also retrieved orthologous sequences of the Nile tilapia to the medaka pigmentation genes defined by Braasch *et al*. (*78*) over Biomart, Ensembl release 96 (www.ensembl.org). Since this Nile tilapia genome is a different genome release than the reference genome used by us, we searched the NCBI database for the obtained Ensembl gene IDs and translated them to the assembly version that we used with the NCBI Genome Remapping Service. Candidate genes for SD included genes previously identified through a literature search (*79, 80*) and a gene ontology analysis based on a GO annotation matching the word “sex” (list of gene IDs of candidate genes for SD and pigmentation in table S6). We further investigated all annotated genes that were partially or fully included in the window(s) with the maximum number of sex-specific SNPs on the sex chromosome (table S2).

### Statistical analysis

Statistical parameters and applied tests are reported in the main text, corresponding Materials and Methods sections and figure legends where appropriate. All statistical analyses were performed in R (v.3.5.0 and v.3.5.2, detailed above including used R packages).

## Acknowledgements

**General**

We thank Daniel Jeffries and Guillaume Lavanchy for code sharing for ancestral state reconstructions and Milan Malinsky for discussions on sex-specific sites identification. Calculations were performed at the sciCORE (http://scicore.unibas.ch/) scientific computing center at the University of Basel, with support by the SIB (Swiss Institute of Bioinformatics), and at the Abel computer cluster at the University of Oslo.

## Author contributions

AB designed the study with input from AE, FR, and WS; AE and AB analyzed all data, MM performed variant calling and phylogenetic reconstructions for convergent sex chromosome evolution, and helped with ancestral state reconstructions as well as with statistics, FR helped in analyzing data for ancestral state reconstructions and sex-specific site identifications, AB and AE wrote the manuscript with final contributions from all authors. All authors read and approved the final manuscript.

## Data and materials availability

Sequencing data used in this study were published elsewhere (*24, 28*). All other data needed to evaluate the conclusions in the paper are present in the paper and its Supplementary Materials.

