## Supplementary Figures 1-11 for "Dynamics of sex chromosome evolution in a rapid radiation of cichlid fishes"

**This PDF file includes:**

Figs S1-S11

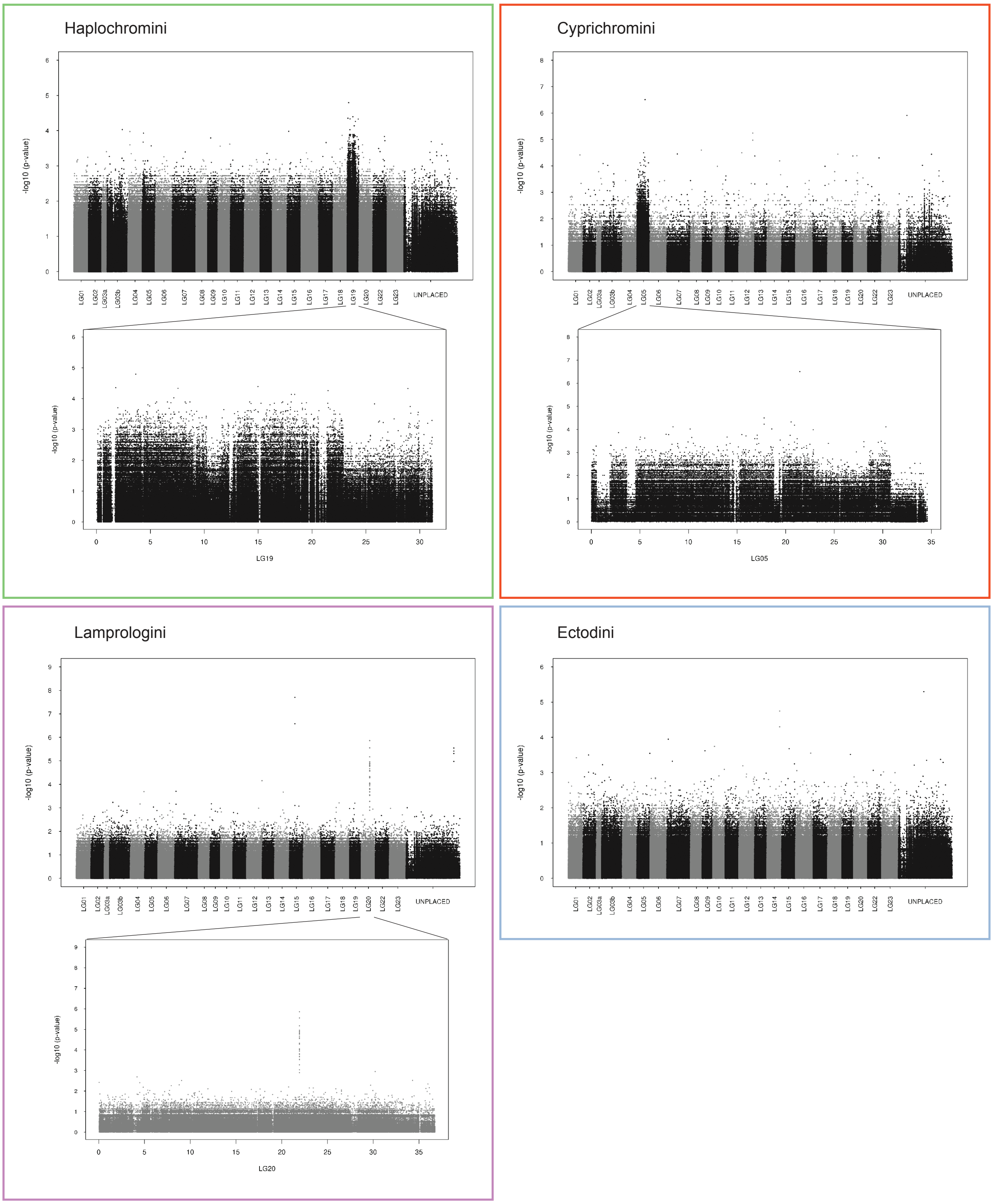


**Fig. S1. Results of tribe-wise GWAS for an association with sex (approach 1).** Manhattan plots of a GWAS analysis for an association with sex per tribe (indicated next to the plot) using the reference genome of the Nile tilapia (*Oreochromis niloticus*). Interchanging black and gray colors delimit chromosomes. Unplaced scaffolds were concatenated into an "UNPLACED" chromosome for visualization. Insets show zooms of regions with an accumulation of SNPs that show an association with sex, i.e. sex chromosomes.

**Supplementary Figure 2**

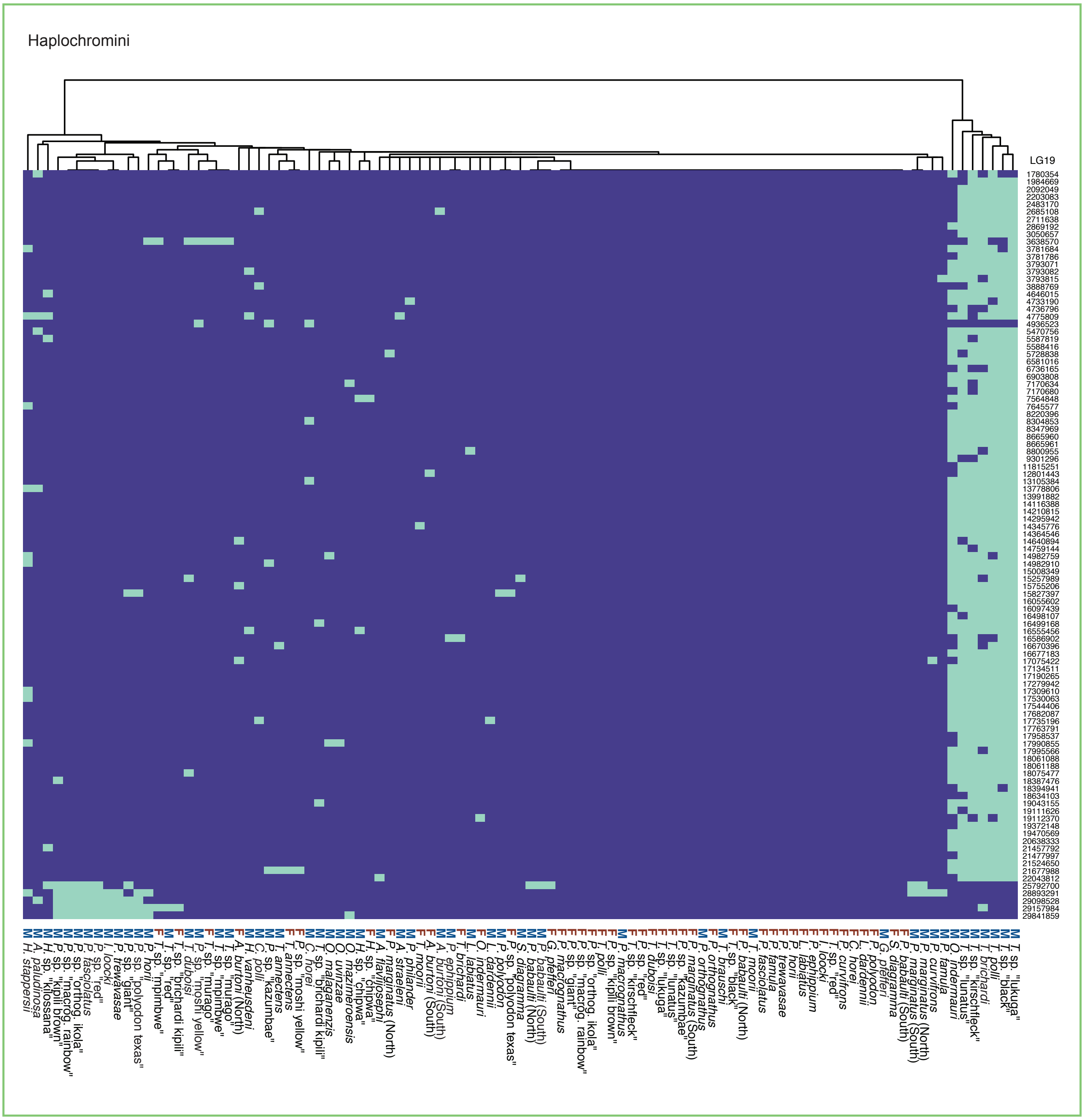

**Supplementary Figure 2 (continued)**

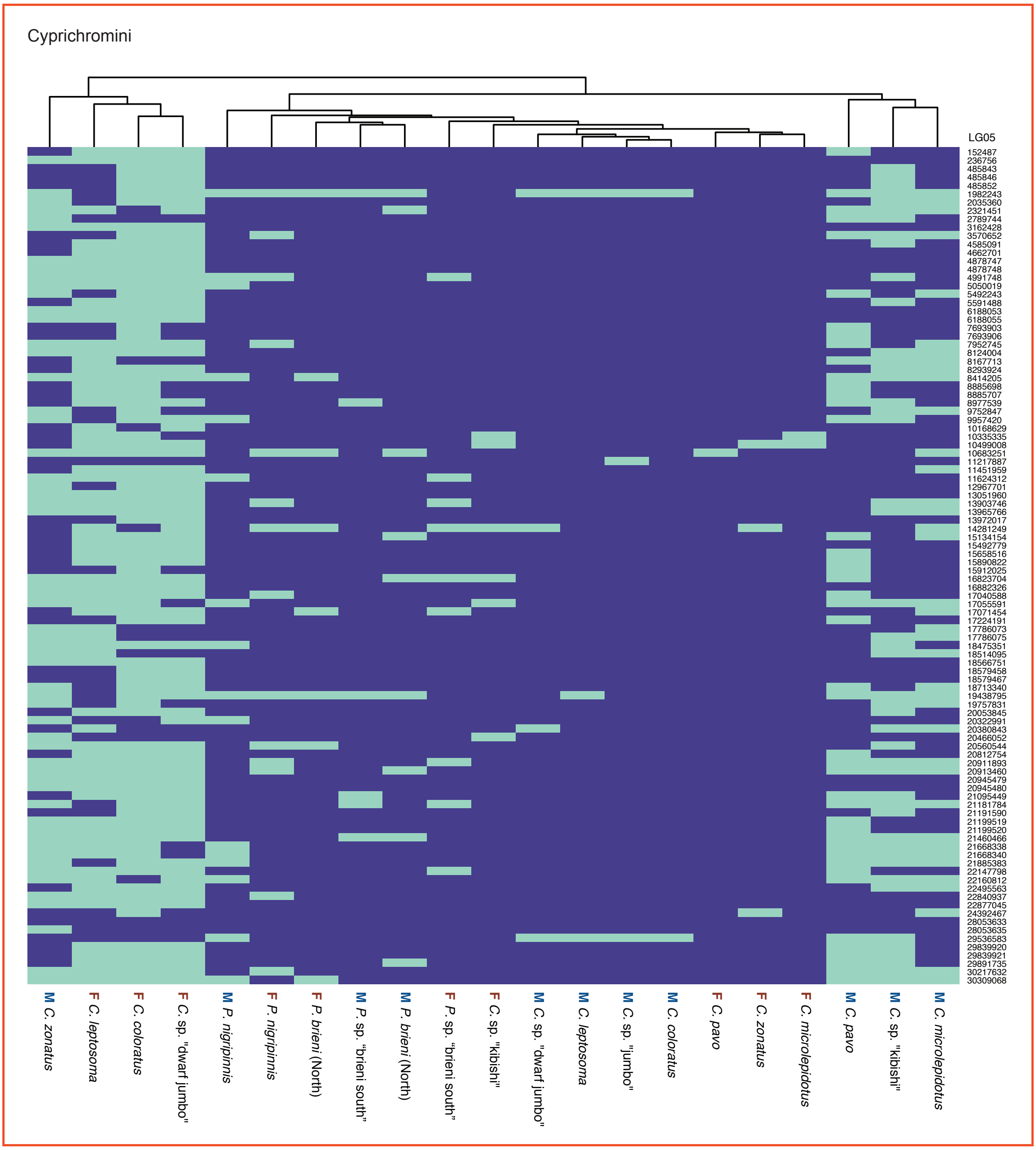

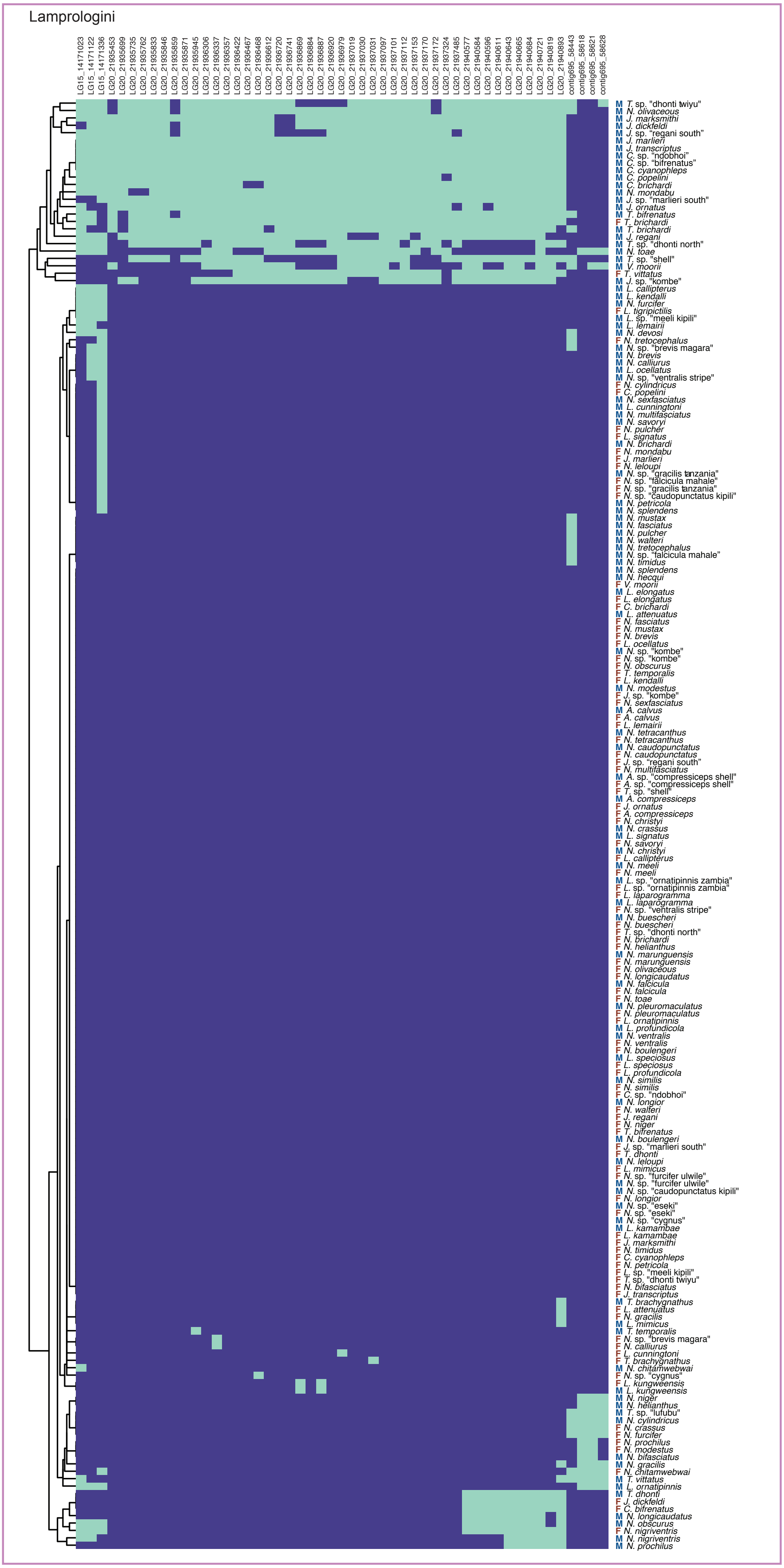


**Fig. S2. Genotypes of GWAS outlier regions (approach 1).** Heatmaps show individual genotypes for outlier SNPs (Haplochromini and Cyprichromini 100 most significant SNPs, Lamprologini three outlier regions on LG15, LG20 and an unplaced contig) of sex-linked regions detected by GWAS shown in fig. S1; purple: homozygous; green: heterozygous.

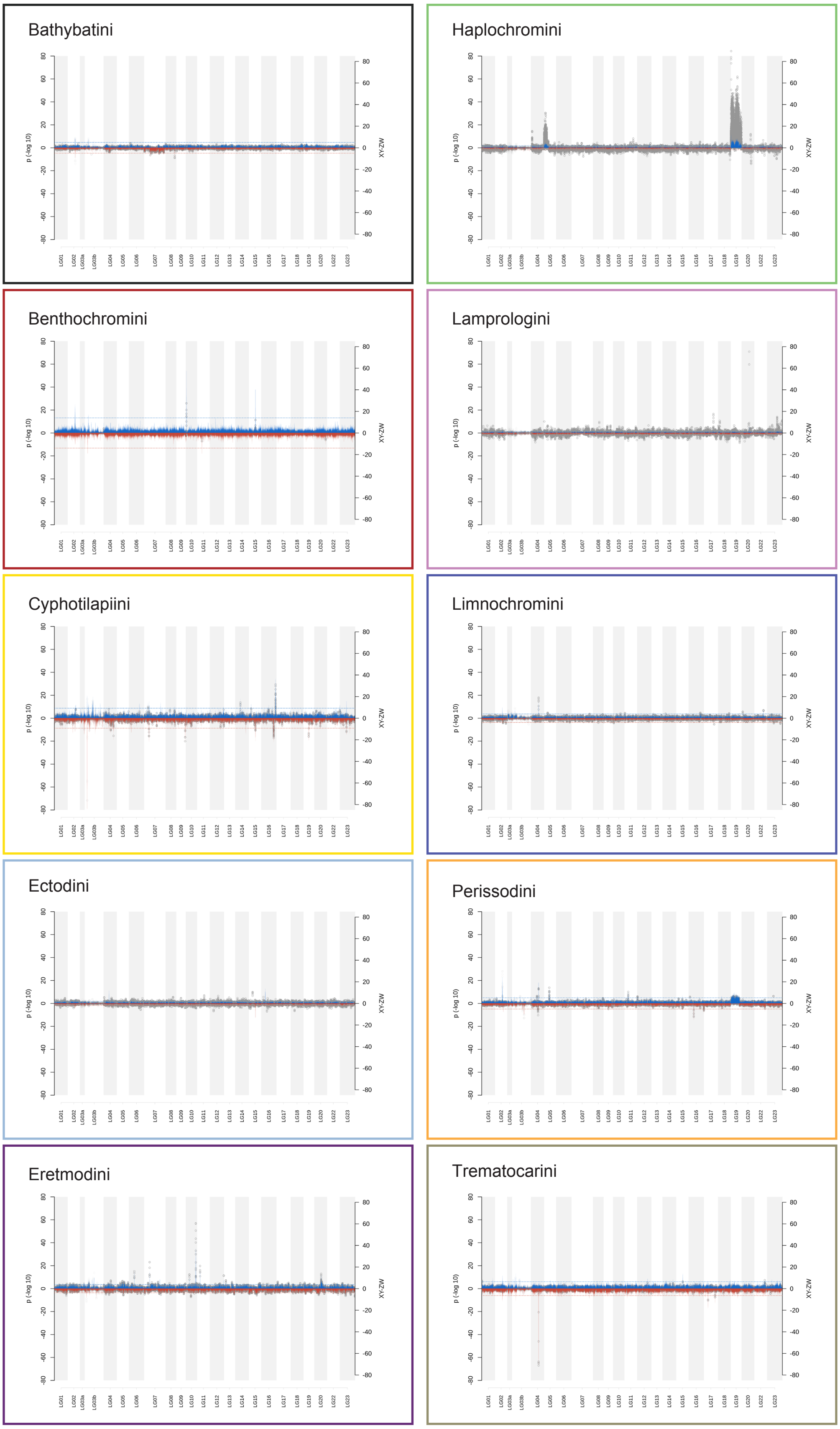


**Fig. S3. Sex-specific SNP windows per tribe (approach 2).** Gray dots show results for tests of overrepresentation of XY- (left Y-axis, positive scale) and ZW- (left Y-axis, negative scale) SNPs in windows of 10 kb with a slide of 2 kb. Repetitive sequence regions on LGs 02, 03a and 03b were excluded in the final call set of sex chromosomes and are plotted in lighter gray. Background shadings refer to LGs of the reference genome. Vertical lines indicate XY-ZW SNP difference per window normalized by the number of species (right Y-axis), windows with more XY-SNPs are plotted with blue lines, those with more ZW-SNPs with red lines. Dotted horizontal lines indicate the largest absolute XY-ZW difference normalized by species number obtained in each tribe over 100 permutations (right Y-axis).

#### Supplementary Figure 4

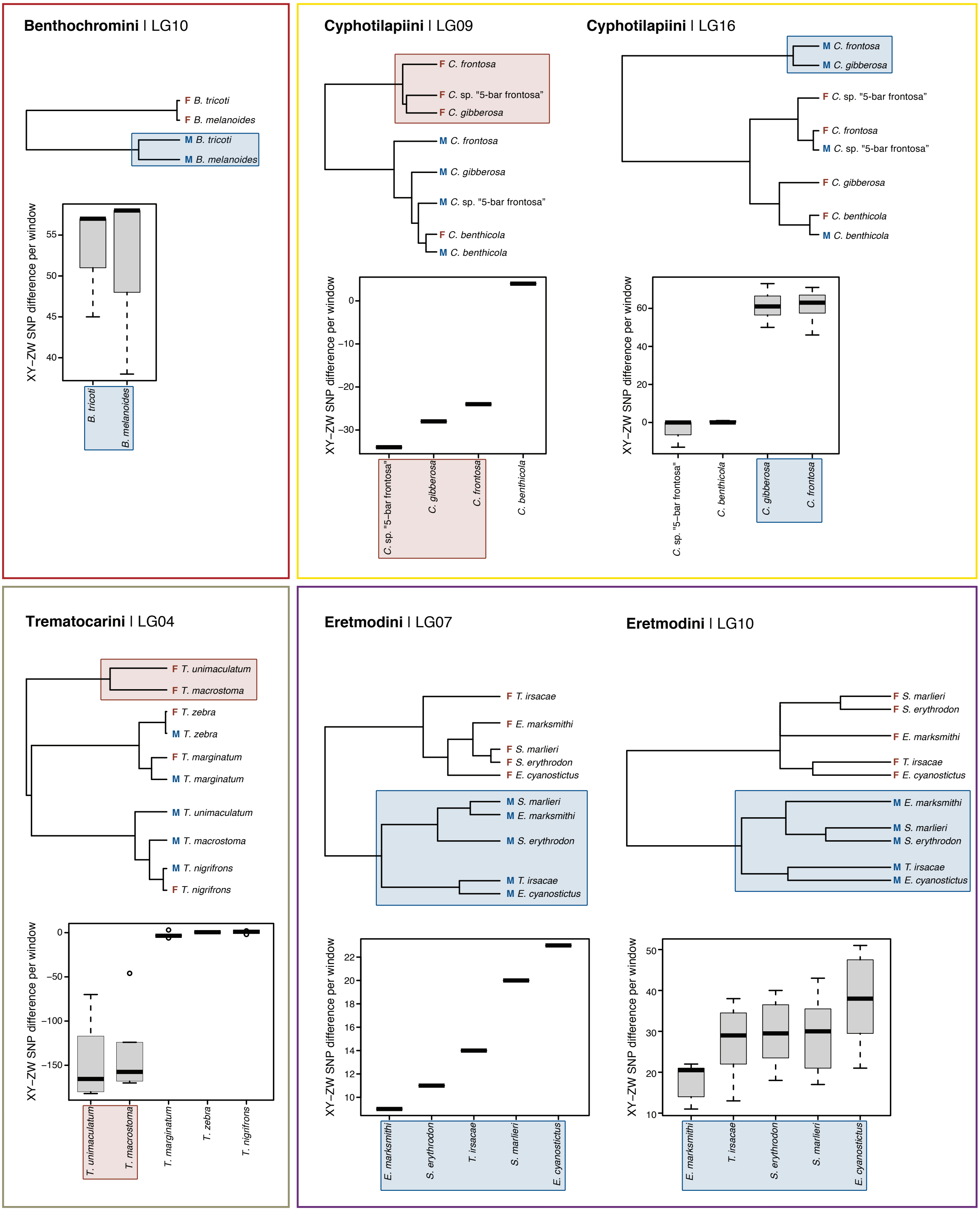

**Supplementary Figure 4 (continued)**

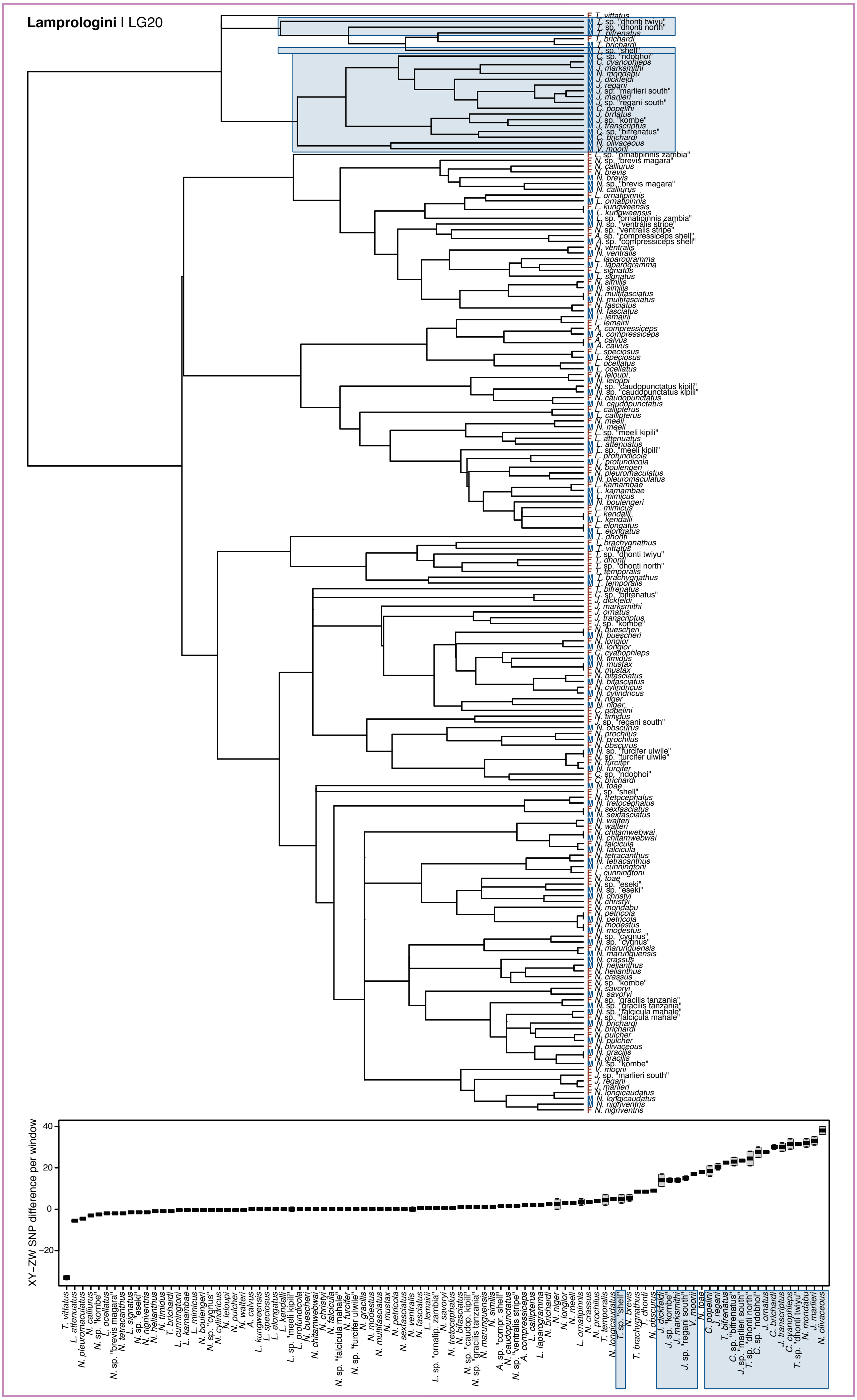

#### Supplementary Figure 4 (continued)

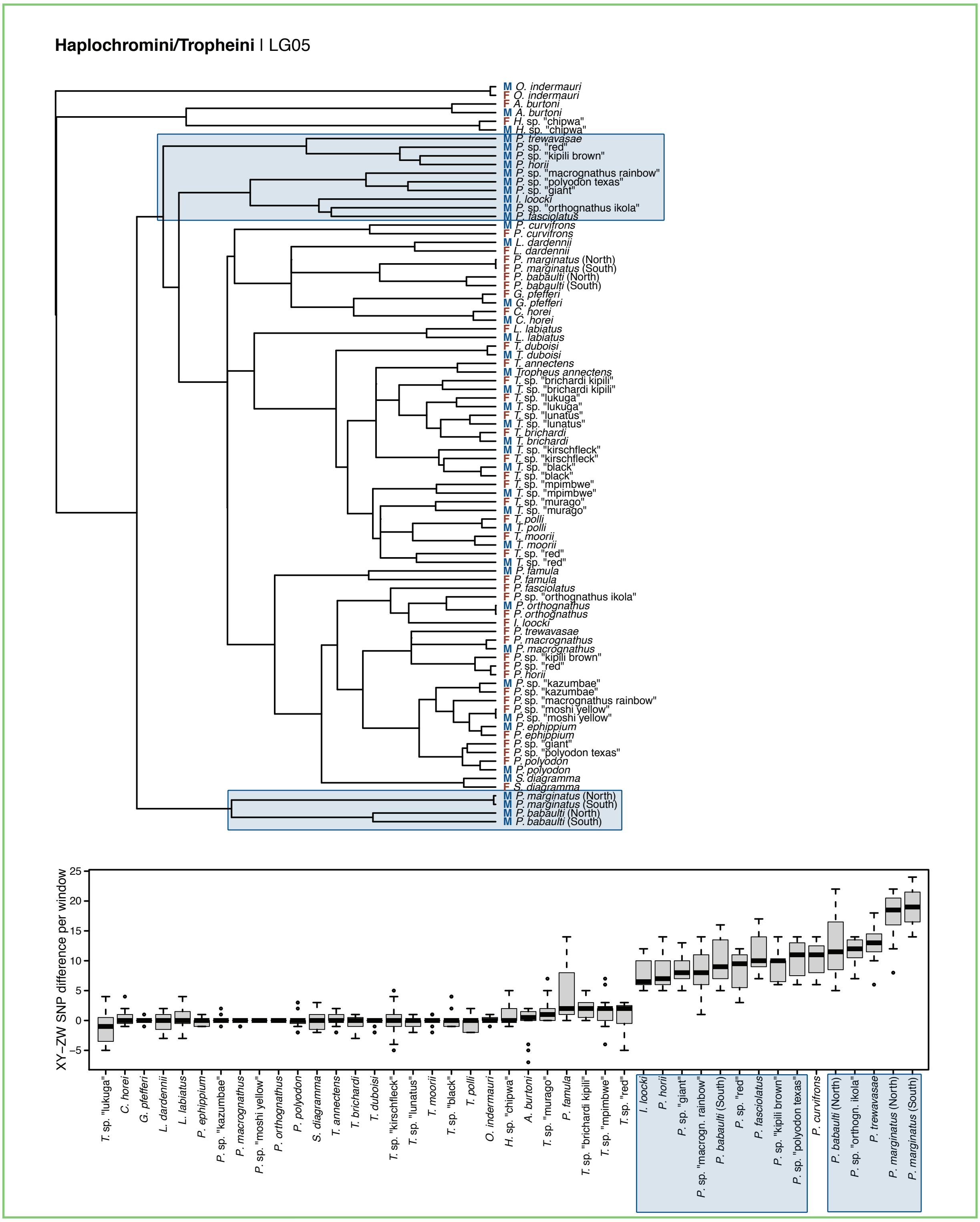

**Supplementary Figure 4 (continued)**

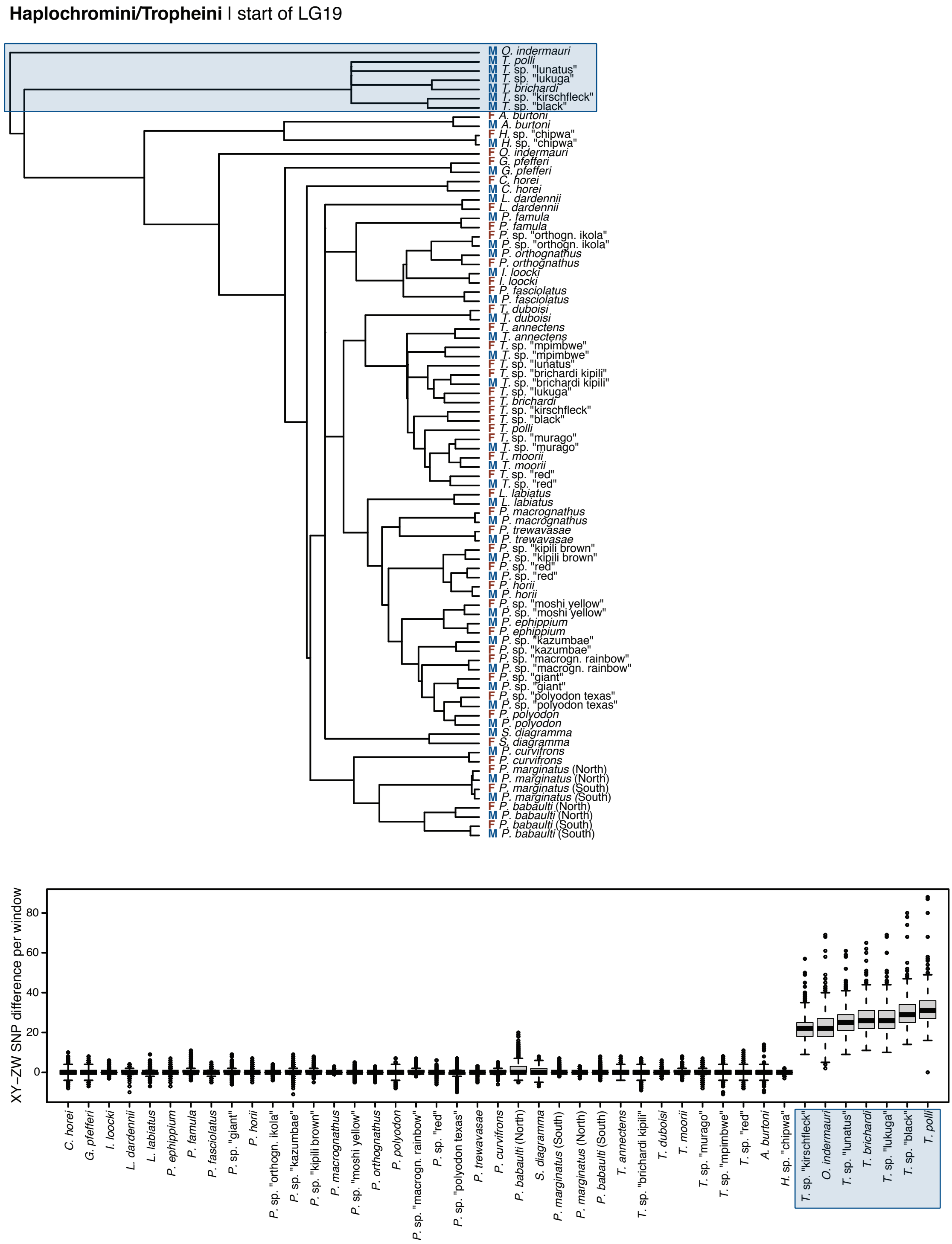

#### Supplementary Figure 4 (continued)

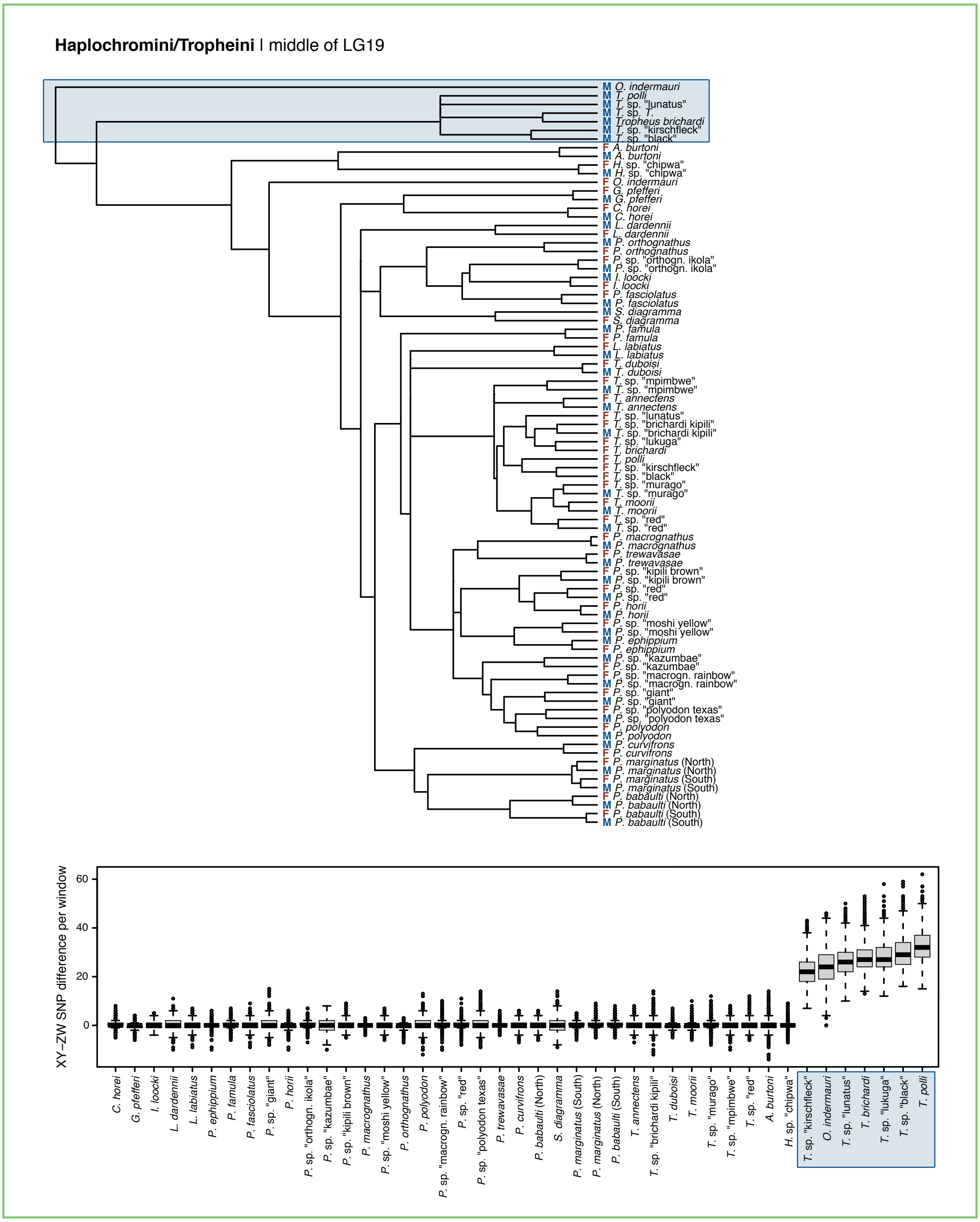

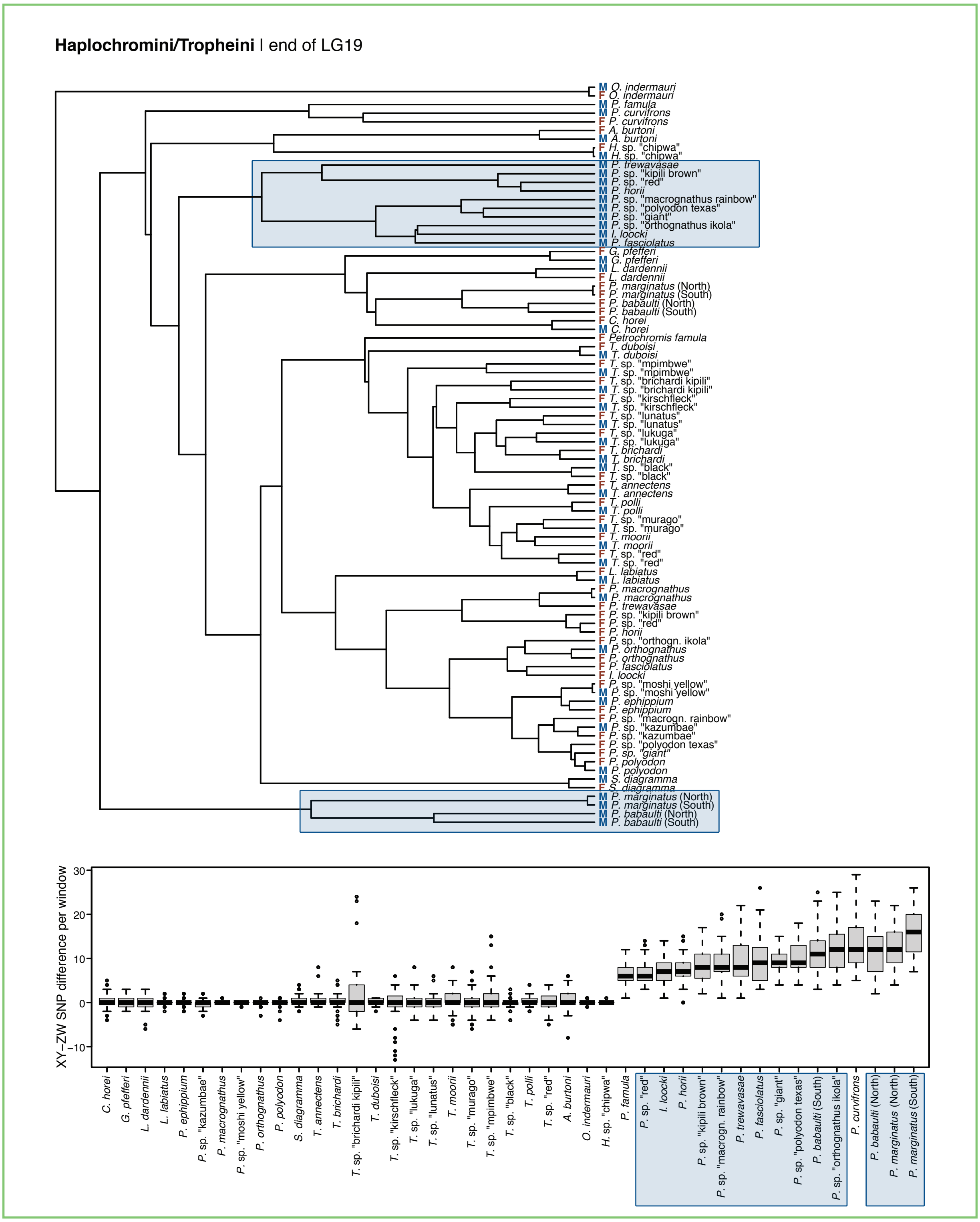


**Fig. S4. Species values of sex-specific outlier regions (approach 2).** Dendograms of samples based on divisive hierarchical clustering of genotype values in outlier windows of approach 2 with individuals clustering by sex colored in blue for XY-systems and in red for ZW-systems and boxplots of per species XY-ZW difference in XY/ZW outlier windows (fig. S3) with color-coding based on the dendograms. Boxplot center lines represent the median, box limits the upper and lower quartiles, and whiskers the 1.5x interquartile range; points represent outliers.

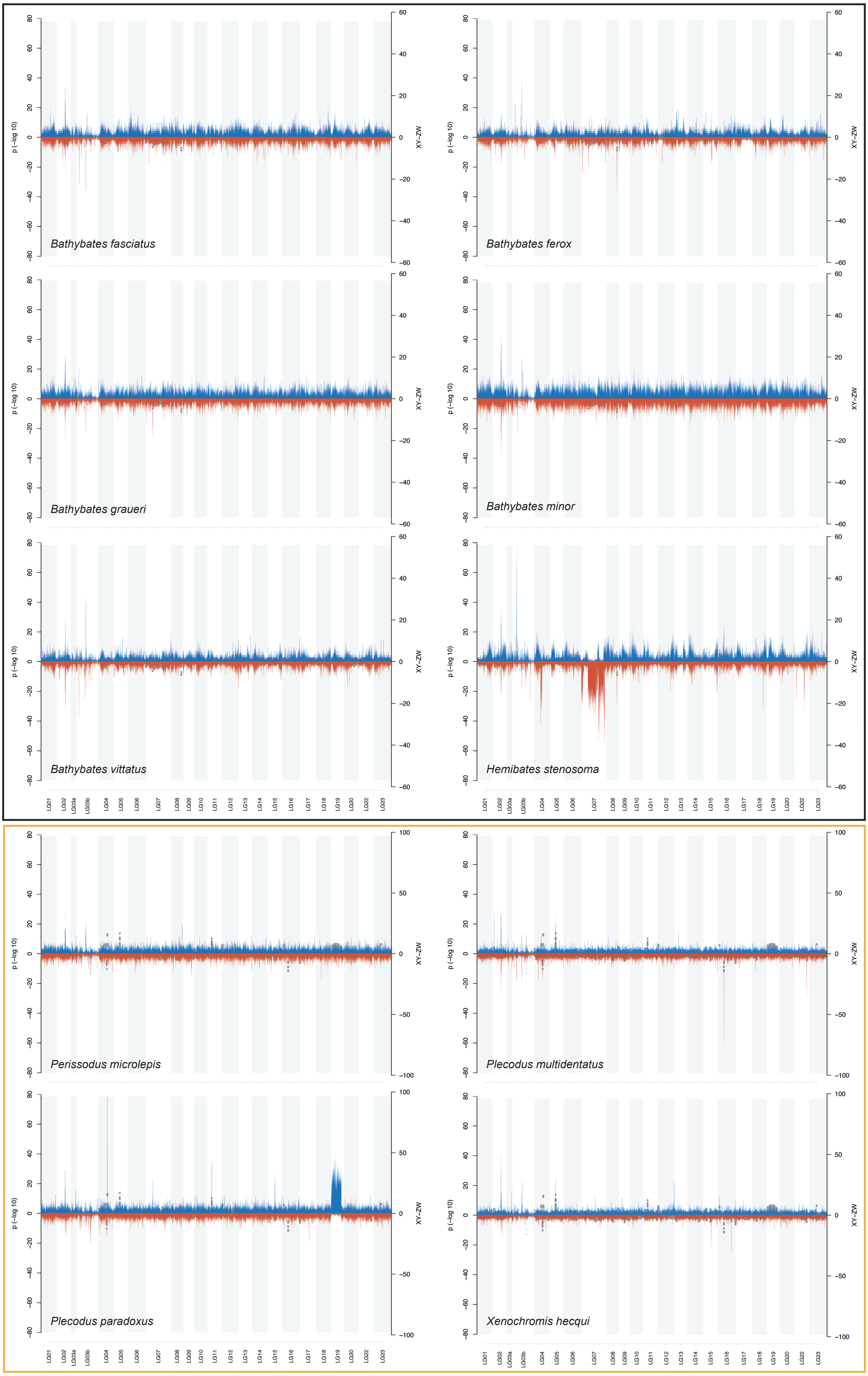


**Fig. S5. Chromosome-wide sex-specific alleles in *Hemibates stenosoma* and *Plecodus paradoxus*.** Visualization of per species XY-ZW differences in windows of 10 kb with a slide of 2 kb overlaid on the tribe-wise test results for sex-specific SNPs for Bathybatini and Perissodini (fig. S3).

#### Supplementary Figure 6

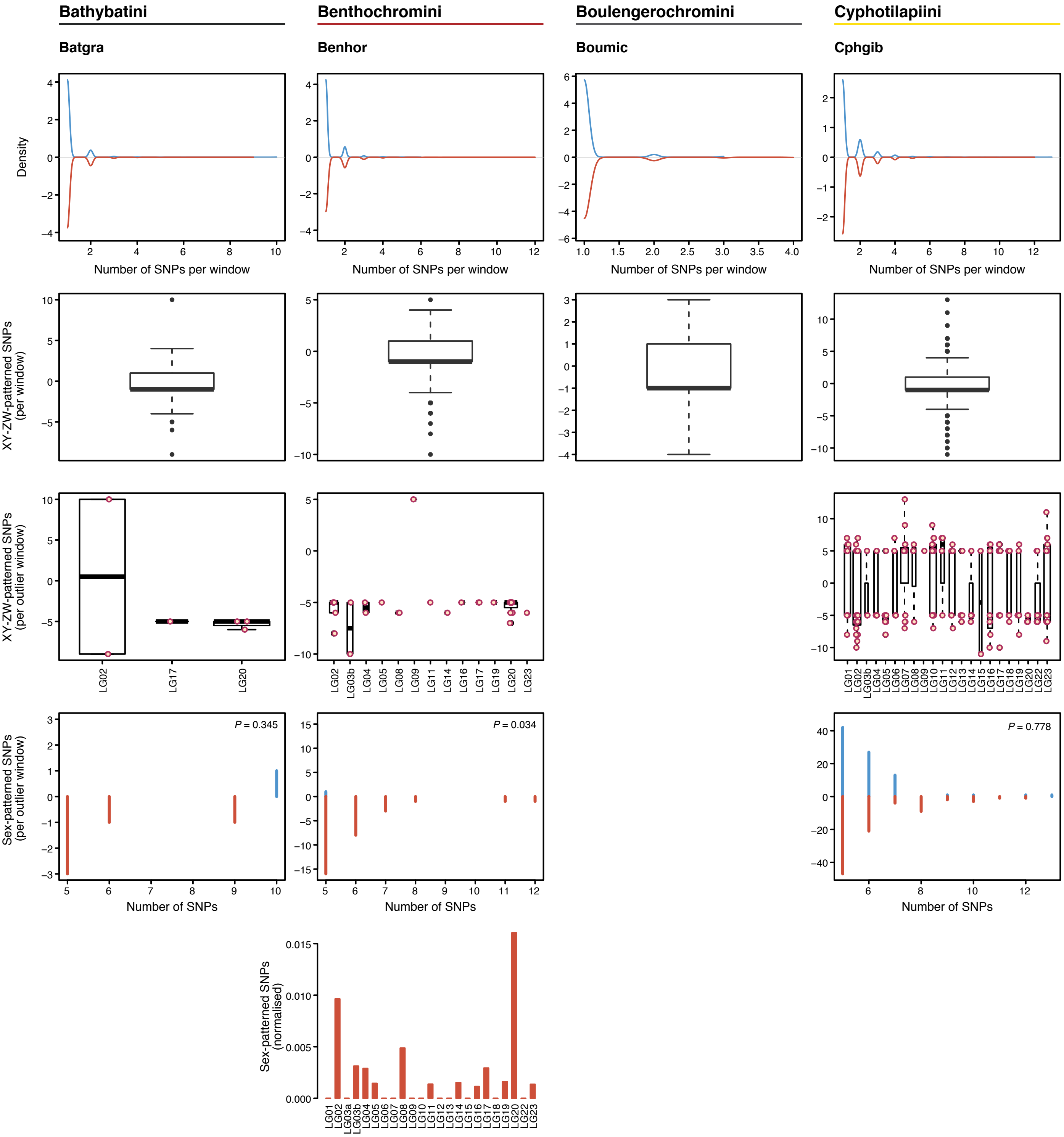

#### Supplementary Figure 6 (continued)

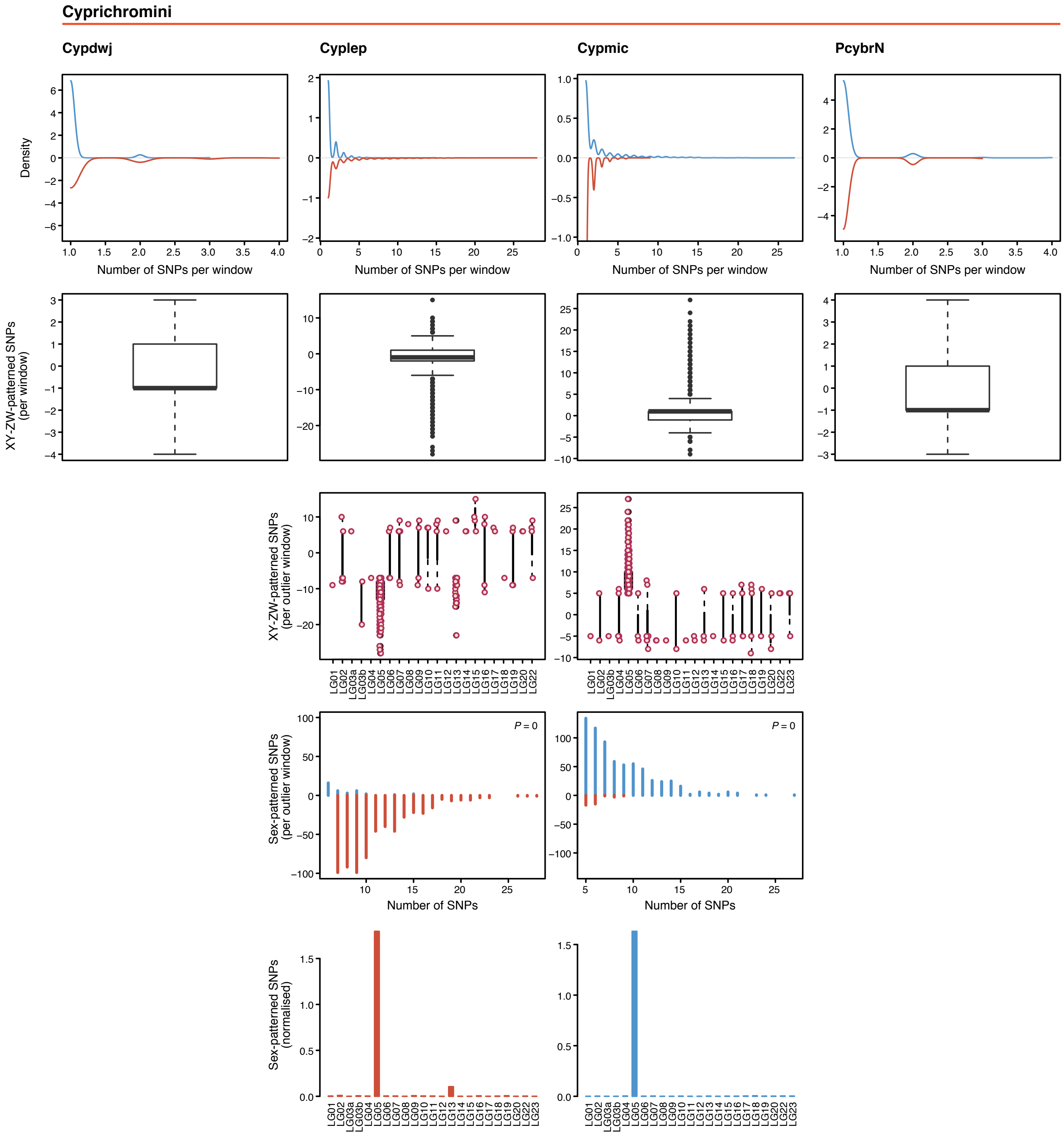

#### Supplementary Figure 6 (continued)

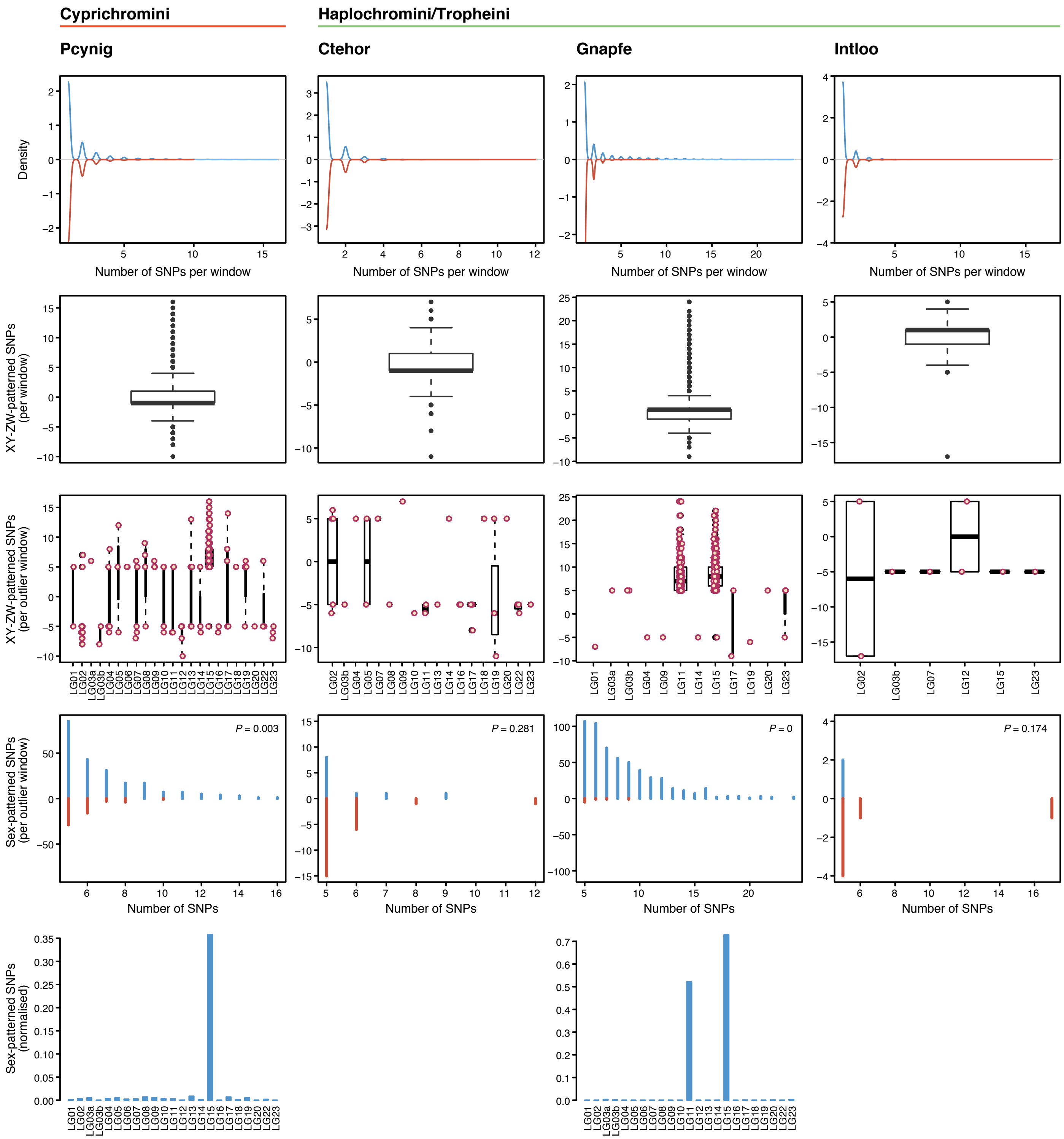

#### Supplementary Figure 6 (continued)

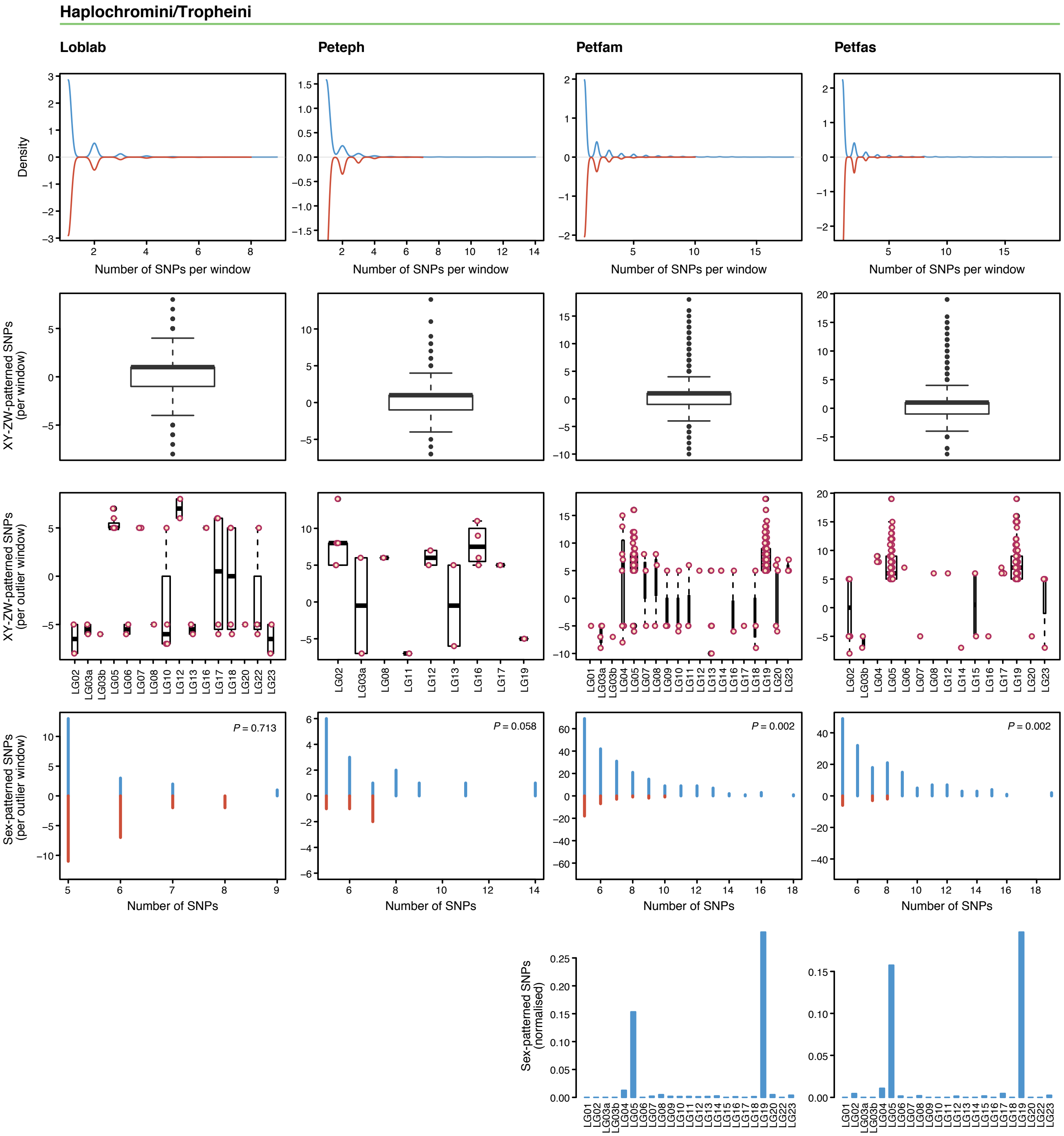

#### Supplementary Figure 6 (continued)

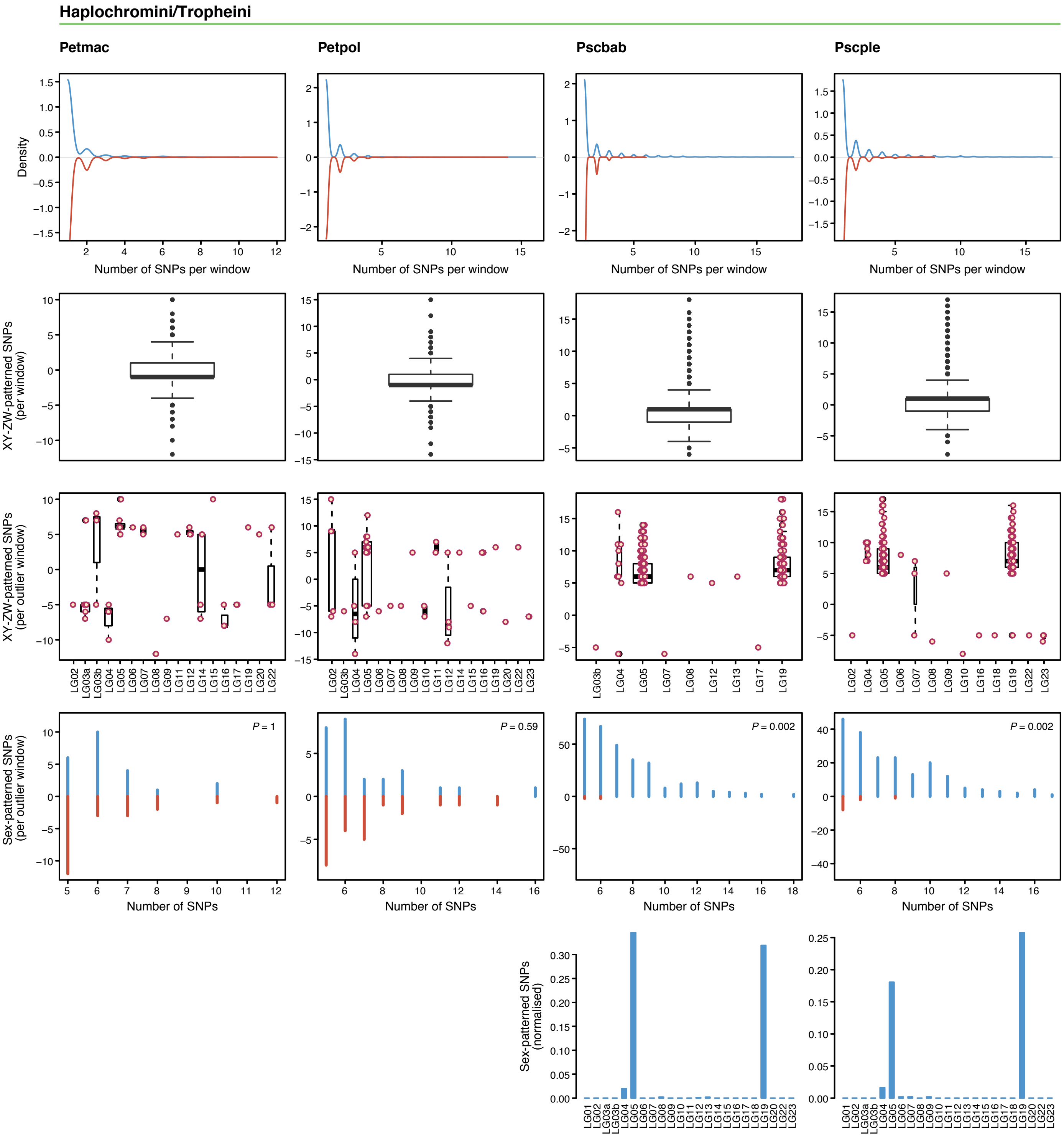

#### Supplementary Figure 6 (continued)

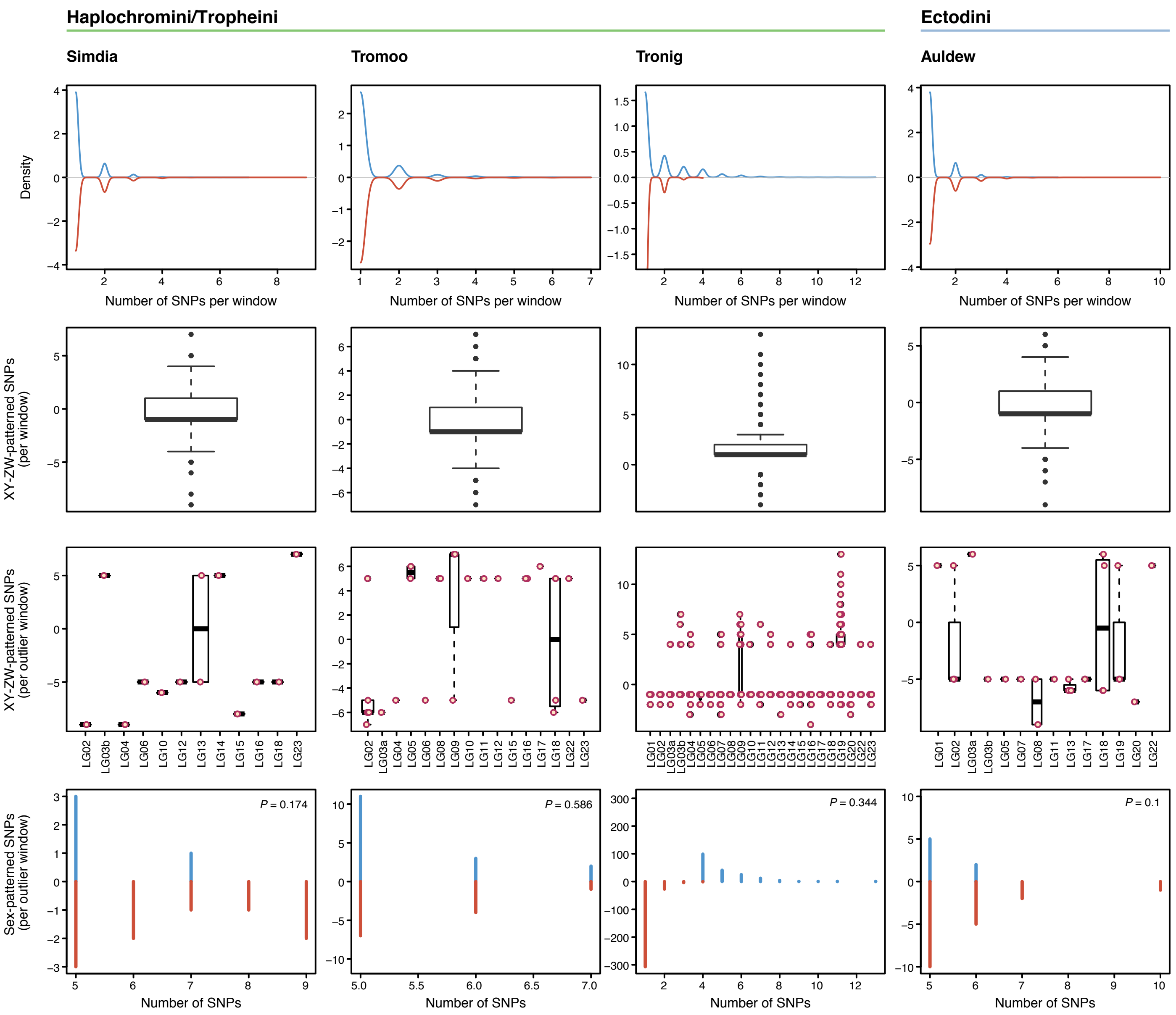

### Supplementary Figure 6 (continued)

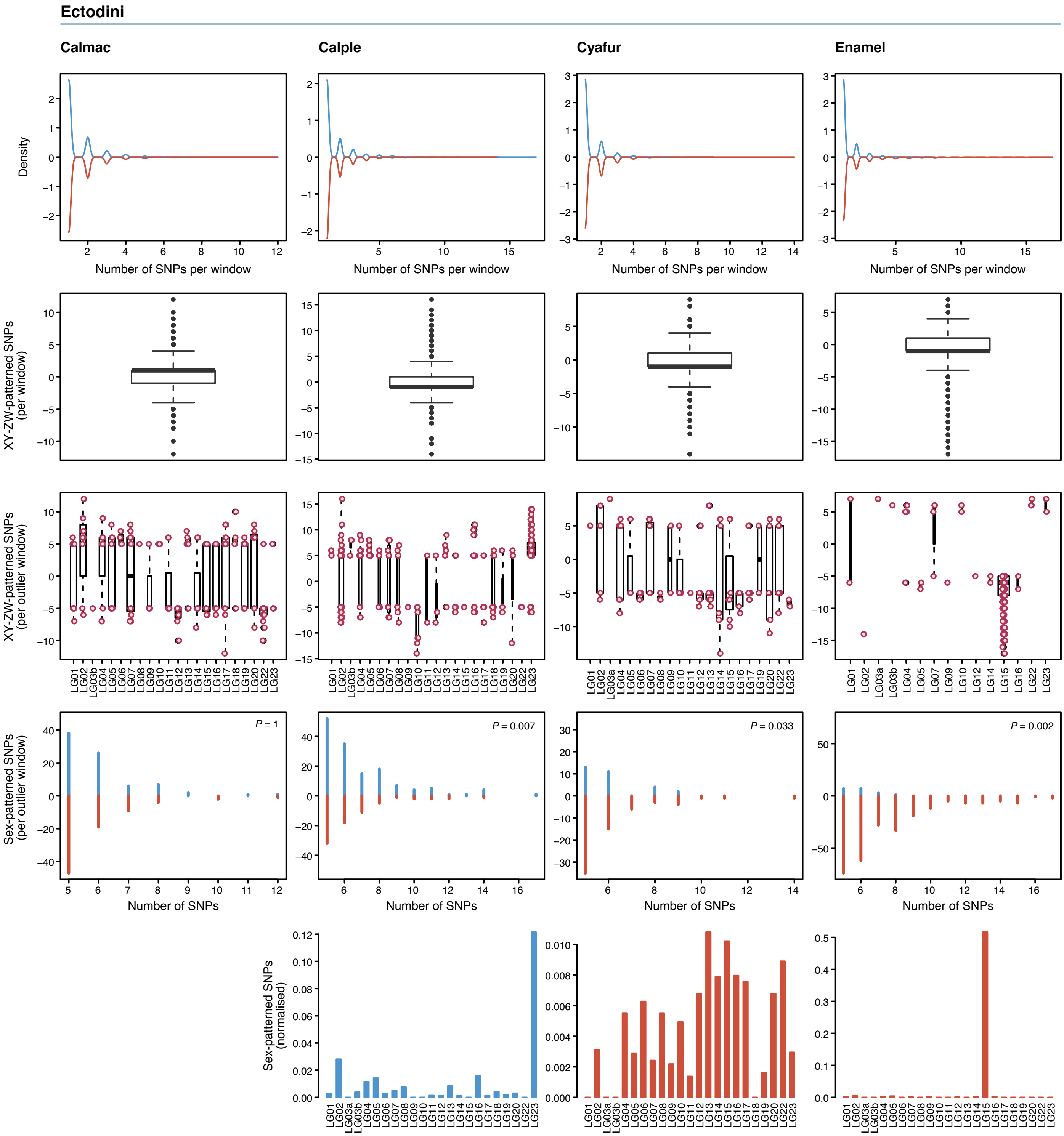

#### Supplementary Figure 6 (continued)

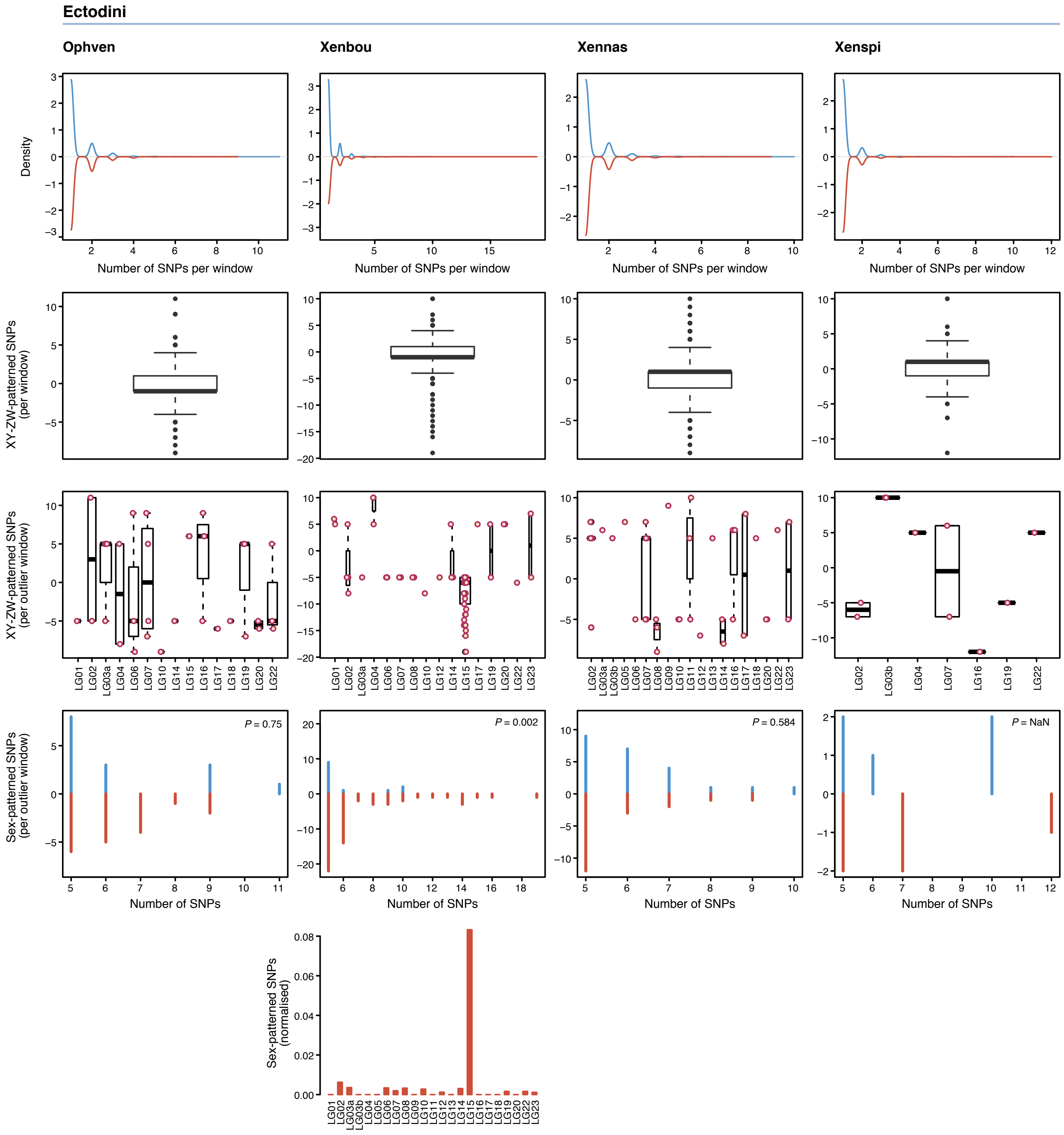

#### Supplementary Figure 6 (continued)

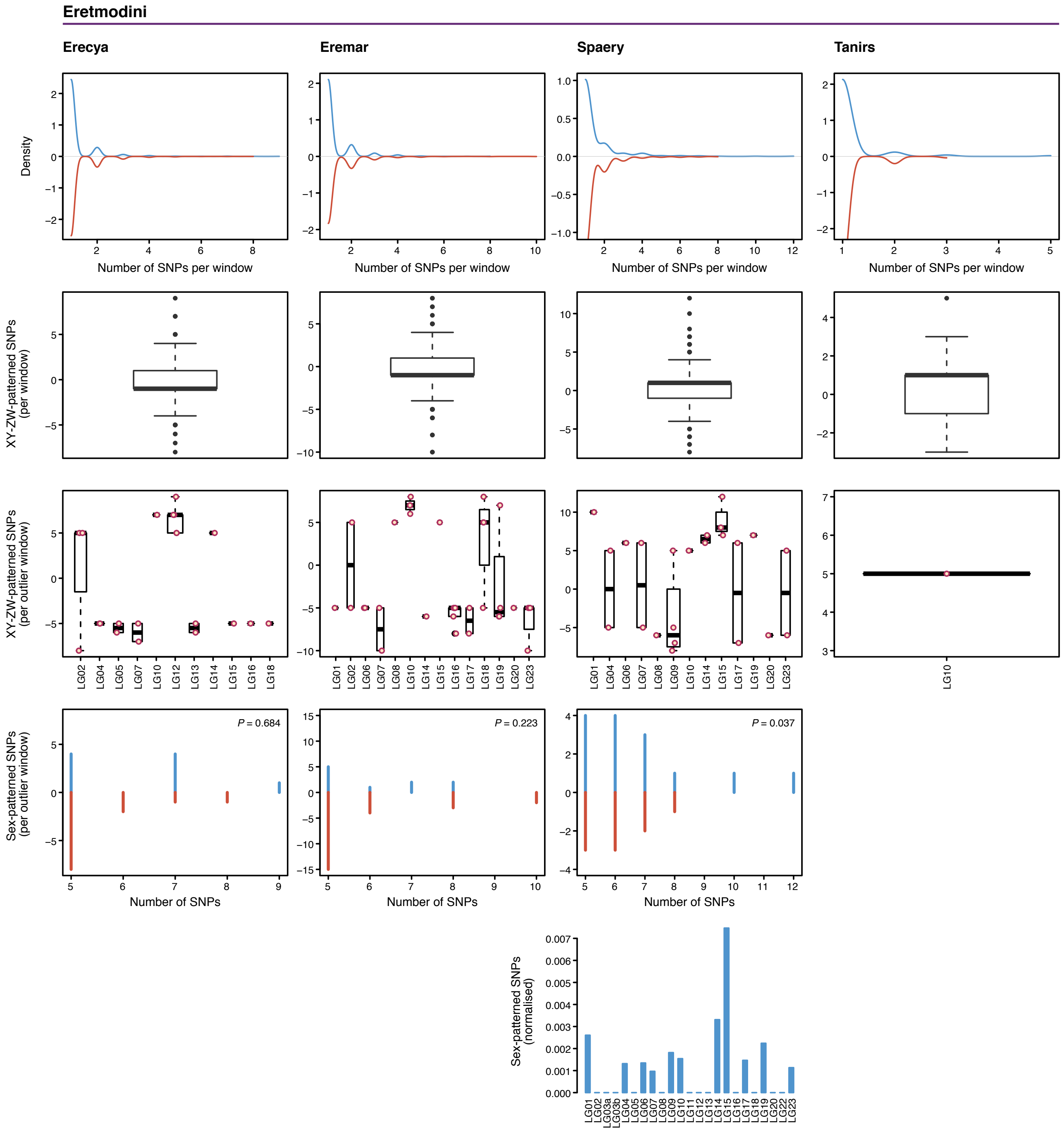

#### Supplementary Figure 6 (continued)

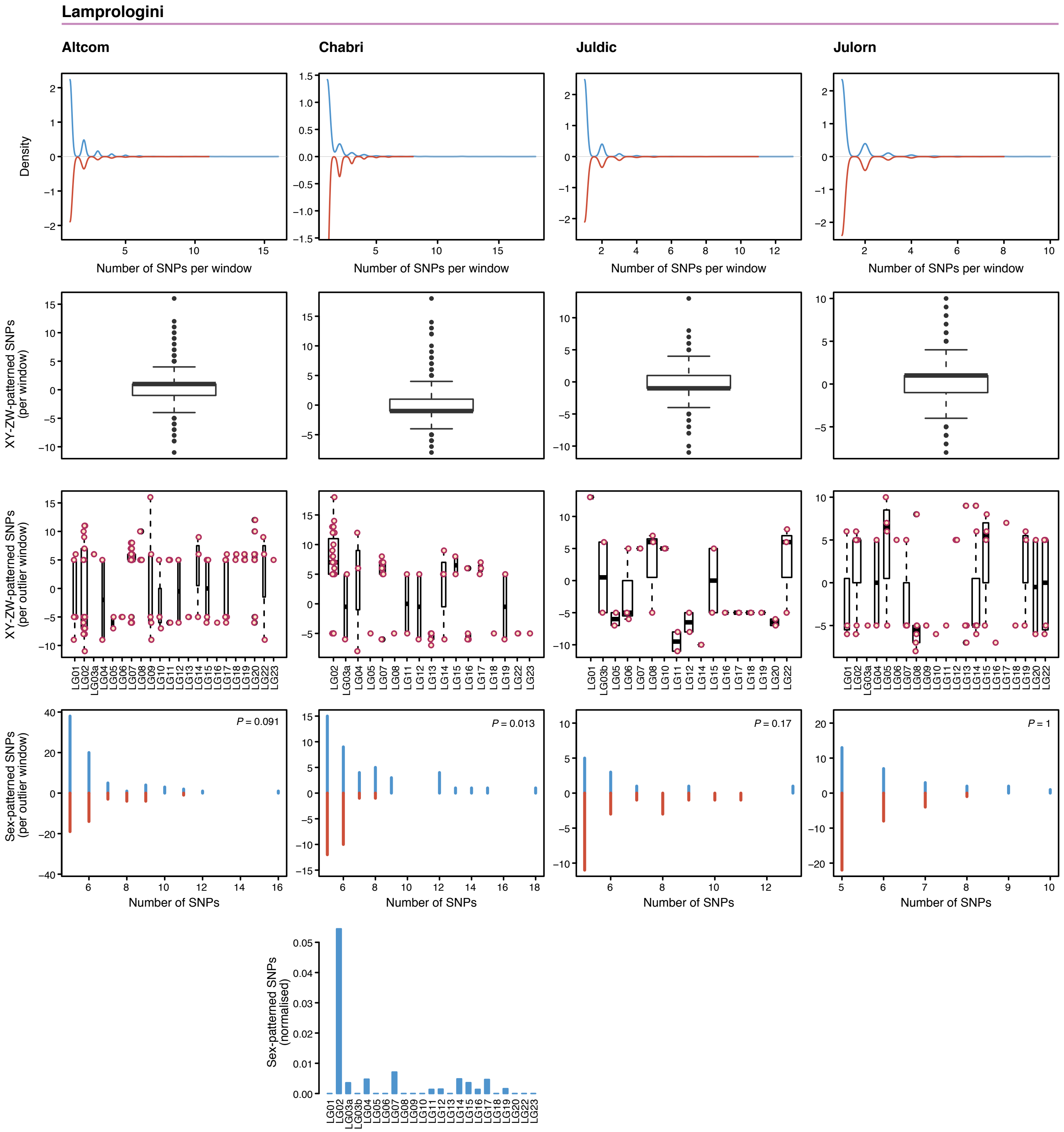

#### Supplementary Figure 6 (continued)

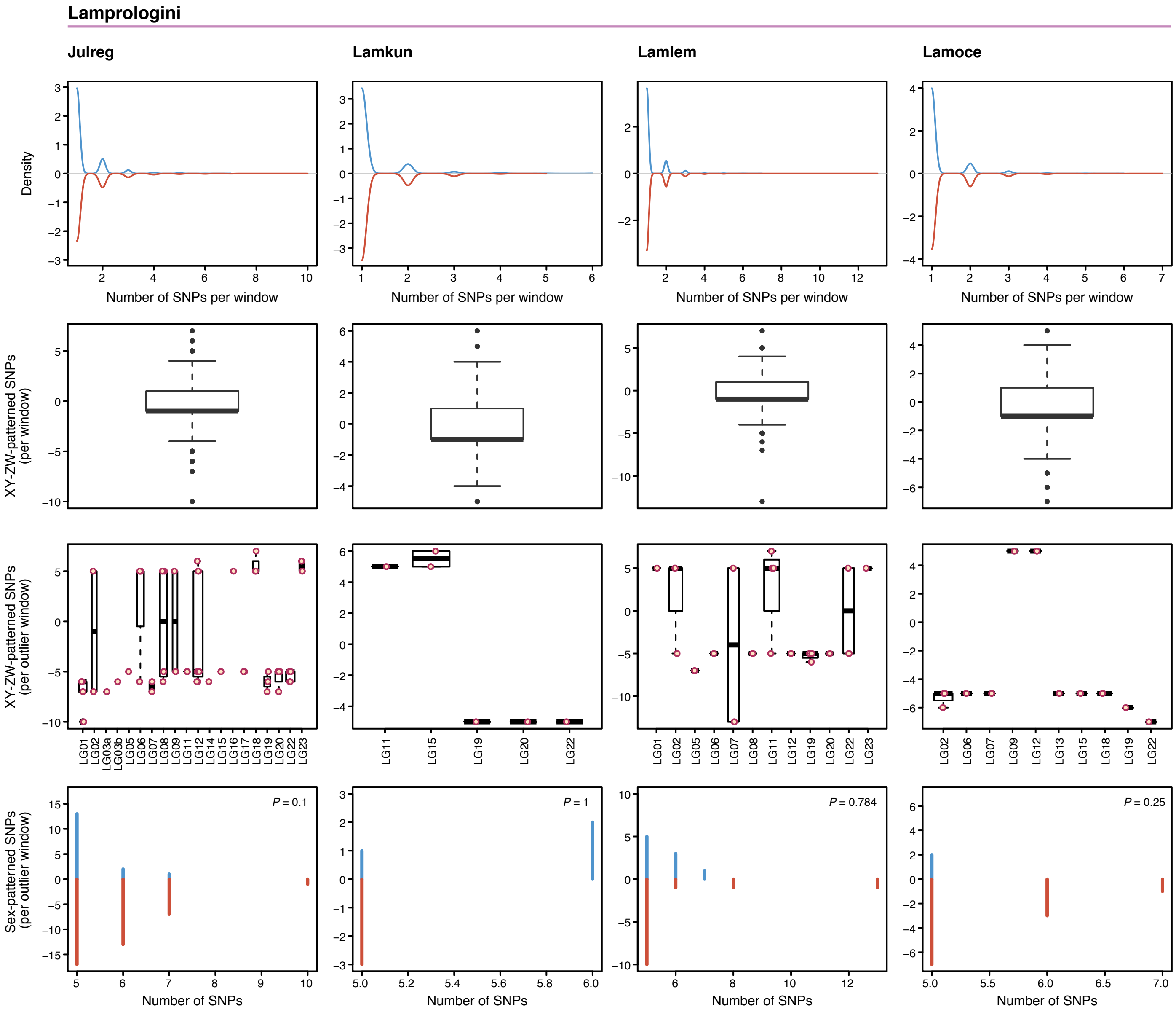

#### Supplementary Figure 6 (continued)

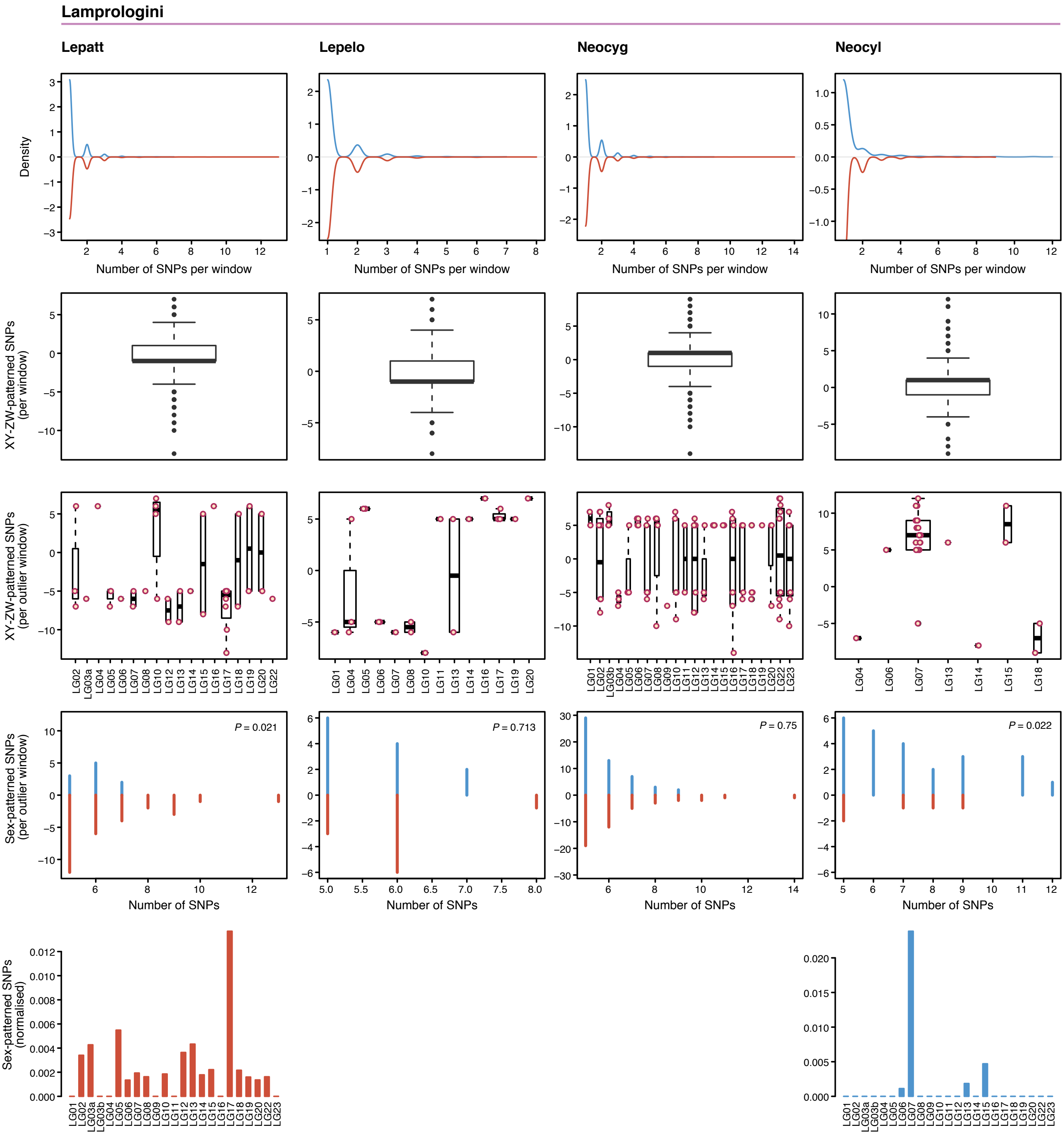

**Supplementary Figure 6 (continued)**

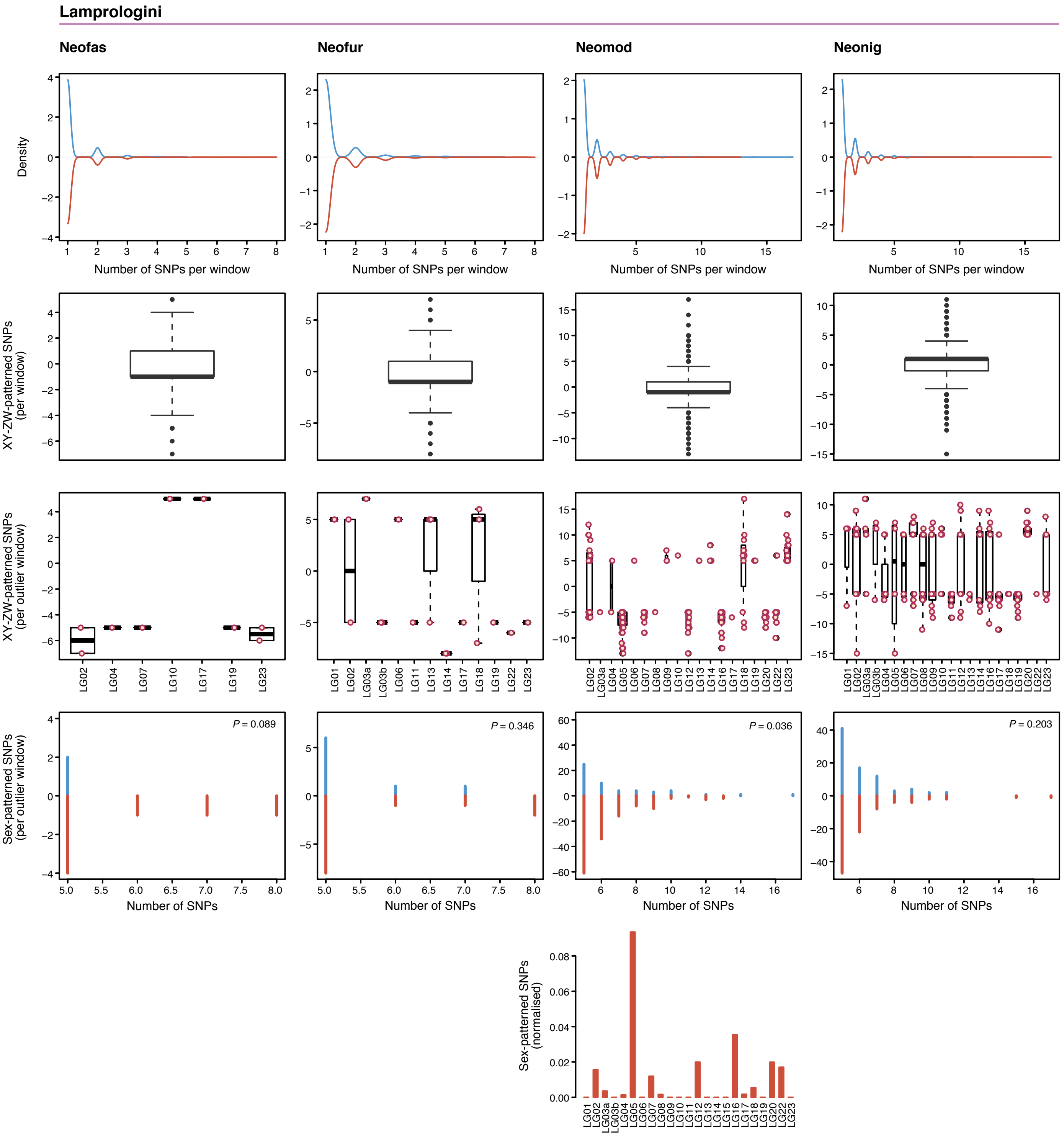

#### Supplementary Figure 6 (continued)

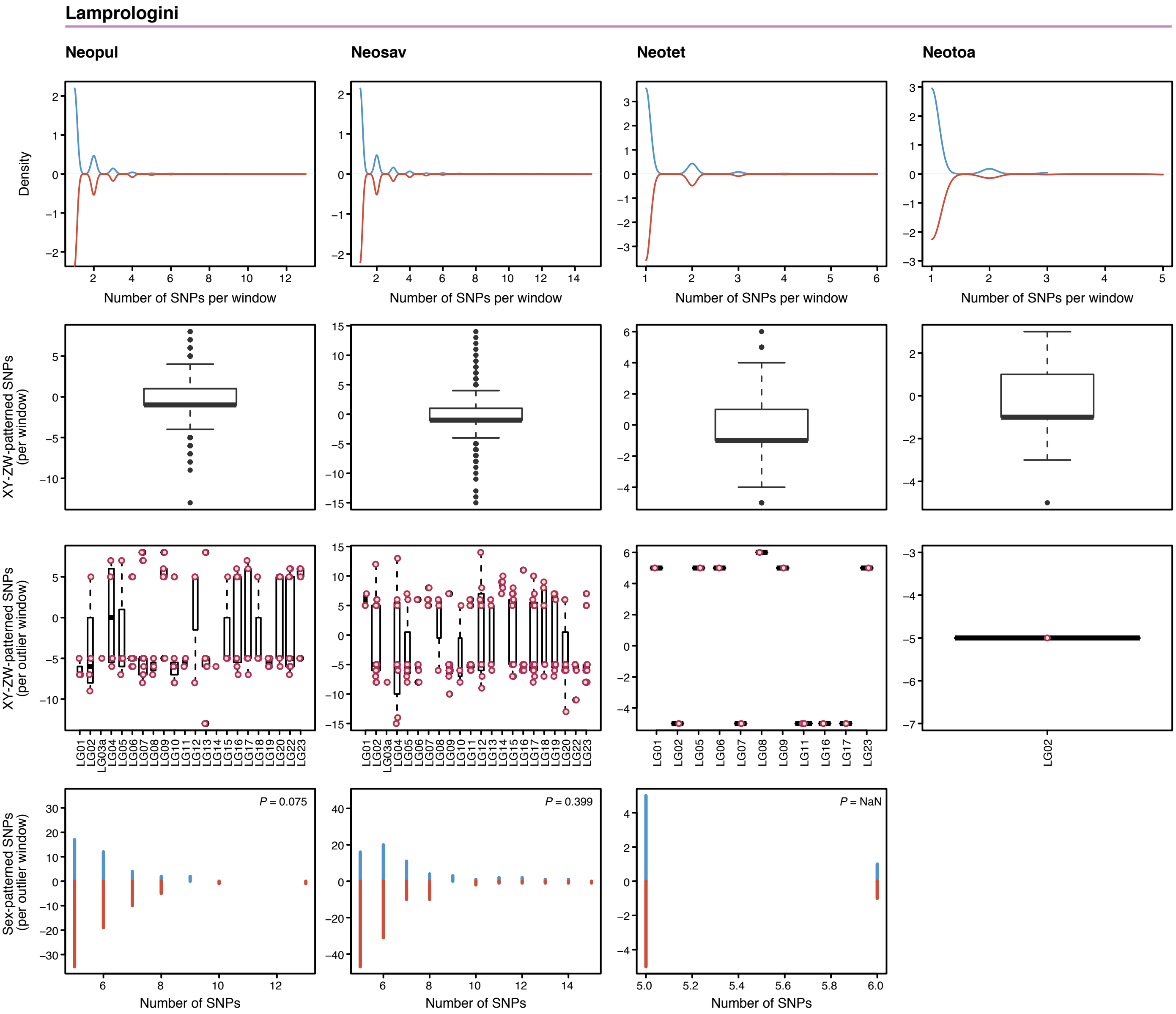

#### Supplementary Figure 6 (continued)

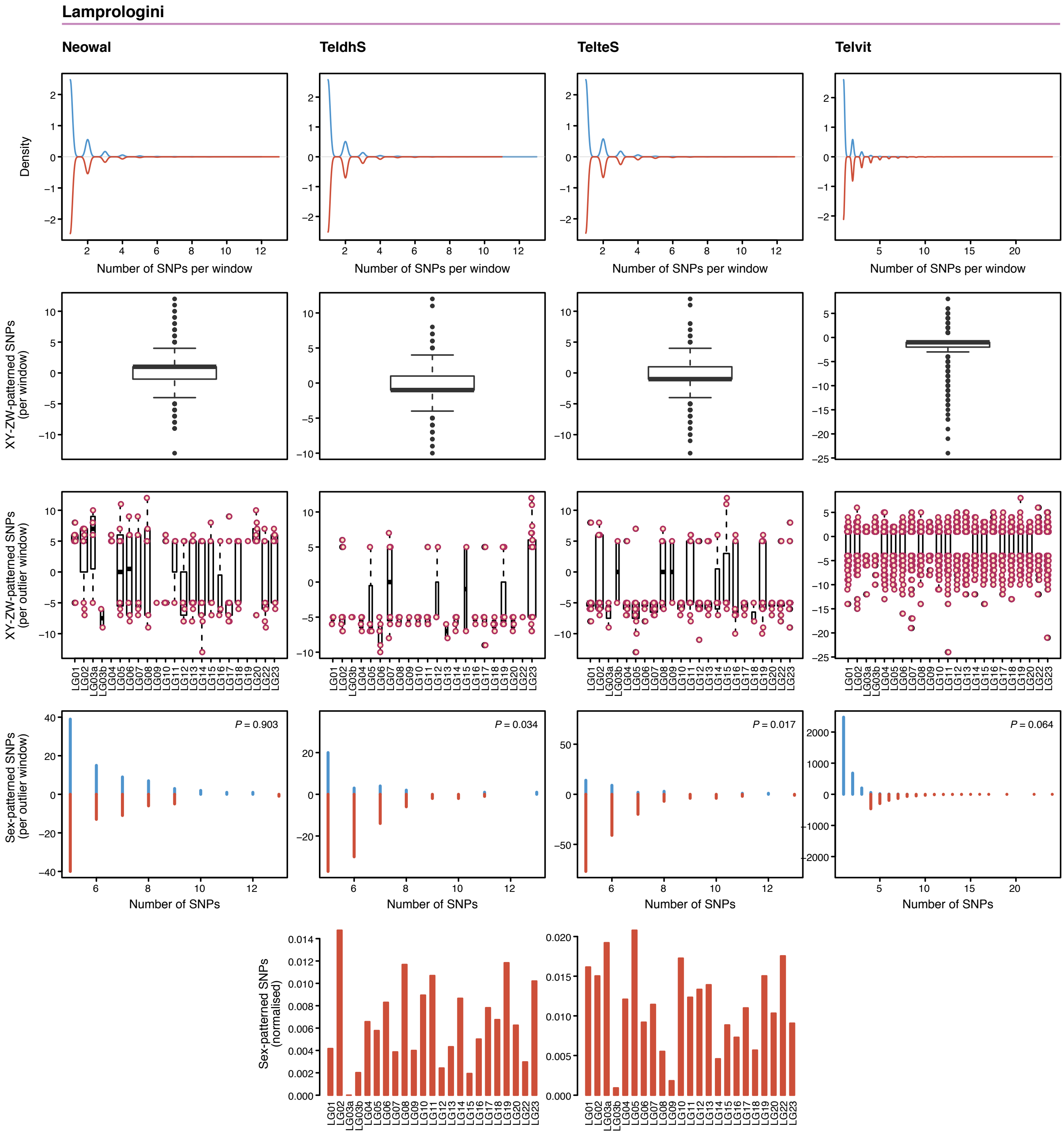

Supplementary Figure 6 (continued)

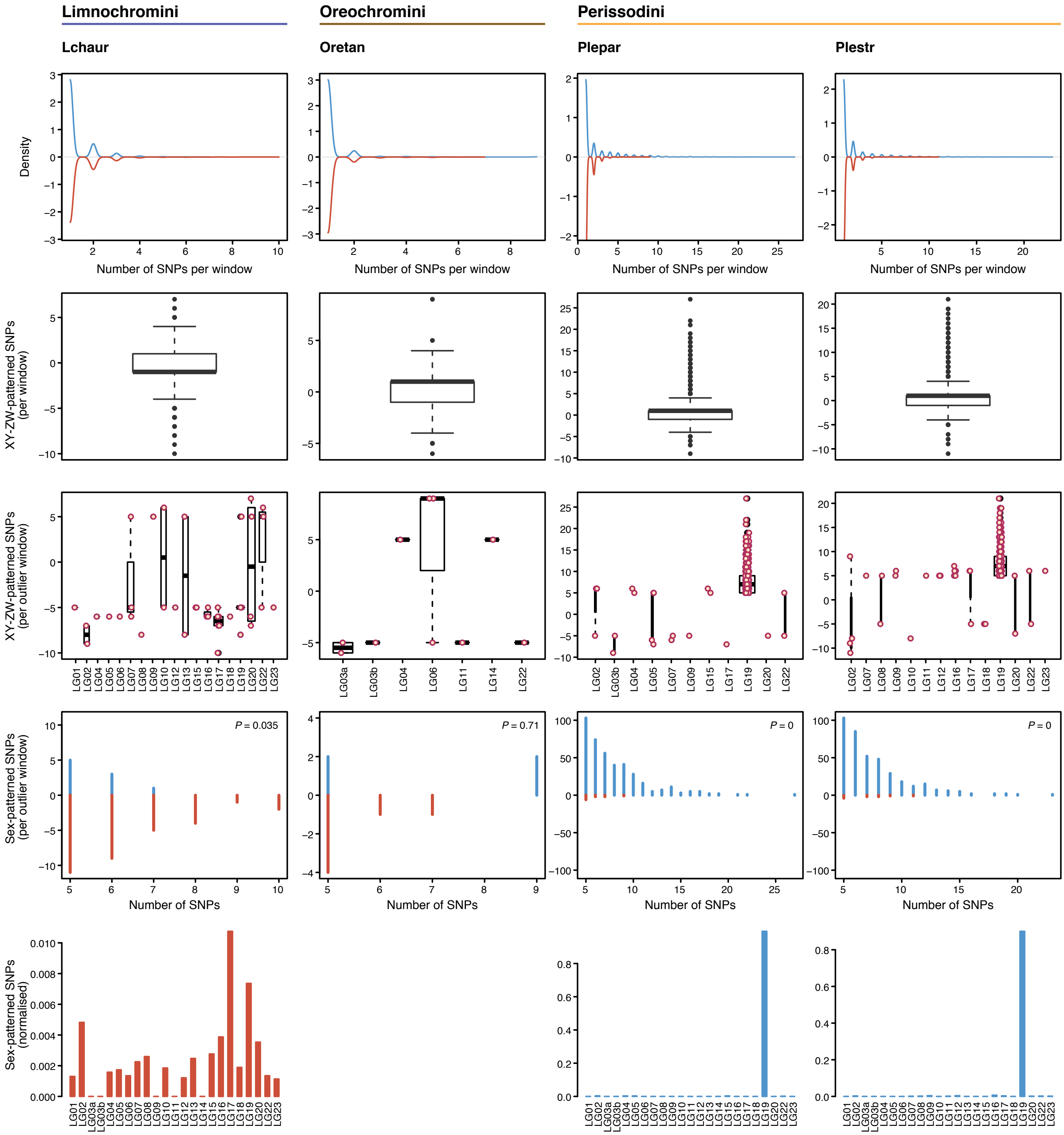

**Supplementary Figure 6 (continued)**

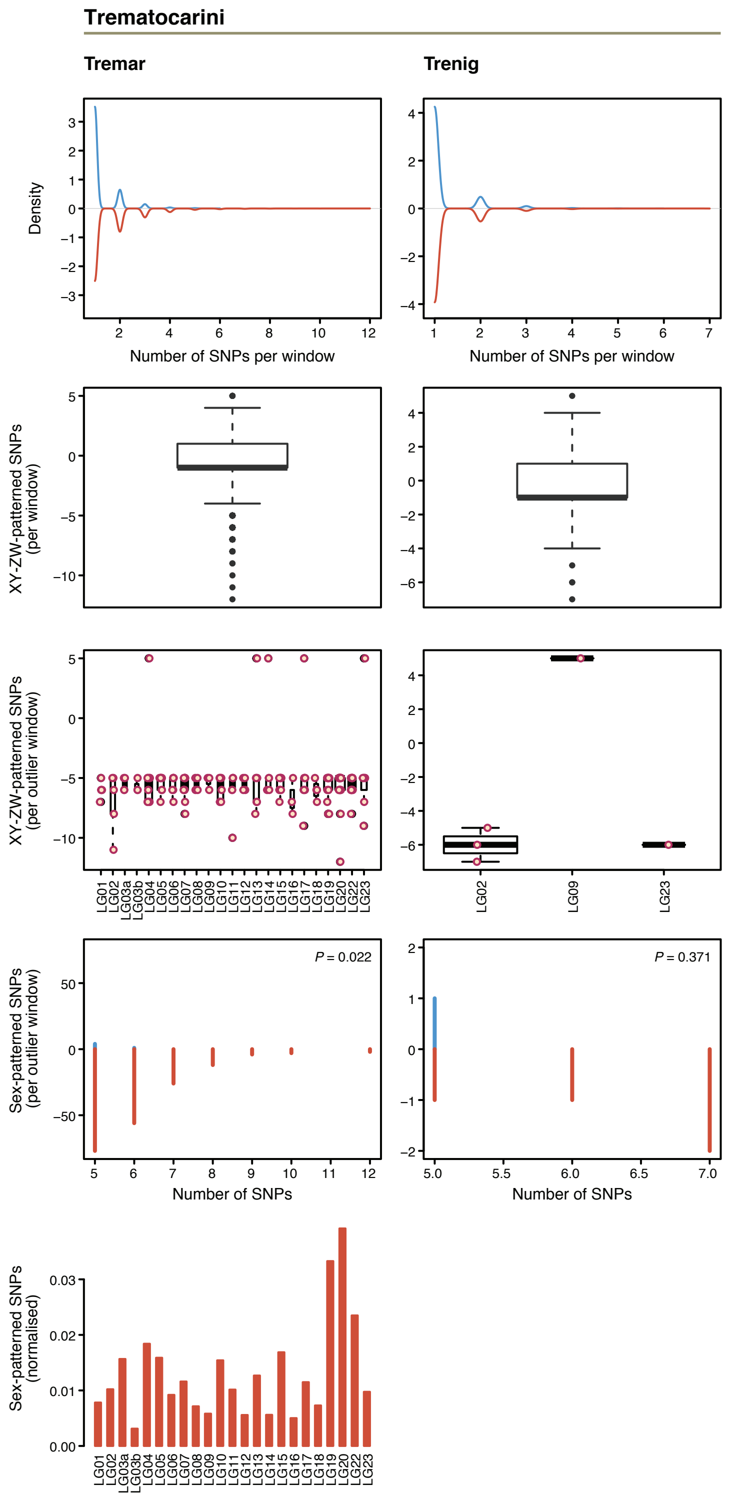


**Fig. S6. Identification of sex chromosomes based on transcriptome data (approach 3)**. Per-species visualization of the results for tests of allele differences based on replicate male and female transcriptome data (approach 3). For each species, four to five plots are shown i) Density plot of the number of sex-specific SNPs per 10 kb non-overlapping window; the density of XY- windows is represented in blue (positive y-axis scale) and the density of ZW-windows in red (negative y-axis scale); ii) Boxplot of the difference between XY- and ZW-SNPs in 10 kb windows. Windows with a higher number of XY-SNPs are represented with positive values (>0) while windows with a higher number of ZW-SNPs are represented with negative values (<0). Boxplot center lines represent the median; box limits the upper and lower quartiles and whiskers the 1.5x interquartile ranges. The width of the boxes is proportional to the number of observations; iii) Boxplot of the difference in XY- and ZW-SNPs for outlier windows only and represented per LG. Boxplot center lines represent the median; box limits the upper and lower quartiles and whiskers the 1.5x interquartile ranges. The width of the boxes is proportional to the number of observations. vi) Histogram of the number of sex-specific SNPs per 10 kb outlier window, XY-windows are represented in blue (positive y-axis scale) and ZW-windows in red (negative y-axis scale). For species with a detected heterogametic system v) Visualization of the number of sex-specific (XY-system in blue and ZW-system in red) SNPs of outlier windows normalized by chromosome length with species-specific Y-axis scales.

**Supplementary Figure 7**

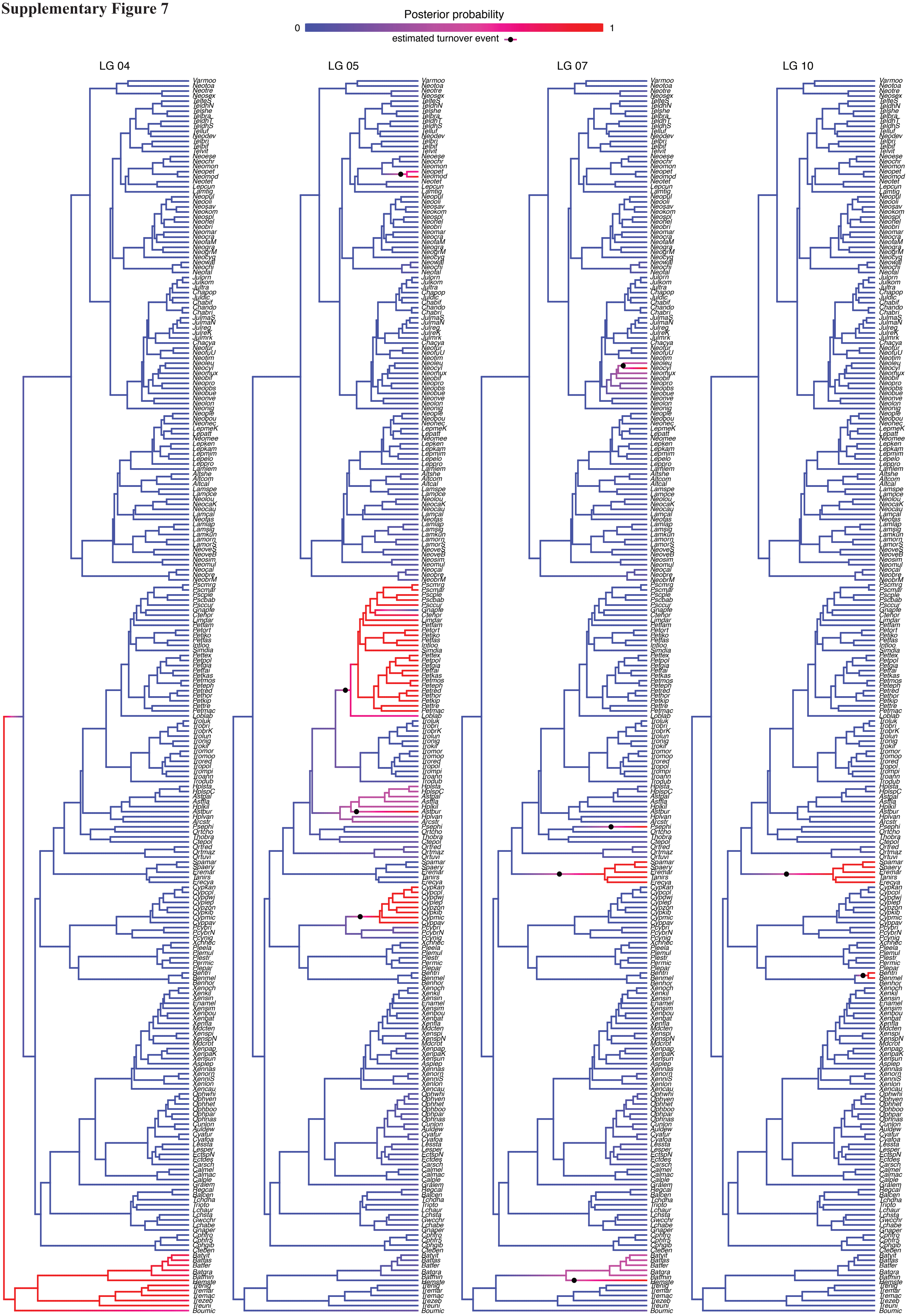

**Supplementary Figure 7 (continued)**

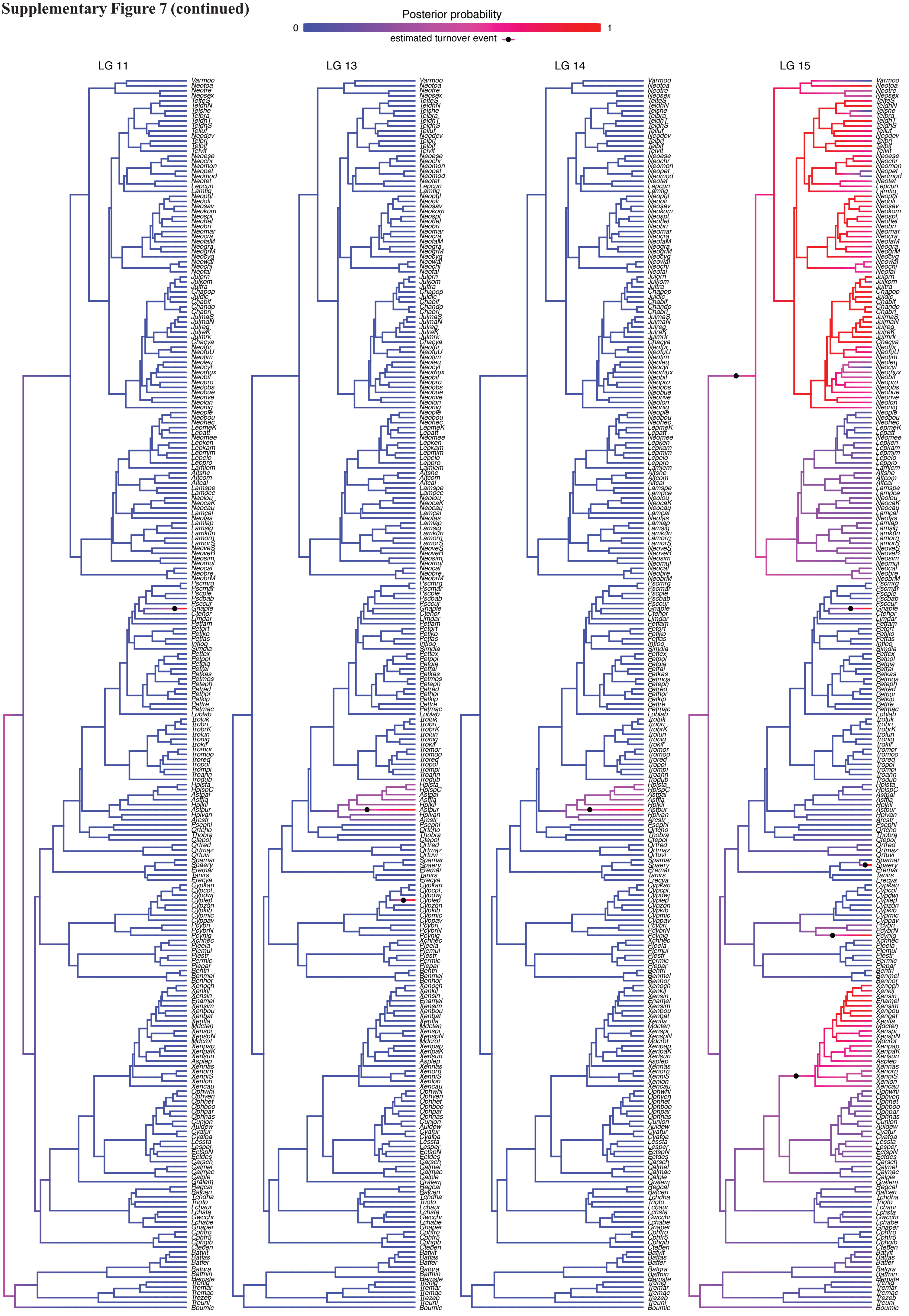

**Supplementary Figure 7 (continued)**

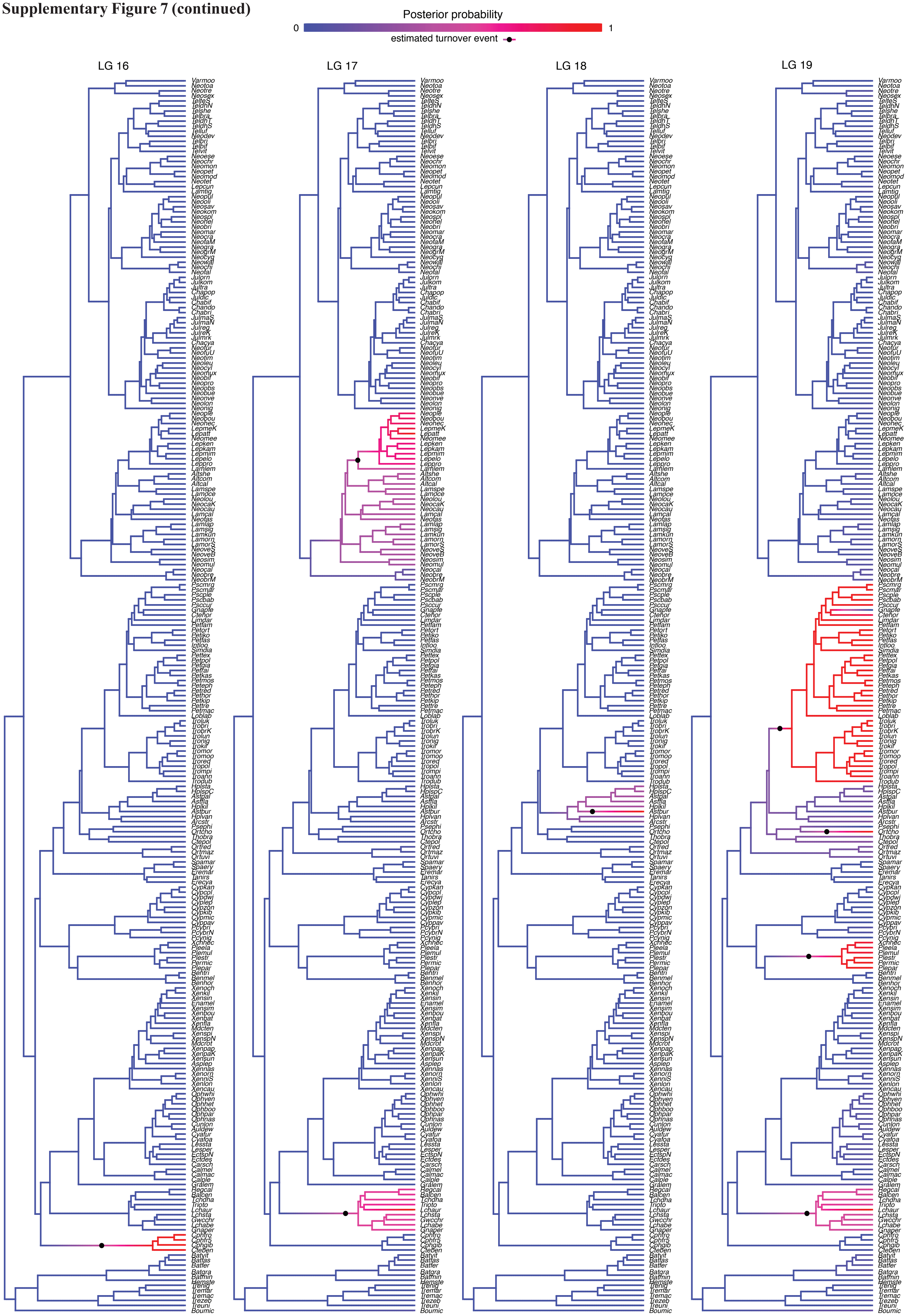

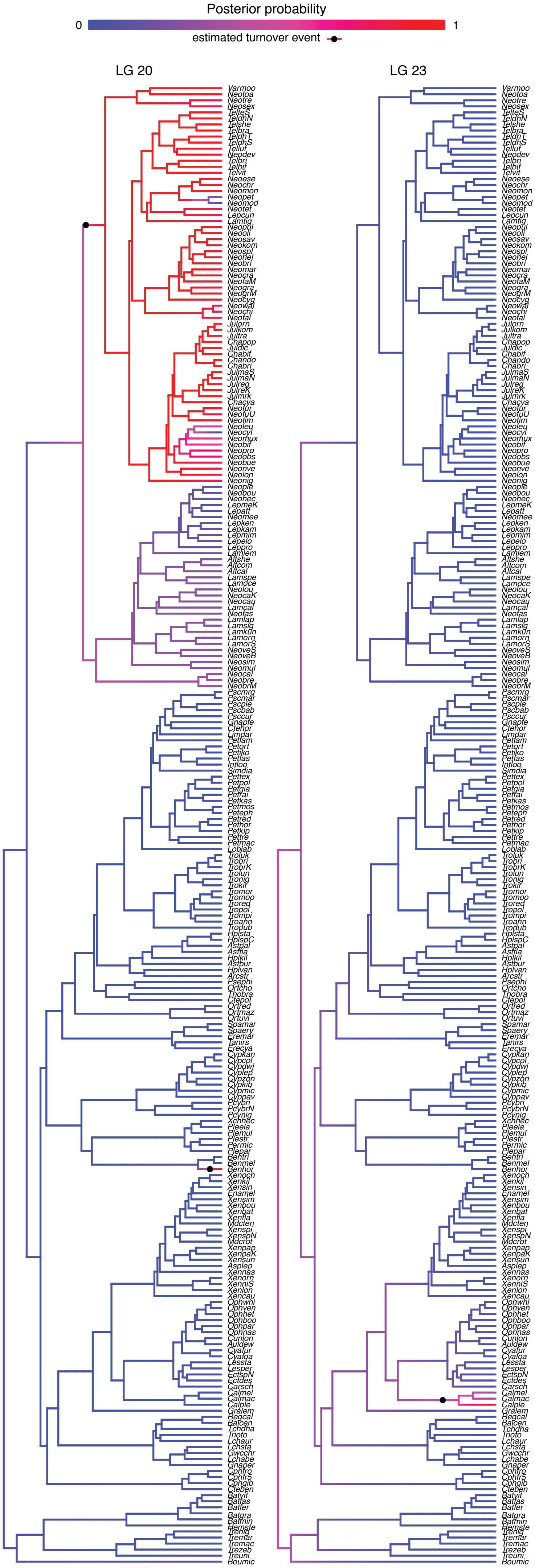


**Fig. S7. Sex chromosome turnover points.** Sex chromosome turnover points visualized per sex-linked LG derived from ancestral state reconstructions based on stochastic character mappings (using the permissive dataset).

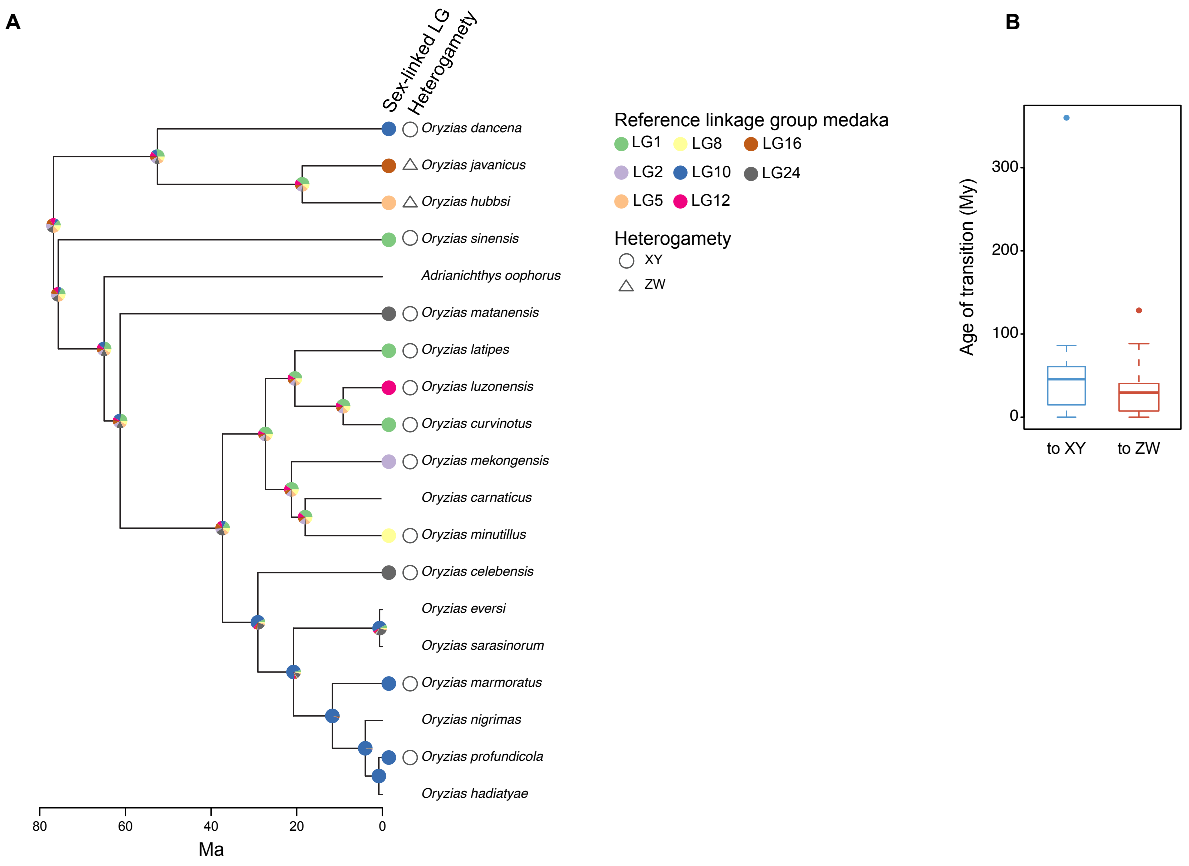


**Fig. S8. Sex chromosome evolution in ricefishes.** (**A**) Stochastic character mapping for sex chromosomes in ricefishes. Colored circles at tips represent sex-associated LGs with respect to the medaka (*Oryzias latipes*) genome. Pie charts at nodes represent the probability for an LG being a sex chromosome at this time. Heterogametic status is indicated at tips (triangle: ZW, circle: XY). (**B**) Age of heterogametic transitions in ray-finned fishes. Transitions to ZW-systems are younger than transitions to XY-systems (Kruskal-Wallis rank sum test *P* = 0.014). Boxplot center lines represent the median, box limits the upper and lower quartiles, and whiskers the 1.5x interquartile rang, points represent outliers. The size of boxes within a plot is proportional to the number of transition events.

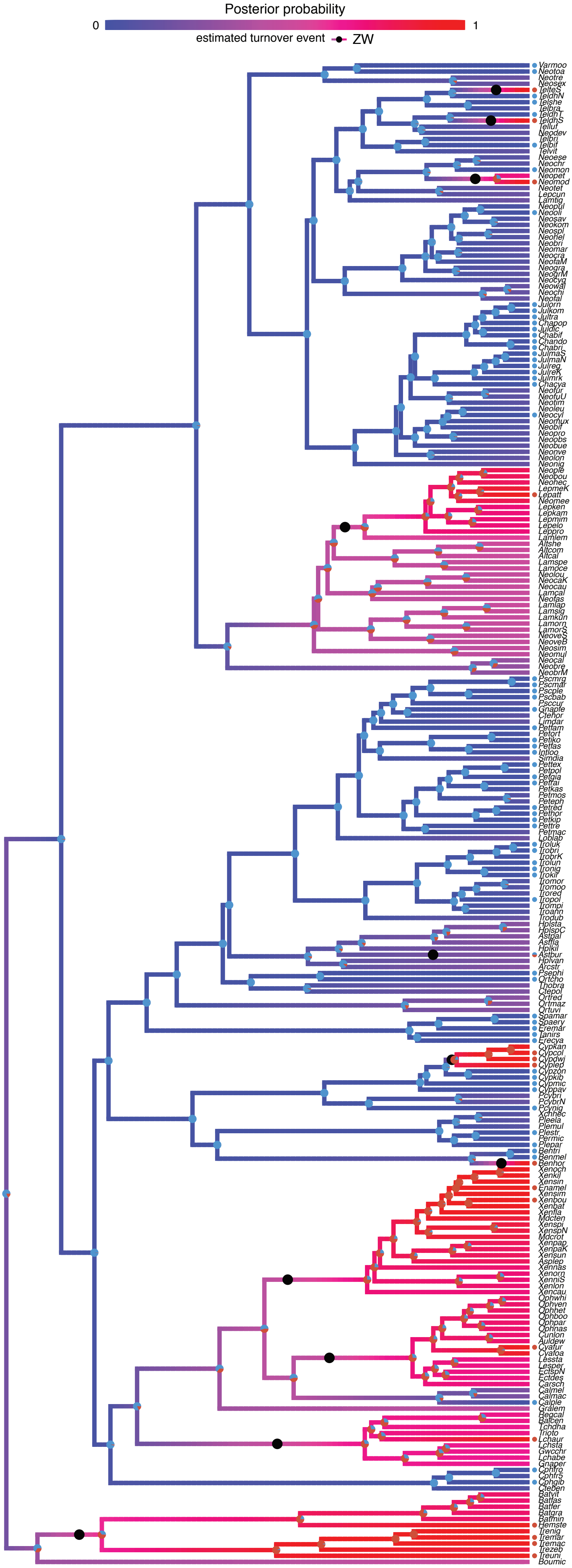


**Fig. S9. Stochastic character mapping for heterogametic status and inference of heterogametic changes.** Circles at the tips of the tree represent the identified heterogametic status (blue: XY, red: ZW). For species without a symbol, no sex chromosome could be identified. Pie charts at internal nodes represent the reconstructed heterogametic state in 1,000 simulations. Black points represent estimated heterogamety turnover points derived from ancestral state reconstructions based on stochastic character mappings.

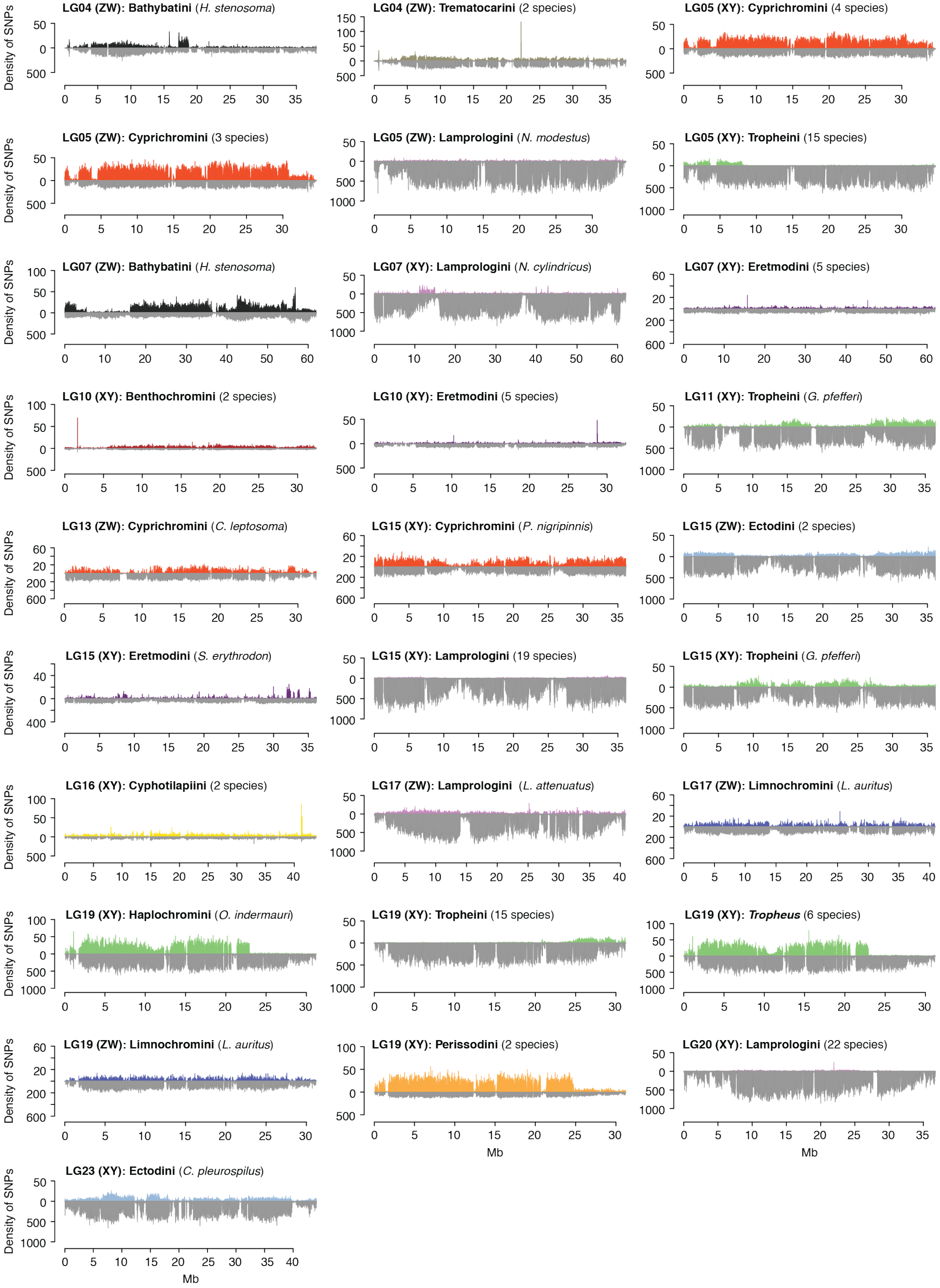


**Fig. S10. Distribution of sex-differentiated SNPs along the sex-associated LGs.** Each plot represents one identified sex chromosomal system. In each plot, upper panels show the density of XY- or ZW-SNPs, lower panels show the distribution of the total number of SNPs called for each window along the LG. Colored bars represent the number of sex-linked SNPs along the linkage group with color-coding according to tribe.

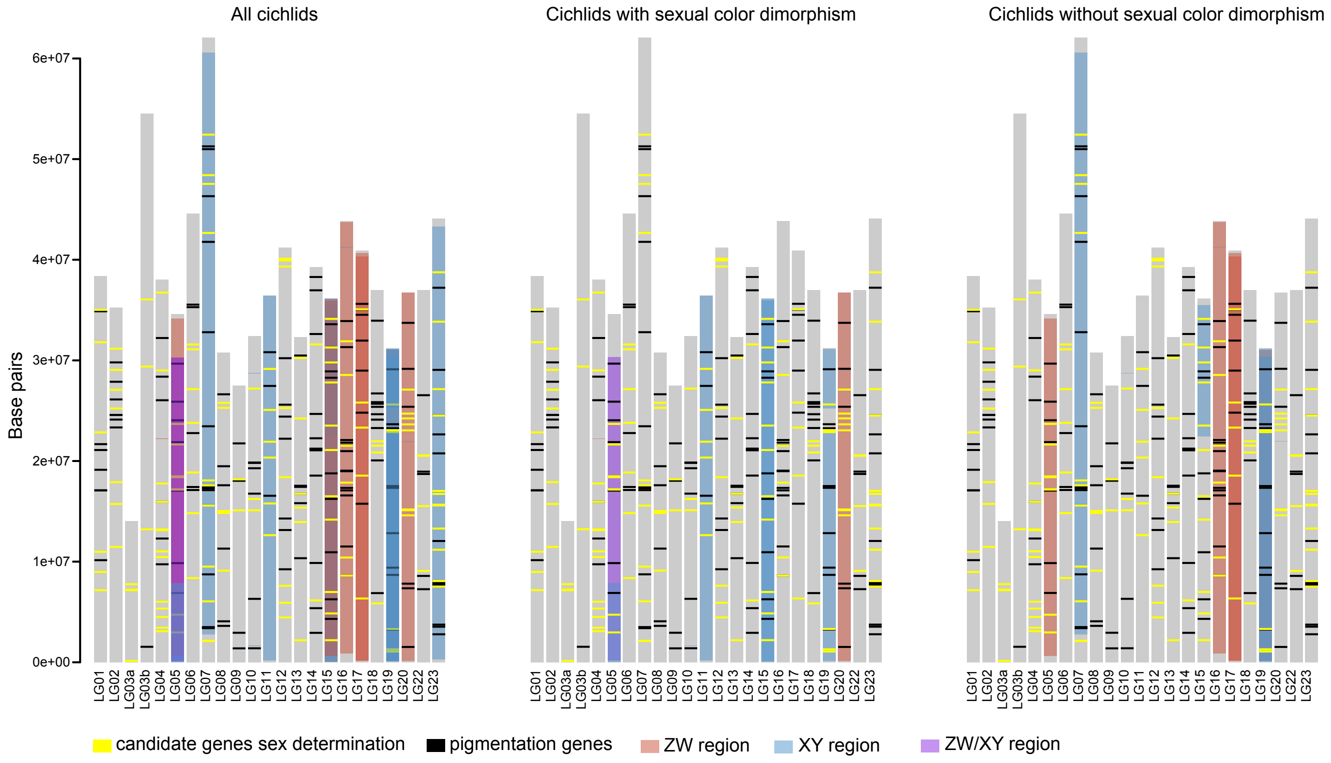


**Fig. S11. Distribution of candidate genes for sex determination and pigmentation.** Gray bars represent LGs and lines candidate genes for sex determination and pigmentation. Sex-differentiated regions are indicated with colored shadings.

**Supplementary Tables**

All supplementary tables are provided as excel file (Supplementary Material 2),

**Table S1** contains information on the included species, the experimental set-up and sample sizes for all analyses.

**Table S2** contains all obtained results concerning sex chromosome identification on the tribe- and subsequent species-level, including degree of sex chromosome differentiation, potential candidate genes in the sex-differentiated region, and source data for  fig. S11.

**Table S3** includes sequencing information for the used transcriptome data.

**Table S4** includes information on sex chromosomal systems in ricefishes (source data for fig. S8A).

**Table S5** includes information on sex chromosomal systems in ray-finned fishes (source data for fig. S8B).

**Table S6** contains reference genome gene IDs (NCBI) for putative candidate genes involved in sex determination and pigmentation (source data for fig. S11).
